# Structural basis for alpha-actinin inhibition of cofilin binding to actin filaments

**DOI:** 10.1101/2025.11.16.688667

**Authors:** Kien Xuan Ngo, Takashi Sumikama, Taro QP Uyeda, Ngan Thi Phuong Le, Han Gia Nguyen, Rémi Vuillemot, Sergei Grudinin

## Abstract

Actin is a conserved cytoskeletal protein essential for morphogenesis, motility, and division. Its versatility arises from filament assembly and regulation by actin binding proteins. Among these, alpha-actinin organizes filaments into bipolar or unipolar networks, whereas cofilin binds preferentially to ADP-actin regions and forms clusters to shorten the half helical pitch (HHP). Here, we investigated the molecular mechanism of how alpha-actinin alters filament and protomer conformations and influences cofilin binding. Using high-speed atomic force microscopy, all-atom molecular dynamics simulations, and principal component analysis, we show that alpha-actinin crosslinking stabilizes actin filaments in the canonical helical state, thereby preventing the cofilin-induced helical shortening required for cooperative filament decoration. Stabilization occurs without significant changes in protomer twist and rise and subdomain geometry and maintains a flattened protomer conformation that restricts twisting needed for cofilin binding. Furthermore, myosin II (S1) and cofilin preferentially bind to normal and shortened half-helical structures, respectively. In contrast, the isolated alpha-actinin-1 actin binding domain mutant (ABD-E235K) and wild-type (ABD-WT) induce local transitions from the canonical double-helical filament to the single-helical protofilament, parallel-helical protofilament state (PPS), and nano-ring structures, while only weakly affecting cofilin binding and cluster formation on PPS. Notably, cofilin can bind and form clusters along single protofilaments without inducing helical-pitch shortening. Together, these findings reveal a mechanistically distinct function of full-length alpha-actinin and its isolated ABD and support a stepwise mechanism in which cofilin cooperative binding to canonical double-helical actin filaments requires initial binding to actin regions with shortened HHP, followed by protomer twisting and further helical shortening.

## INTRODUCTION

Actin is one of the most highly conserved eukaryotic proteins; when assembled into filaments, actin plays a crucial role in various cellular processes, including morphogenesis, motility, division, intracellular transport, and membrane deformation in the cytosol (Pollard and Cooper, 2009). Beyond its cytosolic functions, actin filaments also participate in nuclear activities such as transcription regulation, chromatin remodeling, and DNA repair (Fernandez et al., 2024). Understanding the molecular mechanisms that drive the functional specialization of actin filaments within cells remains a central challenge in cell biology and biophysics.

Actin filaments are right-handed double-helices, and their subunits are arranged in a way that generates two distinct pitches: a long pitch and a short pitch. The long pitch represents the overall helical turn, hereafter referred to as a half helical pitch (HHP) of the filament and spans approximately 36 nm, corresponding to 13 subunits (6.5 protomer pairs) in the canonical actin filament (Egelman, 1994; Fujii et al., 2010; Holmes et al., 1990; Lorenz et al., 1993; Oda et al., 2009; Uyeda et al., 2025). The short 59 Å pitch-helix, on the other hand, refers to the left-handed genetic helix (Holmes et al., 1990), localized helical twist between adjacent actin protomers in the same protofilament. In canonical actin filaments, the twist between protomers *n* and *n* + 2 in the same strand is approximately 26° (equivalent to −167° between *n* and *n* + 1 in two opposite strands), whereas cofilactin filaments exhibit a larger twist of approximately 36° between *n* and *n* + 2 (equivalent to −162° between *n* and *n* + 1) (Galkin et al., 2011). The axial distance (AD) between two adjacent protomers (*n* and *n* + 2) within the same long-pitch strand is approximately 5.5 nm (Fujii et al., 2010; Holmes et al., 1990; Lorenz et al., 1993; Oda et al., 2009) and rise (a distance between n and n + 1 subunits) is approximately 2.75 nm (Reynolds et al., 2022). These structural features are fundamental to the establishment of the mechanical properties of the actin filaments (ben-Avraham and Tirion, 1995; Tsuda et al., 1996), influencing their stability (Okura et al., 2023; Sun and Murrell, 2024).

Alpha-actinin, a major actin crosslinker, plays an essential role in cytoskeletal organization and cellular mechanics (Carol and Carpen, 2004; Sjöblom et al., 2008). Alpha-actinin crosslinks actin filaments at every half-helical turn, leaving numerous bare protomers between crosslinking sites accessible for cofilin and myosin II binding. It organizes actin filaments into bipolar bundles within contractile rings, stress fibers, and Z-disc, and into unipolar arrays in sarcomeres, lamellipodia, cortical actin, and filopodia, which together support diverse and essential cellular functions (Begg et al., 1978; Hampton et al., 2007; Rajan et al., 2023; Taylor et al., 2000; Tojkander et al., 2012; Xue et al., 2010) (**Table 1**).

**Table 1.** A brief comparison of structure, localization, and functions of actin filaments crosslinked by alpha-actinin in bipolar and unipolar configurations within cells (Begg et al., 1978; Hampton et al., 2007; Rajan et al., 2023; Tatyana M. Svitkina, 2020; Taylor et al., 2000; Tojkander et al., 2012).

| **Feature** | **Bipolar**<br>**configuration** | **Unipolar**<br>**configuration** |
|---|---|---|
| Polarity | The barbed and pointed ends are oriented in opposite directions. | The barbed and pointed ends are oriented in the same directions. |
| Crosslinking by alpha-actinin | Forms contractile networks | Forms stable, non-contractile bundles |
| Myosin interaction | Strong<br>(contractile activity) | Weak<br>(non-contractile activity) |
| Cellular function | Force generation, contraction | Protrusion, structural support, migration |
| Localization in cells | Stress fibers, contractile rings, Z-disc | Sarcomeres*, lamellipodia, filopodia, cortical actin, microvilli |

\* Strong interaction with myosin II, force generation and contraction

Recently, we applied an integrated SimHS-AFMfit–MD approach to investigate the dynamic atomic conformations of Ca²⁺-bound and Ca²⁺-free alpha-actinin-2 in the absence of actin filaments, revealing a closed conformation of the tandem calponin homology (CH1-CH2) domains within the actin binding domain (ABD) (Ngo et al., 2026). However, extending this approach to complex alpha-actinin-crosslinked actin filaments remains challenging because of molecular crowding and overlapping features, which hinder reliable image segmentation and computer vision-based structural recognition. In humans, muscle-specific alpha-actinin isoforms 2 and 3 are Ca²⁺ insensitive and regulated by phosphoinositides (Fraley et al., 2003; Fukami et al., 1992; Liu et al., 2004; Ribeiro et al., 2014; Taylor and Taylor, 1994; Young, 2000). In contrast, non-muscle alpha-actinin isoforms 1 and 4 are Ca^2+^-sensitive (Foley and Young, 2014; Pinotsis et al., 2020; Tang et al., 2001). Structural studies have resolved the atomic architecture of these isoforms (Foley and Young, 2014; Pinotsis et al., 2020; Tang et al., 2001), yet their impact on actin filament conformational dynamics at molecular and atomic levels remains unclear.

Cofilin binds actin filaments at a 1:1 molar ratio, forming cofilactin segments that induce supertwisting and reduce the HHP by ∼25%, decreasing protomers to ∼11 per half helix (Galkin et al., 2011; McGough et al., 1997). Cryo-EM revealed that internal and barbed end subunits are flattened, while pointed end subunits are more twisted, resembling G-actin monomers (Carman et al., 2023). Other studies have suggested that twisted subunits (P) at the pointed end, where the D-loop of the penultimate protomer P1 interacts with terminal subunit P, inhibit subunit addition to the pointed end and thereby slow filament elongation (Carman et al., 2023; Narita et al., 2011; Zsolnay et al., 2020). A recent study suggests that cofilin reverts terminal subunits at both the pointed and barbed ends to a G-actin-like conformation while undertwisting the filament short-pitch helix (Palmer et al., 2026). Our previous work showed that shortened bare HHPs near cofilin clusters on the pointed-end side contained fewer protomers and displayed larger mean AD (MAD; 5.0-6.3 nm) than normal HHPs (MAD: 4.3-5.6 nm). However, it remains unresolved whether structural transitions within these shortened bare HHPs, specifically a change from a flattened F-actin to a twisted C-actin–like conformation in actin protomers, are prerequisites for driving shortening helices and cooperative cofilin binding and cluster formation (Ngo et al., 2024). Clarifying these structural features will be essential for understanding actin conformation-dependent cooperative cofilin binding and cluster formation, as well as the polymorphic structures of actin filaments in regulating the spatial and temporal distribution of other actin binding proteins (ABPs) such as myosin II in cells.

Myosin II is a non-processive actin-based motor that assembles into bipolar thick filaments, enabling simultaneous binding and crosslinking of multiple actin filaments to generate contractile forces. Through ATP-dependent mechanochemical cycles, the myosin II subfragment 1 (S1) transitions between weakly and strongly actin-bound states, coupling ATP hydrolysis to lever-arm conformational changes that convert chemical energy into mechanical work. Although previous structural studies of the actomyosin II complex have extensively characterized the conformational states of myosin II, they consistently reported that the overall conformation of actin filaments remains largely unchanged (Behrmann et al., 2012; Mornet et al., 1981; Rayment et al., 1993; Von Der Ecken et al., 2016). Remarkably, a recent study reported that myosin V and VI, which move toward the opposite ends of actin filament, generate forces that remodel the double-helical lattice into supercoiled conformations that are selectively recognized and stabilized by α-catenin, revealing a structural mechanism by which actin mechanics regulate actin binding protein interactions and force-dependent mechanotransduction (Carl et al., 2026).

Here, we combine high-speed atomic force microscopy (HS-AFM), all-atom molecular dynamics (MD) simulations, and principal component analysis (PCA) to investigate how alpha-actinin crosslinking modulates actin filament architecture and dynamics. We focus on (i) the conformational behavior of alpha-actinin in unipolar and bipolar crosslinked filaments; (ii) its crosslinking effects on filament helical twist, rise and AD between protomers, dihedral angles (rotational orientations between inner domain (ID: consisted of subdomains (SD1-SD2) and outer domain (OD: composed of (SD3-SD4), see Methods for classification of actin domains), thermodynamic conformational landscapes within an actin subunit; (iii) binding effect of alpha-actinin-1 actin binding domain mutant and wild-type (ABD-E235K, ABD-WT) on helical feature of actin filament, and (iv) the consequences for cofilin binding and cluster formation.

Our HS-AFM results reveal that high crosslinking densities of alpha-actinin preserve canonical HHP and MAD features in actin filaments more effectively than low alpha-actinin crosslinking density and bare actin filaments in the presence of cofilin, thereby better suppressing cofilin cluster formation. The flattened protomer conformations stabilized by alpha-actinin crosslinking suppress cofilin cluster formation while likely leaving myosin II motor (S1) interactions with the filaments unaffected. In contrast, bare actin filaments exhibit the expected cofilin-induced helical shortening along with cooperative cofilin binding and clustering. Additionally, cofilin and S1 interact with distinct regions of actin filament, cofilin associating with the shortened HHPs and S1 with the normal HHPs, enabling both to stably bind and cluster along the same filament. In addition, the isolated alpha-actinin 1 ABD-E235K and ABD-WT comprising tandem calponin homology (CH1-CH2) domains, transiently binds to actin filaments and induces local transitions from the canonical double-helical filament to the single-helical protofilament (SP), parallel-helical protofilament state (PPS), and nano-ring structure, suggesting that alpha-actinin and its ABD likely regulate actin structures and functions through two distinct mechanisms. The cofilin cluster formation on PPS is largely unaffected, exploring new properties that cofilin could form clusters on single protofilaments without changing filament helical length. MD simulation and PCA results further complement that alpha-actinin crosslinking stabilizes fluctuation in canonical double-helical actin filament twist, rise, AD between adjacent protomers, as well as the dihedral angles and minimum free energy landscape within protomers in the flattened state like F-actin conformation.

## RESULTS

### Analysis of conformational dynamics of alpha-actinin crosslinked actin filaments using HS-AFM

The question of how actin filaments adopt different conformations to regulate the functions and spatial and temporal distributions of various ABPs within cells remains unclear (Uyeda et al., 2025). In this study, we combined HS-AFM, MD simulations, and PCA to investigate the surface topography, atomic-level structural changes, and dynamic behavior of actin filaments crosslinked by alpha-actinin interacting with S1 or cofilin (**Figure S1**).

Generally, HS-AFM images capture three-dimensional (3D) surface structure, incorporating two-dimensional (2D) lateral (x-y) and one-dimensional (1D) height (z) information encoded in pixel intensity (Ando et al., 2024). Here, we employed HS-AFM to analyze the changes in the HHP, the number of protomer pairs per HHP, and the MAD in filaments with a unipolar crosslinking configuration (**Figure 1, Figure S2-S3, Video S1**), as similarly done (Ngo et al., 2024). The unipolar crosslinking configuration, in which the actin filaments shared the same polarity was identified by examining the persistent binding and tilting of S1 to the actin filaments (**Figure S3B, Video S1**); the persistent binding of S1 occurred when ATP was gradually depleted and was explained in captions **Figure S3B** and references (Begg et al., 1978; Ngo et al., 2015). These parameters fluctuated over time (**Figure 1C, Figure S3C**), indicating that HHP, the number of actin protomer pairs per HHP, and the MAD values of actin filaments in the unipolar crosslinking configuration in the presence of ATP and S1 were not static but rather dynamic. Consequently, these parameters were characterized by broad distributions: HHP (36.3 ± 2.3 nm, N = 1055); number of protomer pairs per HHP (5.5 pairs/HHP [5.4%], 6.5 pairs/HHP [80.3%], 7.5 pairs/HHP [13.8%], and 8.5 pairs/HHP [0.5%]); and MAD (5.5 ± 0.3 nm, N = 1055) (**Figure 1D**).

**Figure 1.**
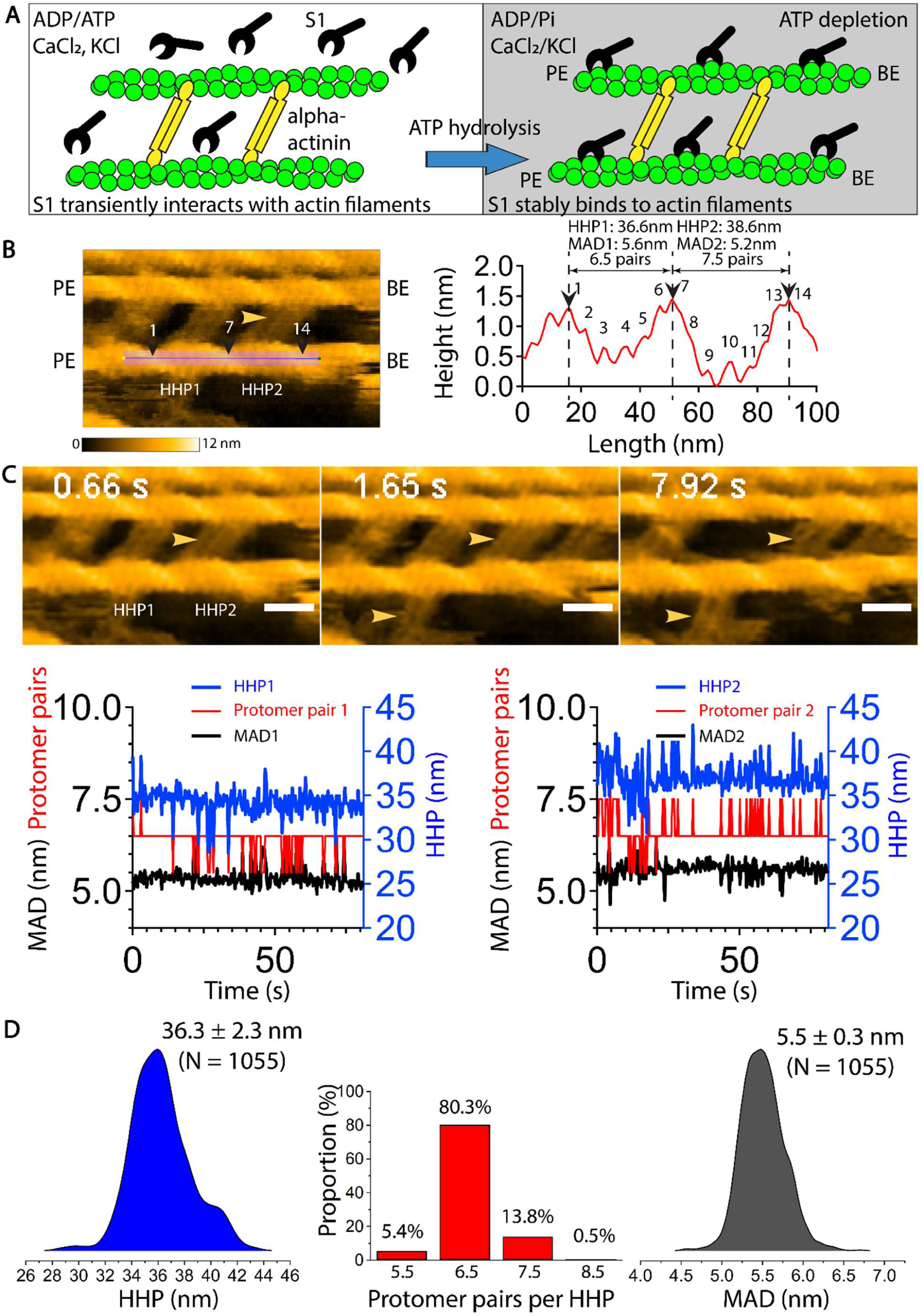
HS-AFM-based analysis of the dynamic conformation of the actin filaments crosslinked with alpha-actinin in a unipolar configuration in the presence of ATP and S1. **(A)** Illustration of the experimental setup for HS-AFM observations of dynamic conformational changes in alpha-actinin-crosslinked actin filaments in the presence of S1, ATP, ADP, CaCl_2_, and KCl at pH 6.8. **(B)** Longitudinal section profile illustrating the analysis of the magnitudes of the two adjacent HHPs, the number of protomer pairs per HHP, and the mean axial distance (MAD) between two adjacent protomers within the same-long pitch strand of actin filament. The height reference was defined along the actin filament. The semi-automatic analytic methods employed were developed previously (Ngo et al., 2024). Yellow arrowheads denote alpha-actinin; black arrowheads represent the filament crossover points used to measure HHP. The polarity of actin filaments in this unipolar model was confirmed by observing the tilted angles of S1, as shown in **Figure S3B**. PE and BE denote pointed end and barbed end, respectively. Still AFM images were similarly reused in Figure S2, S3, S4B. **(C)** Time-dependent changes in the magnitudes of two adjacent HHPs, the number of protomer pairs per HHP, and the MADs in the actin filaments crosslinked with alpha-actinin in the presence of S1. Actin filaments crosslinked by 50 nM alpha-actinin in the presence of 1 mM ATP, 0.1 mM CaCl₂, and 1 µM S1, but without MgCl_2_. Note that two alpha-actinin homodimers may cooperate to crosslink every half-helix of actin filaments, with their orientations varying due to molecular rotation. Color scale: 0–12 nm. Yellow arrowheads denote alpha-actinin. **(D)** Ridgeline and bar plots displaying the distribution of the magnitude of the HHPs, the number of the protomer pairs per HHP, and the MADs, as well as the corresponding number of events (N) (1055 and 1055 HHPs, respectively). The values are mean ± SD and analyzed from 5 filaments.

Next, we analyzed and compared HHP, crosslinking angles of alpha-actinin, and crosslinking lengths between alpha-actinin-crosslinked filaments with and without S1 (**Figure S4, Table 1**). In the presence of ADP and ATP but not S1, the mean HHP in the filaments crosslinked by alpha-actinin was 36.5 ± 2.8 nm (N = 4270) (**Figure S4A-C, Video S2**). These results were similar to those observed in the presence of both S1 and ATP (**Figure 1D, Figure S3, Video S1**). The crosslinking lengths and crosslinking angles showed a small difference between alpha-actinin-crosslinked filaments with and without S1 (**Figure S4D-E, Table 2**). These results indicated that the HHP of the actin filaments in unipolar crosslinking configurations underwent intrinsic thermal fluctuations around a mean value, accompanied by changes in the number of protomer pairs per HHP. We note that an assembly of two alpha-actinin homodimers can stably bind to and crosslink actin filaments (**Figure 1; Figure S3-S4**). Although the crosslinking angles of alpha-actinin and the crosslinking lengths fluctuate around their respective mean values, neither parameters cause a significant change in the HHP, regardless of the presence of S1. These findings suggest that alpha-actinin undergoes substantial conformational rearrangements while preserving the helical periodicity of actin filaments. Under ATP-nearly depleted conditions (**Figure S3B, Video S1**), S1 persistently associated with the actin filaments in these unipolar crosslinking configurations, suggesting that the ATP-dependent interactions of S1 with alpha-actinin-crosslinked actin filaments were largely unaffected and comparable to those with bare actin filaments.

**Table 2.** A summary of alpha-actinin crosslinking angles relative to the filament axis and crosslinking lengths between alpha-actinin-crosslinked actin filaments, as determined from HS-AFM experiments. The data are shown and mean ± standard deviation (SD).

| **Figure S4D**<br>**(HS-AFM)** | Mean ± SD | N |
|---|---|---|
| Crosslinking angle with S1 (°) | 61.2 ± 3.6 | 200 |
| Crosslinking angle without S1 (°) | 67.6 ± 4.5 | 124 |
| **Figure S4E**<br>**(HS-AFM)** |  |  |
| Crosslinking length with S1 (nm) | 39.5 ± 1.7 | 200 |
| Crosslinking length without S1 (nm) | 40.8 ± 1.1 | 124 |

To obtain the complimentary results of these observations, we analyzed crosslinking angles, crosslinking lengths, HHP, and AD in unipolar and bipolar models generated from MD-derived conformational libraries (**Figure S5, Table 3-4**). MD-derived unipolar crosslinking angles (67.9 ± 4.1°) closely matched HS-AFM measurements (61.2 ± 3.6° and 67.6 ± 4.5° with and without S1, respectively). Both crosslinking angles and lengths varied dynamically and differed between unipolar and bipolar assemblies (**Figure S5A,B, Video S13-S14**).

**Table 3.** A summary of alpha-actinin crosslinking angles relative to the filament axis and crosslinking lengths of actin filaments between alpha-actinin-crosslinked actin filaments, as determined from MD simulations. The data are shown as mean ± SD.

| **Figure S5B**<br>(MD simulations) | Crosslinking length (nm) | Crosslinking angle (°) |
|---|---|---|
| Bipolar-flexible | 39.0 ± 0.9<br>(N = 30000) | 88.2 ± 6.7<br>(N = 30000) |
| Bipolar-fixed | 20.2 ± 0.4<br>(N = 10000) | 42.9 ± 0.0<br>(N = 10000) |
| Unipolar-flexible | 30.8 ± 1.1<br>(N = 25000) | 67.9 ± 4.1<br>(N = 25000) |

Despite nearly identical mean AD values across all models (5.57–5.58 nm), bipolar crosslinking produced substantially greater AD fluctuations (0.07–0.11 nm) than bare actin and the unipolar model (both 0.03 nm). Notably, restraining alpha-actinin in the bipolar-fixed model did not suppress these fluctuations (**Figure S5C, Figure S6, Table 4**). In contrast, HHP remained nearly constant across all models (∼36.2–36.3 nm), consistent with the use of 13 fixed actin protomers per filament and previous findings that HHP variations primarily reflect changes in the number of protomers per repeat (Egelman et al., 1982; Fineberg et al., 2024; Ngo et al., 2024).

**Table 4.** Summary of HHP and AD (mean ± SD) values measured from bare actin filaments and actin filaments in bipolar-flexible, bipolar-fixed, and unipolar-flexible models conducted by MD simulations. The data are shown as mean ± SD. Related to **Figure S5C.**

| Model | HHP (nm) | AD (nm) |
|---|---|---|
| Bare actin | 36.25 ± 0.01<br>(N = 14) | 5.57 ± 0.03<br>(N = 34) |
| Bipolar-flexible | 36.26 ± 0.01<br>(N = 14) | 5.58 ± 0.07<br>(N = 59) |
| Bipolar-fixed | 36.21 ± 0.01<br>(N = 13) | 5.58 ± 0.11<br>(N = 78) |
| Unipolar-flexible | 36.20 ± 0.01<br>(N = 14) | 5.57 ± 0.03<br>(N = 37) |

Together, these results indicate that enhanced inter-subunit dynamics arise primarily from bipolar crosslinking geometry rather than alpha-actinin’s intrinsic thermal mobility.

### Alpha-actinin crosslinking stabilizes actin helical twist by suppressing cofilin-induced supertwisting and cluster formation

The HHP and MAD of alpha-actinin-crosslinked actin filaments obtained in this study (**Figure 1, Figure S3, Videos S1-S2**) were comparable with those of the control actin filaments (specifically, those without alpha-actinin crosslinking) investigated in our previous studies (e.g., HHP: 36.8 ± 4.3 nm, MAD: 4.3–5.6 nm) (Ngo et al., 2024, 2015). Here, we further investigated the physical mechanisms by which alpha-actinin stabilizes the helical twists of actin filaments, both in the presence and absence of S1 or cofilin. To do this, we used cofilin, as cofilin non-competitively excludes S1 from binding to actin filaments in the presence of ATP (Ngo et al., 2016), or competitively inhibits S1 binding to actin in the absence of physiological ATP (Wiggan et al., 2012).

To further explore the physical effects of alpha-actinin on the helical twists of the actin filaments, we initially induced crosslinking between the actin filaments by alpha-actinin in F-buffer at pH 6.8. Cofilin was then added at a final concentration of 600 nM; this concentration was significantly greater than its binding affinity (*K_d_*) of 5–10 nM for bare actin filaments (Andrianantoandro and Pollard, 2006), and the results were compared with those of the actin filaments with 600 nM cofilin but without the alpha-actin crosslinking (**Figure 2, Figure S7-S9, Table 5, Video S3-S5**). The kinetics of the peak height and HHP fluctuations were analyzed using the same approach as in our previous studies (Ngo et al., 2024, 2015) (**Figure 2A-C**). The mean peak height and HHP values of the actin segments crosslinked with alpha-actinin and bound with cofilin (referred as partial cofilin-actinin-actin) were 9.9 ± 1.1 nm (N = 2514) and 33.0 ± 4.6 nm (N = 2003), respectively. These clusters exhibited a significantly lower peak height and longer HHP than the full cofilactin clusters (peak height: 10.7 ± 1.0 nm, N = 425; HHP: 27.9 ± 3.0 nm, N = 300; *p* < 0.05) (**Figure 2D**, **Table 5**). However, these values were significantly (*p* < 0.05) greater than the peak height (8.6 ± 0.8 nm) and slightly shorter than the normal HHP (36.8 ± 4.3 nm) of the bare actin segments reported in our previous study (Ngo et al., 2015). These results suggest that although cofilin can bind to alpha-actinin-crosslinked actin filaments. Alpha-actinin crosslinking impedes the characteristic cofilin-induced shortening of the HHP, thereby delaying the formation of full cofilactin clusters. Further analysis revealed that cofilin binding and cluster formation were strongly inhibited in actin filaments with high alpha-actinin crosslinking densities but were weakly suppressed with low crosslinking densities (**Figure S9, Video S4B**). Two key findings emerged from these experiments: (i) High crosslinking densities of alpha-actinin significantly inhibited the super twisting of the half helices more effectively than bare filaments; (ii) High crosslinking densities of alpha-actinin strongly inhibited the cooperative binding of cofilin to form the full cofilin clusters. Together, these findings indicate that high alpha-actinin crosslinking densities stabilize the HHP of actin filaments more effectively than low crosslinking densities or bare filaments when cofilin proteins are present. Bare actin filaments show the expected cofilin-induced helical shortening, accompanied by cooperative cofilin binding and cluster formation (**Figure S7, Video S4A**).

**Figure 2.**
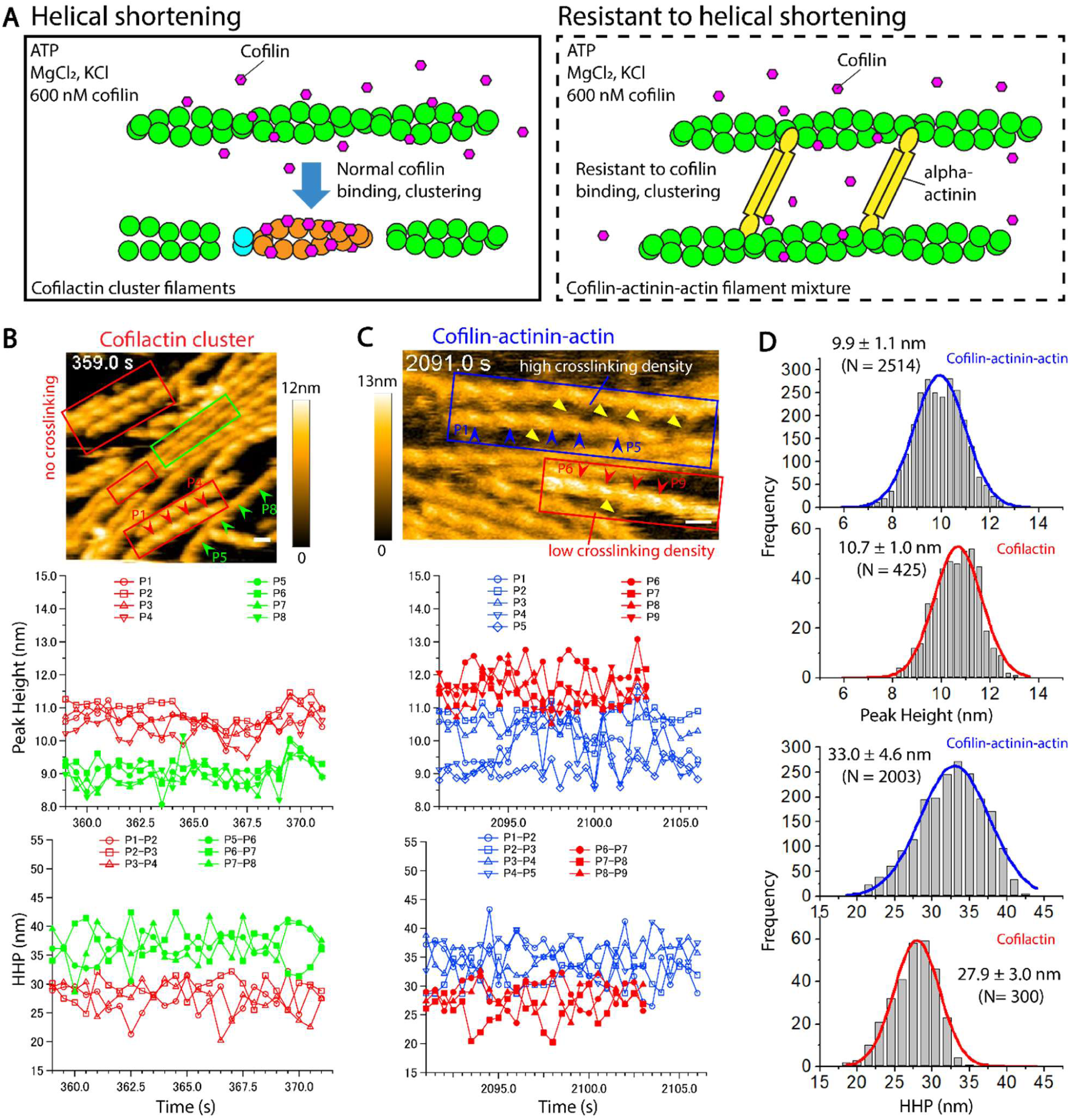
The high crosslinking density of alpha-actinin with actin filaments inhibits cofilin-induced shortening of HHP and the development of full cofilin clusters. **(A)** Experimental setup. **(B)** Time-dependent changes in the peak heights and HHP of actin filaments incubated with only 600 nM cofilin 1 and 1mM ATP. A ∼25% decrease in HHP and the formation of mature cofilin clusters on actin filaments in the absence of alpha-actinin served as positive controls, providing a basis for comparison with the full cofilin clusters and HHP shown in **C**. Red and green arrowheads indicate the peaks in full cofilin clusters (red box) and bare actin segments (green box, served as negative controls), respectively. Time label indicates the time after adding cofilin. Scale bars: 25 nm. Color scale: 0-12 nm. Still AFM image in B was similarly reused in Figure S7. **(C)** Time-dependent changes in the peak heights and HHPs of actin filaments incubated with 400 nM alpha-actinin and 1 mM ATP, and subsequently added with 600 nM cofilin. Red arrowheads denote the peaks in full cofilin clusters within the supertwisted half helices of actin segments (red box, cofilactin clusters) without crosslinking by alpha-actinin, and blue arrowheads indicate those in partial cofilin clusters within the long half helices of actin segments crosslinked with alpha-actinin (blue box, cofilin-actinin-actin filament complex), respectively. The yellow arrowheads denote representative alpha-actinin. Scale bars: 25 nm. Color scale: 0-13 nm. Still AFM image in C was similarly reused in Figure S9. **(D)** Histograms of the peak height and HHP measured from various actin filaments with and without alpha-actinin crosslinking, showing the presence of full cofilactin and partial cofilin-actinin-actin segments. The values are mean ± SD and analyzed from 5-6 filaments for each condition. A two-population *t-test* (*p* < 0.05) indicates that the differences in height and HHP distribution between these segments are significant (*p* = 2.9×10^−38^ for height, *p* = 8.5×10^−68^ for HHP).

**Table 5.** Summary of the peak height and HHP (mean ± SD) in full cofilin clusters (cofilactin) without alpha-actinin and partial cofilin clusters within actin segments crosslinked with alpha-actinin (cofilin-actinin). Statistical differences were assessed using a two-population *t-test*, with a significant difference confirmed at *p* ≤ 0.05. Related to **Figure 2**.

| Segment of | Height (nm) (N) | HHP (nm) (N) |
|---|---|---|
| Cofilactin | 10.7 ± 1.0 (425) | 27.9 ± 3.0 (300) |
| Cofilin-actinin | 9.9 ± 1.1 (2514) | 33 ± 4.6 (2003) |
| *p*-value (*t-test*) | 2.9 × 10<sup>-38</sup> | 8.5 ×10<sup>-68</sup> |

### Myosin II and cofilin preferentially bind to normal and short half helices to facilitate stable cluster formation

Previously, it was experimentally demonstrated that the allosteric regulation of myosin II and cofilin binding to the actin filament occurs without direct competition for binding sites (Ngo et al., 2016), although there is currently no atomic structural evidence to support this hypothesis. To further address this issue, HS-AFM was used to observe the competitive binding of S1 and cofilin to the actin filaments under near-physiological conditions in the presence of ADP and ATP. The actin filaments were gently immobilized on a lipid membrane in F-buffer initially containing 300 nM S1 at pH 6.8, followed by the addition of 100 nM cofilin (**Figure 3, Figure S10, Table 6, Video S5-S6**). Under high-ADP and low-ATP conditions, S1 transiently associated with and dissociated from the actin filaments. Under this condition, cofilin was unable to form clusters even though most of the actin subunits were not occupied by S1 clusters for approximately 10 min following the cofilin addition (**Figure S10A, Video S5**), consistent with our previous fluorescence microscopic observation (Ngo et al., 2016). The histogram of peak heights for all bare actin segments (8.1 ± 0.8 nm, N = 1122) supports the conclusion that cofilin remained unbound or sparsely bound without forming cooperative clusters. The transient interaction of S1 with the actin filaments consumed ATP because both actin and S1 are ATPase enzymes, and since ATP became depleted, S1 clusters gradually formed; however, these S1 clusters were spatially segregated from the cofilin clusters on the same actin filaments. Analysis of the peak heights and HHPs in the stable mature S1-actin and cofilactin segment clusters (**Figure 3A-C**, **Table 6**) revealed that the taller S1 clusters (14.9 ± 1.4 nm (N = 1051) contained longer HHPs (33.9 ± 4.4 nm (N = 805)) than the cofilin clusters (10.5 ± 1.0 nm (N = 935) and 25.7 ± 3.6 nm (N = 767)), as assessed by the statistical significance (*p* < 0.05). Interestingly, these clusters remained segregated, on the same actin filaments under ATP-depleted conditions (**Figure 3, Video S6**). It is likely because normal HHPs, which supported S1 binding; these represented stable half helices similar to those in canonical actin filaments without or with alpha-actinin, whereas the shorter HHPs, which accommodated the cofilin clusters, corresponded to the constrained or supertwisted half helices.

**Figure 3.**
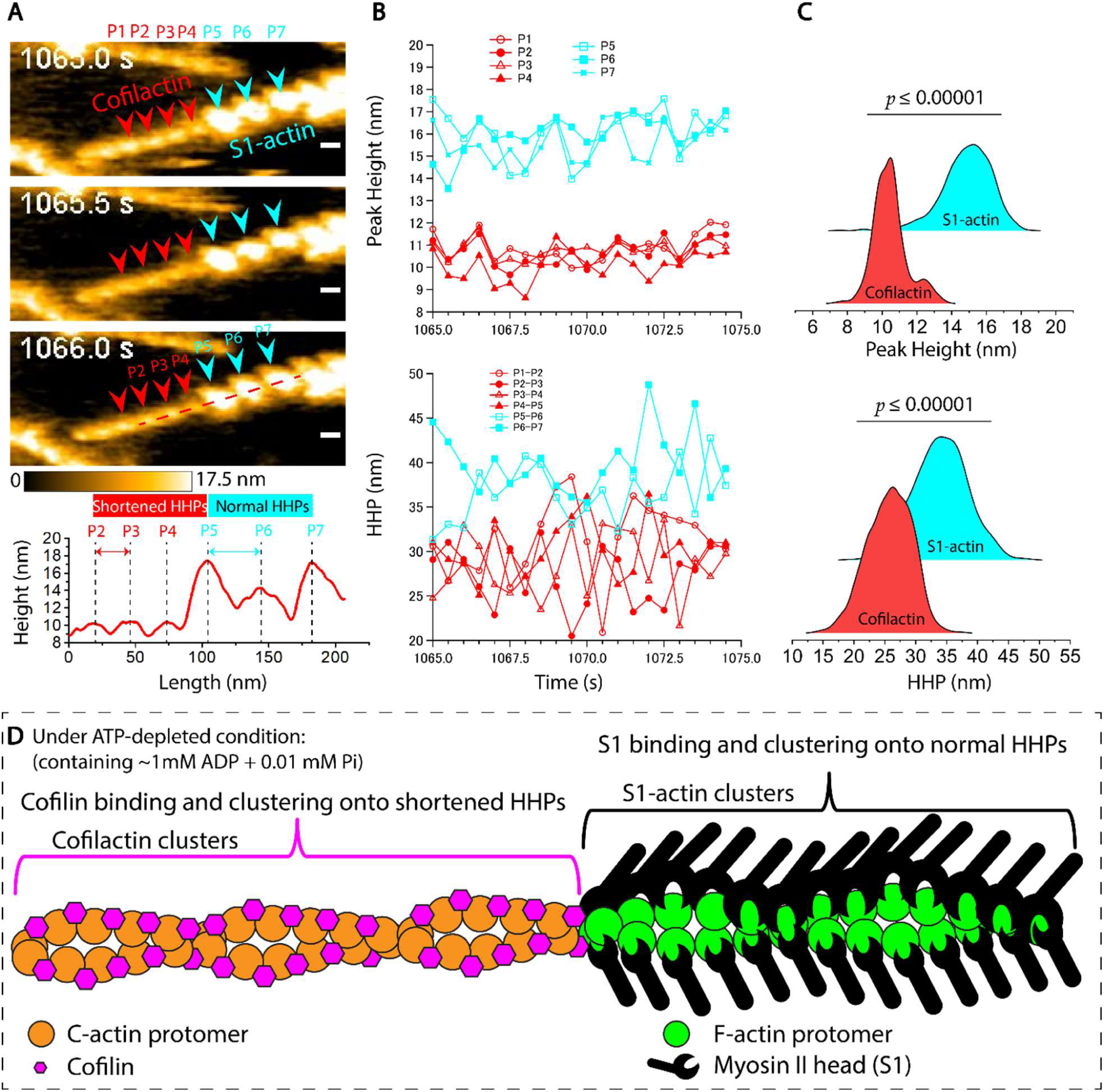
Actin filaments adopt long or short helical pitches to promote the assembly of myosin II (S1) and cofilin clusters, respectively. **(A)** Still AFM images and longitudinal section profile show actin filaments with distinct short and normal HHP regions where cofilin (cofilactin segment) and S1 (S1-actin segment) bind. These filaments represented one of the configurations in **Figure S10**. Actin filaments were initially imaged in F-buffer containing 0.01 mM ATP, 1 mM ADP, and 300 nM S1, followed by adding 100 nM cofilin. Red and cyan arrowheads mark the peak heights determined between every half helical turn in cofilactin and S1-actin cluster regions, respectively. Time labels correspond to frames in **Video S6** after the addition of 100 nM cofilin, during which sTable1 clusters were formed near cofilin clusters on actin filaments when ATP levels were depleted. Scale bars: 25 nm. Color scale: 0-17.5 nm. Still AFM images were similarly reused in Figure S10B. **(B)** Time-dependent variations in peak heights and HHP within cofilactin and S1-actin cluster segments. **(C)** Ridgeline plots displaying the distribution of peak heights and HHP in cofilactin and S1-actin cluster regions. The average peak heights (mean ± SD) are 10.5 ± 1.0 nm (N = 935) for cofilactin and 14.9 ± 1.4 nm (N = 1051 peaks) for S1-actin segments. The average HHPs (mean ± SD) are 25.7 ± 3.6 nm (N = 767 HHPs) for cofilactin and 33.9 ± 4.4 nm (N = 805 HHPs) for S1-actin. Statistical differences between mean values were analyzed using a two-population *t-test* and significant at (*p*<0.00001). **(D)** Sketches illustrating cofilin-induced helical shortening and the formation of corresponding clusters on shortened HHP region, compared to the stable clustering of S1-tuned normal HHP region under ATP-depleted conditions. The differences in protomer conformations are depicted in Figure S10C with reference to previously reported atomic structures (Fujii et al., 2010; Fujii and Namba, 2017; Galkin et al., 2011).

**Table 6.** Summary of the peak height and HHP (mean ± SD) in cofilin clusters (cofilactin) and S1 clusters (S1-actin) segments formed and segregated on the same actin filaments. Statistical differences between the mean values were assessed using a two-population *t-test*, with a significant difference confirmed at *p* ≤ 0.05. Related to **Figure 3**.

| Segment of | Peak Height (nm) (N) | HHP (nm) (N) |
|---|---|---|
| S1-actin | 14.9 ± 1.4 (1051) | 33.9 ± 4.4 (805) |
| Cofilactin | 10.5 ± 1.0 (935) | 25.7 ± 3.6 (767) |
| *p*-value (*t-test*) | <0.00001 | 3.3 ×10<sup>-245</sup> |

Consistent with the results from MD simulations, HS-AFM observations further corroborated that alpha-actinin crosslinking stabilizes actin filament helical periodicity and inter-subunit conformations (MADs), thereby limiting the structural remodeling required for cooperative cofilin binding and cluster formation.

### Transient alpha-actinin ABD binding remodels actin filaments into single-helical, parallel-helical protofilament state, and nano-ring structure without suppressing cofilin cluster formation

The K255E mutation in the actin binding domain (ABD-K255E, *k_d_* ∼ 5.3 µM for actin filament) of non-muscle human alpha-actinin-4 exposes a buried actin-binding site between the CH1 and CH2 domains, increasing its affinity for actin filaments while abolishing Ca²⁺ sensitivity (Shams et al., 2016; Weins et al., 2007). This disease-associated mutation causes inherited kidney disease.

Similarly, the E254K mutation in the human filamin A ABD is a gain-of-function mutant that relieves CH2-mediated autoinhibition in the CH1-CH2 domains, increasing constitutive actin filament binding affinity (Iwamoto et al., 2018).

To evaluate the effects of alpha-actinin binding on actin filament structure, we used human alpha-actinin-1 ABD-E235K mutant, which is homologous to the E254K mutation in filamin A ABD and functionally mimics alpha-actinin-4 ABD-K255E. ABD-E235K bound transiently to actin filaments, with dwell times of 0.39 ± 0.23 s (N = 1171), 0.38 ± 0.18 s (N = 1494), and 0.46 ± 0.24 s (N = 1494) at concentrations of 1, 2, and 3 μM, respectively. Despite these transient interactions, ABD-E235K did not alter the mean HHP (37.1 ± 2.2 nm), which remained comparable to the helical periodicity of canonical actin filaments (**Figure 4A,B; Figure S11, Video S7**).

**Figure 4.**
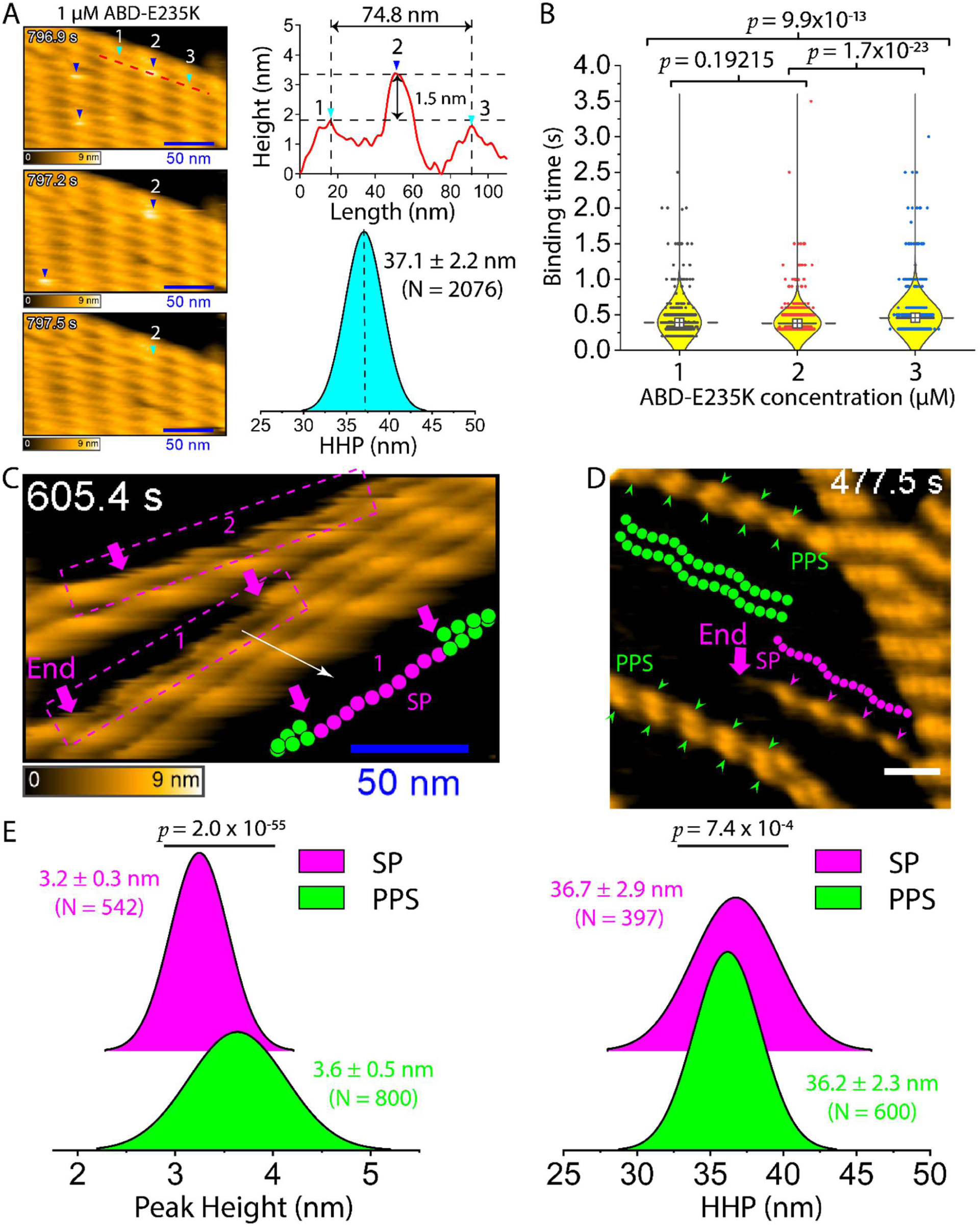
Transient binding of ABD-E235K induces a transition of the canonical right-handed double-helical actin filament to the single-helical protofilament (SP) and parallel-helical protofilament state (PPS), weakly affecting cofilin binding and cluster formation. **(A)** Representative HS-AFM images showing transient binding events of individual ABD-E235K molecules at 1 μM on actin filaments. Length and height profiles confirm binding of single ABD-E235K molecules, corresponding to an apparent height increase of ∼1.5 nm. Notably, ABD-E235K binding events were detected and analyzed at the top of the filaments but were difficult to resolve when the molecule bound to the sides of the filaments. Histograms of the actin filament half helical pitch (HHP) measured in the presence of 1 and 2 μM ABD-E235K are integrated and shown as mean ± SD (37.1 ± 2.2 nm, N = 2,076 HHPs, and analyzed from 10 filaments). Time labels correspond to frames in Video S7 following the addition of ABD-E235K. Blue arrowheads indicate single ABD-E235K molecules bound onto actin filaments. Cyan arrowheads indicate the crossover points of the double-helical actin filament. Still AFM images were similarly reused in Figure S11A. **(B)** Dwell times of ABD-E235K molecules on actin filaments. Mean dwell times were determined as 0.39 ± 0.23 s (N = 1171), 0.38 ± 0.18 s (N = 1494), and 0.46 ± 0.24 s (N = 1494) at ABD-E235K concentrations of 1, 2, and 3 μM, respectively. For each condition, a series of binding events from more than 10 filaments was analyzed. The HS-AFM imaging rate used for binding time quantification ranged between 0.3 – 0.5 second per frame. Statistical significance was assessed using a two-sample *t*-test. (**C, D**) Representative HS-AFM images of the SP and PPS formed in actin filaments following incubation with 2 μM ABD-E235K. Actin filaments were pre-incubated with ABD-E235K for approximately 30 min at room temperature before immobilization on the same lipid membranes for HS-AFM imaging. Purple arrows indicate the ends of the SP. Purple and green arrowheads denote peak heights of the SP and PPS, respectively, used to measure HHP. Scale bars: 50 nm. Still AFM images in C and D were similarly reused in Figure S12B,C. **(E)** Histograms of the peak height and the HHP of the SP and PPS. Mean ± SD values were analyzed from 5 filaments for each and shown, with statistical significance assessed using a two-population *t*-test.

Interestingly, transient ABD-E235K binding occasionally induced a transition from the canonical right-handed double-helical filament to a single-helical protofilament (SP) and parallel-helical protofilament state (PPS) (**Figure 4C,D, Figure S12, Video S8**). In both SP and PPS, the intrinsic helical periodicity of each protofilament remained clearly preserved (HHPs for the SP and PPS: 36.7 ± 2.9 nm and 36.2 ± 2.3 nm, respectively), with no appreciable change in helical length relative to the canonical HHP. By contrast, the apparent filament height was reduced by more than half relative to the canonical double-helical filament, with peak heights of 3.2 ± 0.3 nm for the SP and 3.6 ± 0.5 nm for the PPS (**Figure 4E**). Because the atomic structures of these newly identified filament states remain unknown, we used the reduced height of the SP and PPS as criteria to distinguish these structures from canonical actin filaments. Although the protofilaments were gently adsorbed onto the positively charged lipid membrane, preventing their complete separation in the HS-AFM images, their individual helical organization remained intact. This observation suggests that individual actin protofilaments intrinsically retain their helical conformation even after dissociation from the intact double helix. We confirmed that the SP and PPS images were not an imaging artifact caused by a double AFM tip (**Figure S12**). Notably, similar SP and PPS were also observed when actin filaments were incubated with the calponin homology domain (CHD) of Rng2 (**Figure S13**), which has been captured in our previous study (Hayakawa et al., 2022), suggesting that the CH domains of alpha-actinin and Rng2 may remodel actin filament architecture through a common mechanism.

Next, we pre-incubated actin filaments and ABD-E235K in solution and subsequently examined how cofilin binds and forms clusters on the PPS-like-helical pitch. We refer to this cofilin-bound state as the PPS-cofilin complex, whose atomic structure remain unknown, to distinguish it from canonical short-pitch cofilactin formed on canonical double-helical actin filaments. Interestingly, cofilin also bound to PPS-like filaments, where it formed clusters. However, unlike canonical cofilactin, these PPS-cofilin complex segments did not undergo the characteristic helical shortening, retaining an HHP of 35.9 ± 4.2 nm and a peak height of 9.3 ± 0.6 nm (**Figs. S14-S15, Video S9**).

We next investigated whether the continued presence of free ABD-E235K in solution influences cofilin-induced structural remodeling. Actin filaments were first immobilized on a lipid membrane, after which 2.0 μM ABD-E235K was added and imaged by HS-AFM for ∼30 min before the subsequent addition of 3.5 μM cofilin. Despite prolonged incubation (>40 min), cofilin-decorated filaments exhibited only a modest reduction in HHP to 35.0 ± 3.4 nm while unbound ABD-E235K remained in solution (**Figs. S16, S17A, Video S10A**). By contrast, after buffer exchange to remove unbound ABD-E235K, cofilin rapidly assembled into mature clusters and shortened the HHP to the characteristic value of fully decorated canonical cofilactin segments (27.3 ± 1.8 nm) (**Figure S17B, Video S10B**). This result supports the idea that continuous occupancy of actin by transient bindings of ABD-E235K molecules may also perturb the cofilin-induced structural transition of double-helical actin filaments and delay cofilin cluster maturation.

To further investigate the role of the E235K mutation, we performed similar HS-AFM experiments using wild-type ABD (ABD-WT). Surprisingly, 2 μM ABD-WT also bound to actin filaments and strongly induced the formation of both PPS and new nano-ring structures (**Figure 5, Figure S18, Video S11**). Furthermore, HS-AFM imaging of actin filaments preincubated with 1 μM ABD-WT, following by the immobilization on a lipid membrane, revealed that ABD-WT could bind to and form stable clusters on actin filaments (**Video S12**). These findings suggest that the ABD of alpha-actinin-1, independently of the E235K mutation, can induce structural remodeling of canonical actin filaments, promoting their transition into PPS and/or nano-ring conformations.

**Figure 5.**
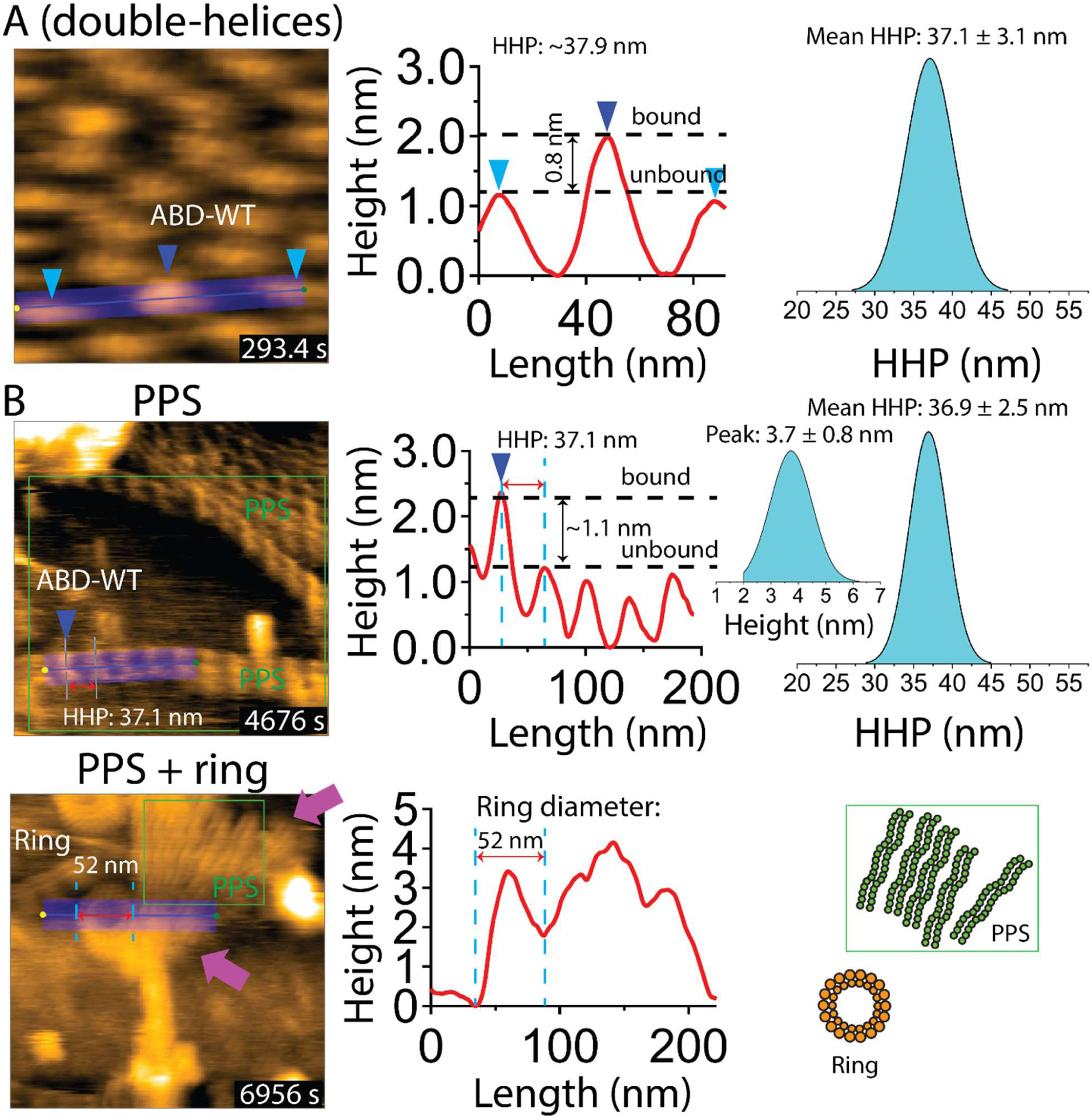
Binding of ABD-WT induces a transition of the canonical right-handed double-helical actin filament to the parallel-helical protofilament state (PPS) and nano-rings. **(A)** Representative HS-AFM images showing the binding of individual ABD-WT molecules at 2 μM on actin filaments. Length and height profiles confirm binding of single ABD-WT molecules, corresponding to an apparent height increase of ∼0.8 nm. Histograms of the actin filament half helical pitch (HHP) measured in the presence of 2 μM ABD-WT are shown as mean HHP ± SD (N = 1355 HHPs, and analyzed from 16 filaments). Blue arrowheads indicate single ABD-WT molecules bound onto the PPS. Cyan arrowheads indicate the crossover points of the double-helical actin filament. **(B)** Representative HS-AFM images of the PPS and nano-rings formed in actin filaments following incubation with 2 μM ABD-WT as shown in A. Histograms of the peak height and the HHP of the PPS. Mean ± SD values are shown (N = 513 HHPs and 668 peak heights analyzed from 5 filaments). Blue arrowheads show the ABD-WT binding on PPS. Purple arrows indicate the ends of the PPS. In both A and B, actin filaments were gently immobilized on lipid membranes and subsequently introduced into the imaging chamber containing 2 μM ABD-WT, followed by HS-AFM imaging at the indicated time points. A still AFM image in B was similarly reused in Figure S18B.

Together, these results demonstrate that transient ABD-E235K binding delays but only weakly inhibits cofilin binding and cluster formation. Nevertheless, the formation of PPS-cofilin complex without PPS-like-helical pitch shortening suggests new properties of cofilin that cofilin can bind to and form clusters on PPS without changing filament helical length.

### Molecular dynamics simulations reveal that alpha-actinin crosslinking preserves the intrinsic helical architecture and cooperative dynamics of actin filaments

To elucidate how alpha-actinin influences the structural dynamics of actin filaments at atomic level, we used complementary all-atom MD simulations and PCA to investigate actin protomers and alpha-actinin-crosslinked actin filaments (**Figure 6**). The computational systems included bare actin filaments together with unipolar and bipolar alpha-actinin-crosslinked filament models constructed from the actin filament structure (PDB: 8D15) and full-length alpha-actinin 2 (PDB: 1SJJ) based on the Taylor’s configuration (Liu et al., 2004), followed by refinement using an experimentally validated Mg²⁺–ADP hydration model of actin (see Methods). Each system was simulated for a total of 500 ns using three replicas, after which 10000 trajectories from each replica, corresponding to 300 ns, were analyzed by PCA to quantify collective conformational motions, including twisting and flattening vibrations of individual actin protomers and alpha-actinin.

**Figure 6.**
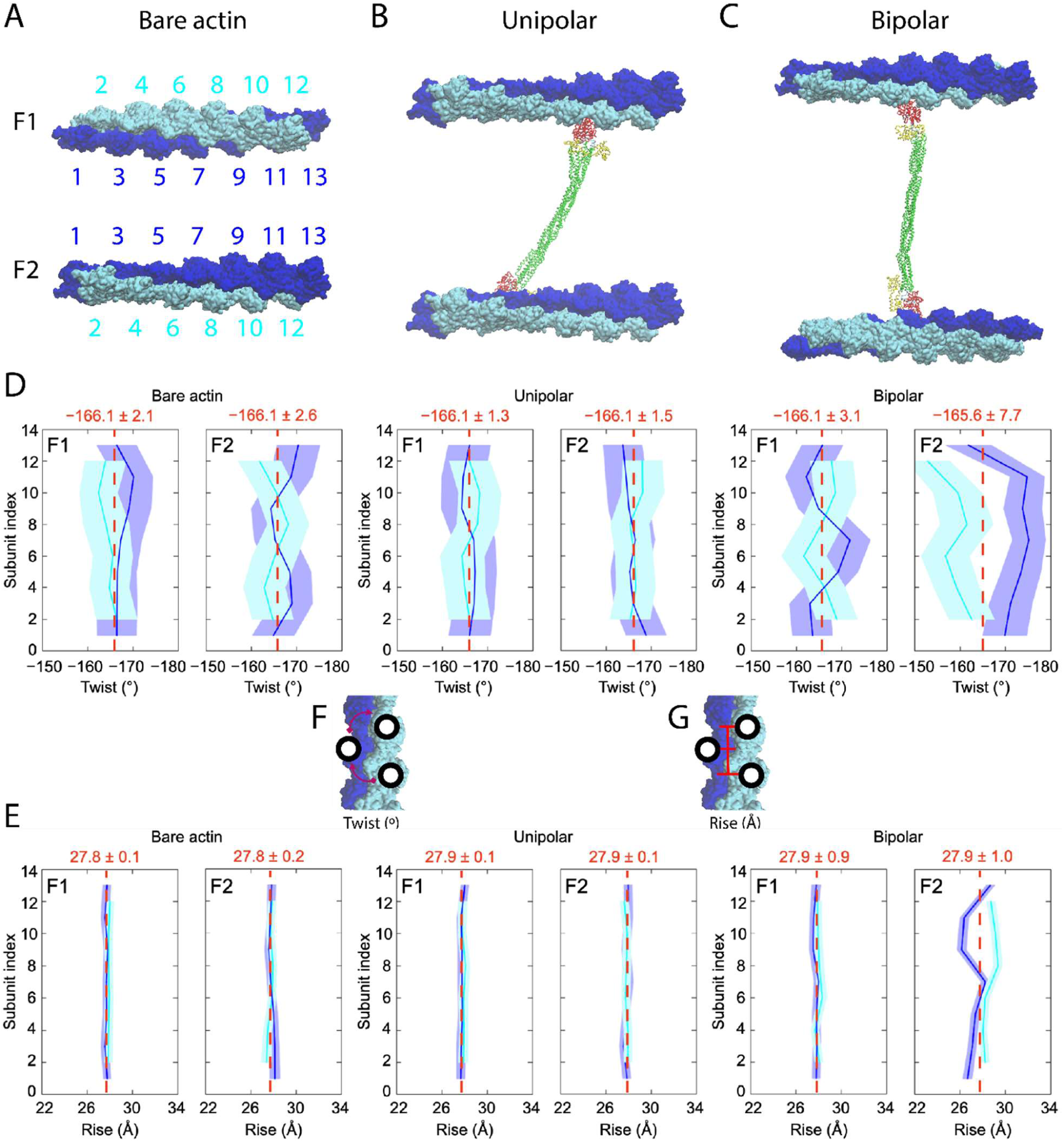
Analysis of actin filament twist and rise by molecular dynamics simulations. **(A–C)** Surface representations of bare actin filaments and alpha-actinin-crosslinked actin filaments in the unipolar and bipolar configurations. The configurations at the last ones during MD simulations are shown. Odd- and even-numbered actin protomers are shown in blue and cyan, respectively, and are labeled sequentially from 1 to 13 in filament 1 (F1) and filament 2 (F2) across all models. Each model comprises 26 actin protomers, either in the absence or presence of a single alpha-actinin molecule (see Methods). **(D, E)** Distributions of the helical parameters, twist and rise, for the actin filaments. Strand colors correspond to those shown in **A–C**. Shaded areas indicate the 70% confidence intervals derived from 30,000 snapshots collected from the 300 ns production run of MD simulations. Vertical red dashed lines denote the mean ± SD values for odd- and even-numbered protomers. (**F, G**) Schematics illustrating the definitions of twist and rise.

To characterize the helical architecture of actin filaments, we employed a previously established method (Carl et al., 2026; Reynolds et al., 2022) to quantify two key helical parameters: the rise, defined as the axial distance between adjacent actin protomers (n and n + 1) on opposite strands of the actin double helix, and the twist, defined as the rotational angle between the same pair of adjacent protomers (**Figure 6F-G**).

Analysis of the MD trajectories revealed that alpha-actinin crosslinking induces only minor perturbations to the global helical architecture of actin filaments. Throughout the 300 ns simulations analyzed, the mean values of both twist and rise remained highly similar between bare and crosslinked filaments, irrespective of whether the filaments adopted unipolar or bipolar arrangements (**Figure 6, Figure S19, Videos S13-S16**). For the unipolar model, the arrangement of actin and alpha-actinin was mostly maintained in the Taylor configuration throughout the simulation (**Figure S6**). In contrast, the bipolar model, which was initially built in the Taylor configuration (**Figure S5A, S6**), gradually and continuously changed alpha-actinin crosslinking orientation during the simulation. The configuration at the last frame in the simulation is shown in **Figure 6A-C**. Moreover, odd- and even-numbered actin protomer conformations exhibited nearly identical mean helical parameters (twist, rise) and comparable distribution widths between bare actin and unipolar models (**Figure 6D-E**). In contrast, the bipolar model exhibited the similar mean helical parameters but greater fluctuation than the bare actin and unipolar models. This indicates that alpha-actinin crosslinking preserves the native double-helical geometry of actin filaments. Rather than altering filament architecture, alpha-actinin primarily stabilizes the spatial organization of neighboring filaments while maintaining the intrinsic structural flexibility of individual filaments.

To investigate whether alpha-actinin affects cooperative filament dynamics, we next quantified the correlations of inter-subunit motions. Representative correlation of twist indices 1, 2, and 3 are illustrated in **Figure S20**. Twist index 1 measures the rotational angle between protomers *n* and *n* + 1, index 2 between *n* + 1 and *n* + 2, and index 3 between *n* + 2 and *n* + 3. Correlation analysis revealed a conserved dynamic network in all filament models, characterized by positive correlations between neighboring subunits (indices 1 and 3) within the same long-pitch strand and negative correlations between adjacent subunits (indices 1 and 2) on opposite strands. This alternating correlation pattern was preserved in both unipolar- and bipolar-crosslinked filaments, indicating that alpha-actinin crosslinking does not perturb the intrinsic cooperative torsional dynamics of actin filaments.

A similar pattern was observed for longitudinal motions. Correlation matrices of rise fluctuations showed positive correlations between neighboring subunits within the same strand (indices 1 and 3) and anticorrelated motions between opposing strands (indices 1 and 2) (**Figure S21**), reflecting the intrinsic axial cooperativity of the actin double helix. These correlation patterns remained essentially unchanged following alpha-actinin crosslinking, suggesting that alpha-actinin constrains the relative positioning of adjacent filaments while preserving the longitudinal flexibility and cooperative dynamics of individual actin protomers.

Collectively, these simulations establish that alpha-actinin functions as a mechanical crosslink that reinforces higher-order actin filament organization without perturbing the native helical geometry or intrinsic cooperative dynamics of actin filaments. This structural framework provides a complementary hypothesis for understanding how alpha-actinin stabilizes actin networks while preserving the conformational plasticity required for cytoskeletal remodeling, force transmission, and the regulated binding of other actin binding proteins.

### Crosslinking-dependent modulation of actin protomer dynamics

To distinguish intra-subunit dynamics from the previously characterized inter-subunit motions (rise and twist), we next performed PCA to identify the principal conformational motions within individual actin protomers. PCA showed that PC1 (accounting for more than ∼13-16% of the total variance) captured a conserved vibration of SD1-SD2 to SD3-SD4 in bare actin, unipolar, and bipolar filaments. Though the conformational fluctuations along PC1 differed among the three models, with the bipolar model exhibiting a broader conformational space than the bare actin and unipolar models, the flattened conformational vibration mode of actin protomers was preserved despite crosslinking (**Figure 7, Video S17**). PC2 (∼7-8% of the total variance) reflected independent subdomain motions, with SD2 rotating and SD3-SD4 moving in opposite directions with SD2 in the bare actin model. Bipolar filaments showed more coordinated behavior, with SD2 undergoing a combined rotation and translation and SD3-SD4 rotating as a single unit. The unipolar model retained SD2 rotation but exhibited a distinct downward displacement of SD4 (**Figure S22**).

**Figure 7.**
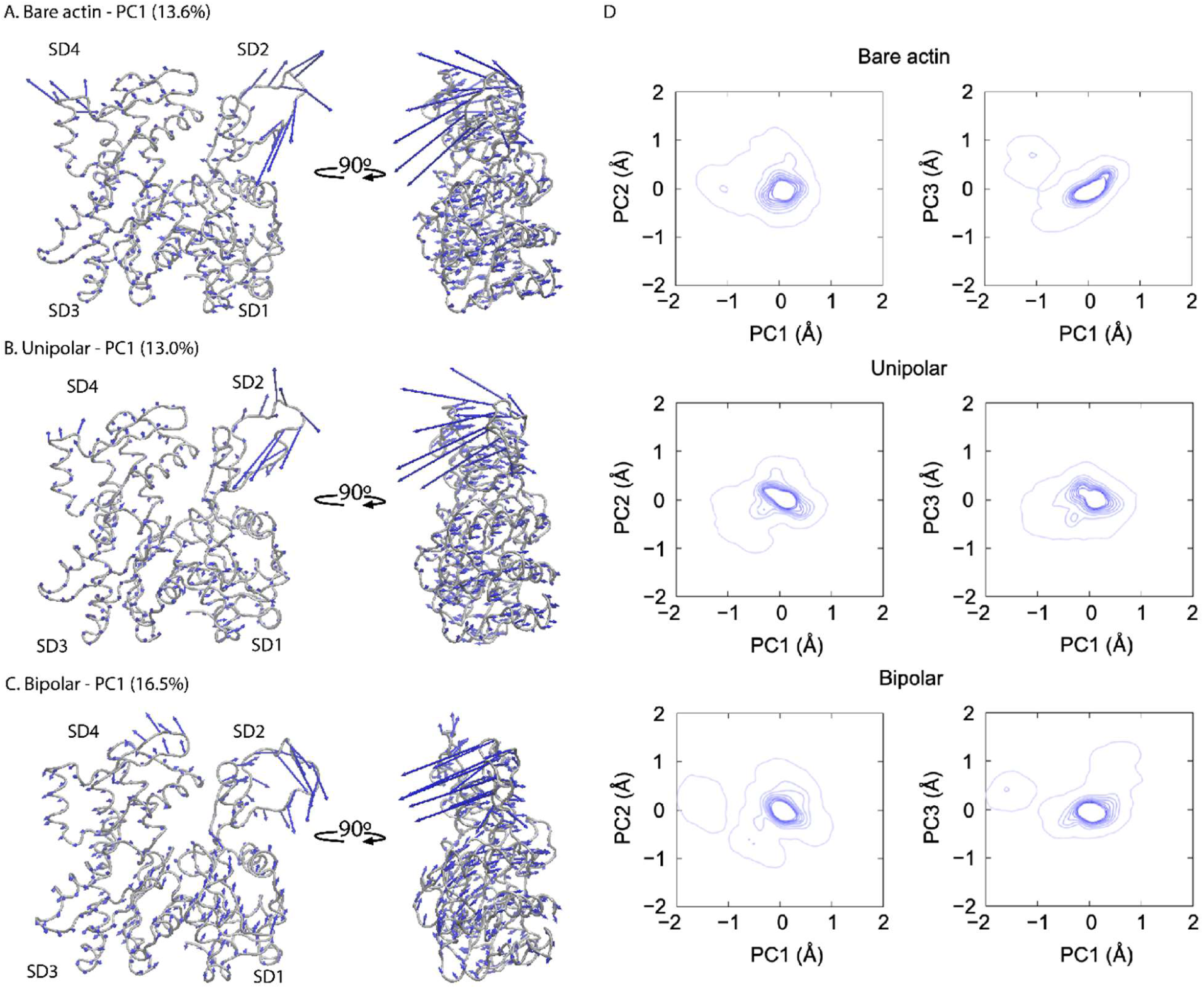
Analysis of atomic conformational dynamics of actin protomers from the bare actin, unipolar, and bipolar filament models using MD simulations and PCA. (**A, B, C**) Principal component 1 (PC1) captured flattening motions of actin protomers in bare actin, unipolar, and bipolar filament models. Actin structures are shown in tube representation with the four subdomains (SD1-SD4) highlighted. The eigenvalues of PC1 indicate the percentage contribution of the dominant motions. Eigenvector components depicting the direction of motion are indicated by blue arrows. (**D**) Distributions of conformational ensembles projected onto the common PC1-PC2 and PC1-PC3 spaces. PCA modes were obtained from conformational libraries generated by MD simulations for 300 ns.

We also observed a different trend for PC3 (∼6-7% of the total variance). Bare filaments most closely resembled the bipolar mode, where SD1 and SD3 rotated in opposite directions and SD2 showed complex motion, whereas the unipolar model displayed same-direction rotation of SD1 and SD3 and inward movement of SD2 toward SD4 (**Figure S23**). The distributions of conformational ensembles projected onto the common PC1-PC2 and PC1-PC3 spaces suggest that PC1 captured the largest protomer-flattening vibrations in the bipolar model, whereas these motions were comparable between the bare actin and unipolar models.

To quantify local flexibility of actin protomers, we calculated the root-mean-square fluctuation (RMSF) of each Cα atom from its average position over the MD trajectories (**Figure S24-S25**). Consistent with the PCA results, the bipolar model exhibited slightly larger fluctuations in SD2-SD4 than either the bare actin or unipolar models, corroborating the enhanced protomer dynamics observed in the common principal component space. Together, these findings indicate that although the principal protomer-flattening vibration is conserved, alpha-actinin crosslinking modulates higher-order dynamic modes (PC1, PC2, PC3) in unipolar and bipolar actin architectures.

### Thermodynamics of conformational changes of actin filament subunits

To characterize the thermodynamic landscape of actin subunit conformations in the bare actin, unipolar, and bipolar models, we calculated two-dimensional free-energy surfaces from the joint probability distributions of the SD2-SD1-SD3-SD4 dihedral angle and the SD2-SD4 distance, which together describe the relative orientation of the four actin subdomains and the degree of protomer flattening (see Methods). Our all-atom MD simulations revealed that the global free-energy minima of the bare actin, unipolar, and bipolar model are nearly identical, exhibiting a single dominant lowest-energy basin. The distributions of the sTabletate are centered at approximately - 5.2 ± 1.9° for the dihedral angle and 2.82 ± 0.05 nm for the SD2-SD4 distance (**Figure 8**), demonstrating that actin protomers in all three models predominantly reside in the canonical flattened low-energy state. The time evolutions of the dihedral angle, SD2-SD4 distance, and potential energy are shown (**Figure S26-S27**). Notably, the bipolar model also accesses an additional higher-energy metastable basin, indicative of enhanced conformational plasticity: unlike the bare actin and unipolar models, the bipolar model displays an additional metastable free-energy state (∼3-5 kcal mol⁻¹ above the global minimum) associated with larger SD2-SD1-SD3-SD4 dihedral angles (0-2.5°), indicating that bipolar crosslinking enables actin protomers to transiently sample an alternative conformational state (**Figure 8**).

**Figure 8.**
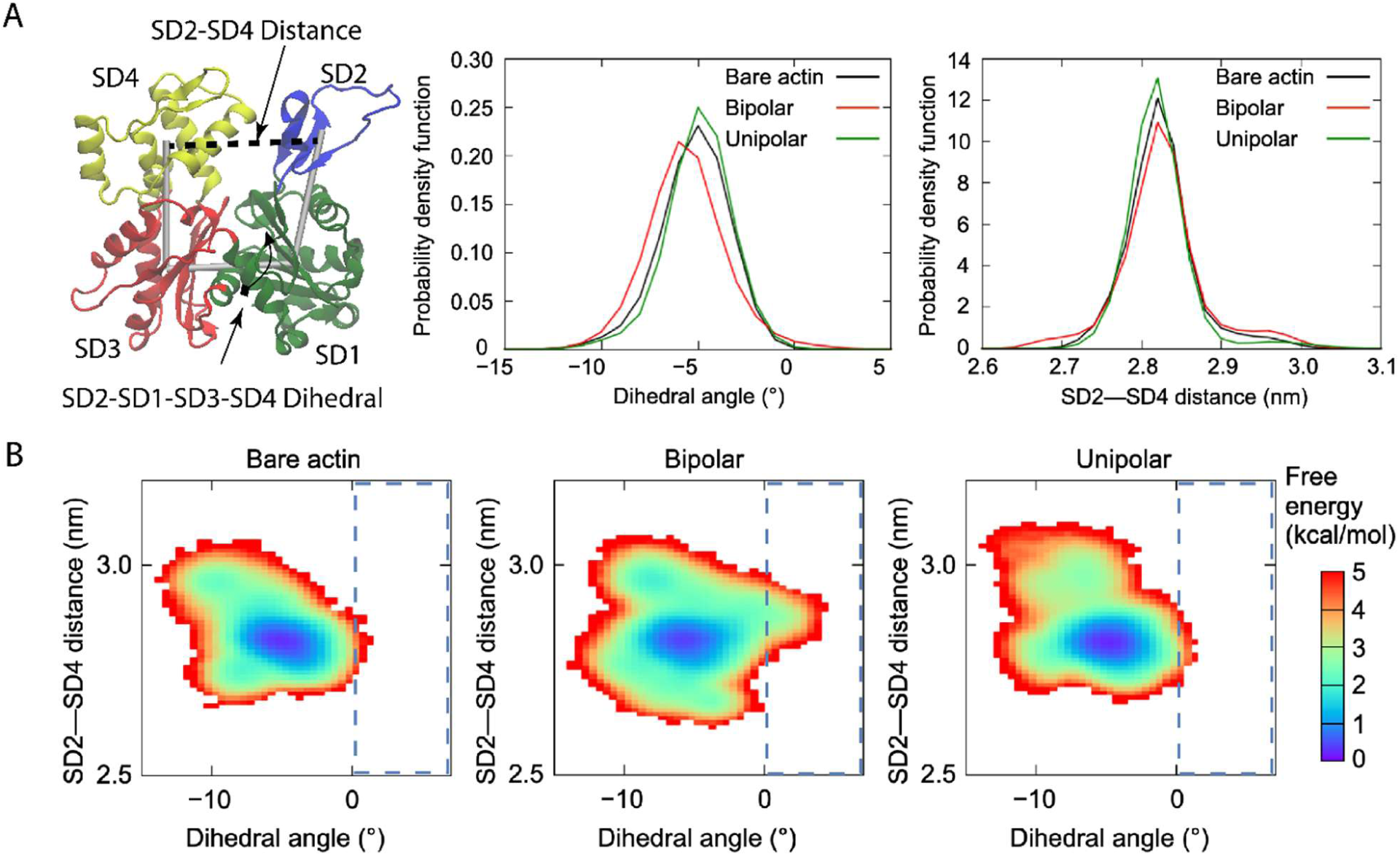
Thermodynamic conformational landscapes of ADP-bound actin subunits in bare and alpha-actinin-crosslinked actin filaments. **(A)** Ribbon representation of an actin monomer highlighting the four subdomains: SD1 (green), SD2 (blue), SD3 (red), and SD4 (yellow). The centers of mass of Cα atoms within each subdomain were used to define the SD2-SD1-SD3-SD4 dihedral angle and the SD2-SD4 inter-subdomain distance. **(B)** Two-dimensional free-energy landscapes of ADP-bound actin subunits from all-atom molecular dynamics (MD) simulations of bare actin filaments and alpha-actinin-crosslinked actin filaments in the unipolar and bipolar configurations. Free-energy surfaces were constructed as a function of the SD2-SD1-SD3-SD4 dihedral angle and the SD2-SD4 distance at 300 K. Blue dashed boxes indicate the positive dihedral-angle region associated with high free energy.

To assign structural identities to the free-energy minima, we compared the resulting landscapes with previously reported conformational states of actin subunits at distinct positions along the filament. In the previous reports, the resulting free-energy landscapes revealed distinct minima corresponding to the flattened and twisted conformations of actin subunits. The internal subunits and the terminal subunit at the barbed end exhibited much smaller dihedral angles (-2.3° and -3.0°, respectively) with short-pitch rotation angles of approximately -166°, indicative of the canonical flattened filament conformation (Iyer et al., 2025). In contrast, the twisted state was characterized by an SD2-SD1-SD3-SD4 dihedral angle of approximately -20.3°, consistent with the conformation adopted by the terminal subunit at the pointed end of actin filaments (Iyer et al., 2025). At the pointed end, however, the terminal subunit (P; -18.5°) and penultimate subunit (P-1; -19.6°; short-pitch rotation angle, -166.3°) retained large dihedral angles characteristic of the twisted conformation, closely resembling that of the actin monomer (Narita et al., 2011).

### Conformational dynamics of alpha-actinin in unipolar and bipolar crosslinking models

We next analyzed the atomic conformational dynamics of alpha-actinin in alpha-actinin-crosslinked actin filaments under both unipolar and bipolar models, using all-atom MD simulations and PCA (**Figure 9, Figure S28, Videos S18-S19**).

**Figure 9.**
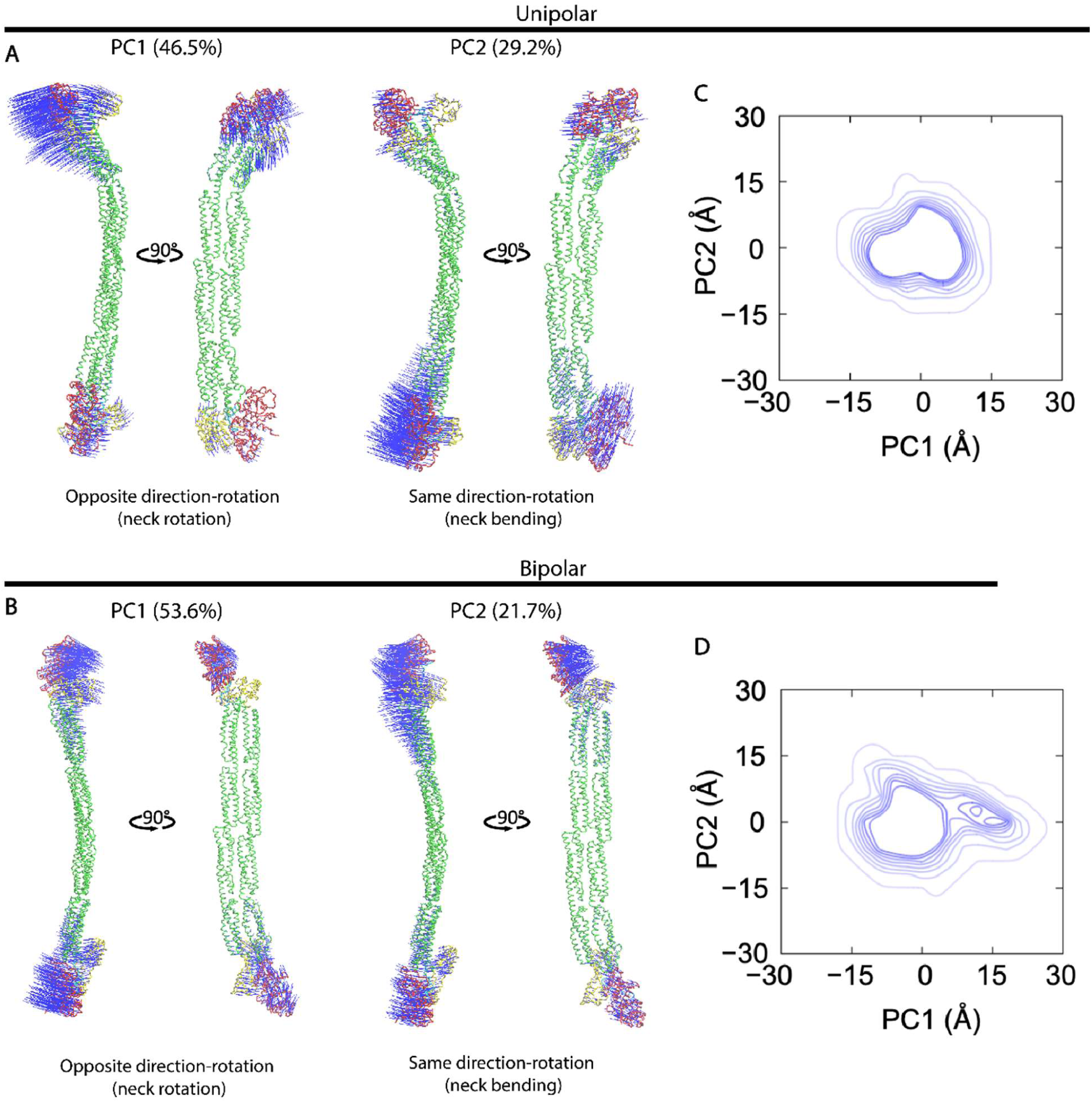
Analysis of the atomic conformational dynamics of alpha-actinin derived from the unipolar and bipolar models using all-atom MD simulations and PCA. (**A, B**) The alpha-actinin structures are illustrated in tube conformations, with the actin binding domain (ABD), neck, rod, and EF12-EF34 domains represented in red, cyan, green, and yellow, respectively. The calponin homology 1 and 2 (CH1-CH2) domains in alpha-actinin ABDs retained in open conformations when binding to actin filaments. The eigenvalues of principal components 1 (PC1) and PC2 were used to determine the percentage contribution of each motion. Eigenvector components indicating the direction of motion are shown with blue arrows. (**C, D**) Distributions of conformational ensembles projected onto the common PC1-PC2 space. PCA modes were obtained from conformational libraries generated by MD simulations.

In the unipolar configuration, PC1, PC2, and PC3 primarily captured opposite-direction neck rotation (46.5% of the total variance), same-direction neck bending (29.2%), and same-direction neck twisting (9.1%), respectively, as indicated by the trajectories marked with blue arrows (**Figure 9A, Figure S28A, Video S18**). In the bipolar model, as the same as in the unipolar configuration, PC1 and PC2 corresponded to opposite-direction neck rotation (53.6%) and same-direction neck bending (21.7%), respectively, but PC3 corresponded to an opposite-direction neck twisting (8.9%) (**Figure 9B, Figure S28B, Video S19**). Projection of the conformational ensembles onto the common PC1-PC2 and PC1-PC3 spaces revealed that PC1 captured greater opposite-direction neck rotation in the bipolar model than in the unipolar model. In contrast, the conformational ensembles described by PC2 and PC3 largely overlapped between the two models, indicating similar neck bending and twisting motions (**Figure 9C,D, Figure S28C,D**). RMSF analysis of Cα atoms further supported these findings by showing greater flexibility of the CH1-CH2 domains in the bipolar model (**Figure S28E**). Together, these results demonstrate that alpha-actinin undergoes enhanced neck rotation in the bipolar configuration, whereas its neck bending and twisting motions remain largely conserved regardless of crosslinking geometry.

It is worth noting that experimentally generating bipolar crosslinking on a two-dimensional lipid surface is technically challenging, as shown in previous EM studies (Meyer and Aebi, 1990; Taylor et al., 2000) and our HS-AFM observations, making experimental imaging difficult (see Methods). Thus, MD simulations provide a powerful means to explore alpha-actinin and actin filaments in the bipolar model.

We next compared the atomic conformation and dynamics of alpha-actinin in the unipolar and bipolar models with those in the free state (Ngo et al., 2026). In both crosslinking models, the tandem CH1-CH2 domains within the ABDs adopted an open and flexible conformation (**Figure 9, Video S18-S19**), in contrast to the closed state observed in the models of free alpha-actinin (Ngo et al., 2026) and smooth muscle bound-alpha-actinin (Liu et al., 2004). These findings highlight a significant difference in the conformation of alpha-actinin ABDs when bound to actin filaments versus in its free, unbound state, and consistent with the previous structural study with open ABDs in bound-alpha-actinin (Galkin et al., 2010).

### Residue-specific alpha-actinin–actin interactions underlie configuration-dependent filament dynamics

We aim to map the residue-level interactions between alpha-actinin and actin in the unipolar and bipolar assemblies by analyzing residue contact probabilities. To distinguish intrinsic actin dynamics from motions coupled to alpha-actinin, we additionally analyzed a bipolar-fixed model in which the alpha-actinin were restrained. Comparison of the flexible and fixed models allows us to determine whether correlated motions arise from dynamic actin–alpha-actinin interaction or from the bipolar crosslinking architecture itself.

The local residue contact probabilities between the two ABDs of alpha-actinin and the actin protomers showed substantial differences among the unipolar-flexible, bipolar-flexible, and bipolar-fixed crosslinking models (**Figure S29-S30, Table 7**). The actin monomer consists of 4 subdomains defined as follows: subdomain 1 (SD1), residues 1-32, 70-144, and 338-375; subdomain 2 (SD2), residues 33-69; subdomain 3 (SD3), residues 145-180 and 270-337; and subdomain 4 (SD4), residues 181-269. Our MD simulations identified key alpha-actinin–actin residue pairs with contact-based probability greater than 0.5, as detailed here: LEU26–ASP56 (SD2), LEU26–GLU93 (SD1), ASP27–ARG95 (SD1), ASP59–SER350 (SD1), GLU115–LYS50 (SD2), ARG233–ASP25 (SD1), and GLU236–ARG28 (SD1) in the unipolar-flexible model; GLU31–ARG95 (SD1), GLU149–ARG95 (SD1), ASP201–ARG147 (SD3), GLU226–SER350 (SD1), ARG233–GLU57 (SD2), GLU236–ARG28 (SD1), and ASP867–LYS373 (SD1) in the bipolar-flexible model; and ASP27–ASN92 (SD1), ARG233–ASP56 (SD2), ARG233–GLU167 (SD3) and GLU236–ARG95 (SD1) in the bipolar-fixed model. The time evolutions of these interaction pairs were shown to be highly stable (**Figure S30**), though different from contact residues shown in Liu and Taylor model (Liu et al., 2004). These model-dependent contact patterns refine the residue-level map of the alpha-actinin–actin interface reported previously (McGough et al., 1994; Sutherland-Smith et al., 2003) and indicate that alpha-actinin ABD interactions are concentrated predominantly on SD1, with additional contacts involving SD2 and SD3 but no high-probability contacts detected with SD4.

**Table 7.** A summary of residue-level contact probability maps between alpha-actinin and actin filaments, derived from all-atom MD simulations using three crosslinking models. The unipolar-flexible crosslinking model represents parallel actin filaments with the same polarity, crosslinked by alpha-actinin, which flexibly adjusted their conformation during simulations. In contrast, the bipolar-flexible and bipolar-fixed crosslinking models describe anti-parallel actin filaments with opposite polarity, also crosslinked by alpha-actinin, but either allowing flexible conformational changes or maintaining fixed Cα positions during simulations. Key alpha-actinin ABD-actin interacting residue pairs were identified based on contact probability densities exceeding 0.5. Related to **Figure S29-S30**.

| Alpha-actinin - Actin | Unipolar (300 ns) | Bipolar (300 ns) | Bipolar fixed (100 ns) | Unipolar in Taylor | Bipolar in Taylor |
|---|---|---|---|---|---|
| 26 LEU 17 VAL |  |  |  |  | 1 |
| 26 LEU 21 PHE |  |  |  | 1 |  |
| 26 LEU 28 ARG |  |  |  | 1 |  |
| 26 LEU 31 PHE |  |  |  |  | 1 |
| 26 LEU 56 ASP | 0.647467 |  |  |  |  |
| 26 LEU 92 ASN |  |  | 0.3304 |  |  |
| 26 LEU 93 GLU | 0.540367 |  |  |  |  |
| 27 ASP 28 ARG |  |  |  | 1 |  |
| 27 ASP 29 ALA |  |  |  |  | 1 |
| 27 ASP 31 PHE |  |  |  |  | 1 |
| 27 ASP 50 LYS |  | 0.394433 |  |  |  |
| 27 ASP 56 ASP |  |  |  |  | 1 |
| 27 ASP 88 HIS |  |  |  |  | 1 |
| 27 ASP 92 ASN |  |  | 0.6876 |  |  |
| 27 ASP 95 ARG | 0.676 |  |  |  |  |
| 28 PRO 88 HIS |  |  |  |  | 1 |
| 29 ALA 88 HIS |  |  |  |  | 1 |
| 29 ALA 93 GLU |  |  |  | 1 |  |
| 29 ALA 95 ARG |  |  |  | 1 |  |
| 30 TRP 57 GLU |  |  |  |  | 1 |
| 31 GLU 87 HIS |  |  |  |  | 1 |
| 31 GLU 92 ASN |  |  |  |  | 1 |
| 31 GLU 95 ARG |  | 0.7094 |  |  |  |
| 34 GLN 91 TYR |  |  |  |  | 1 |
| 59 ASP 350 SER | 0.6115 |  |  |  |  |
| 61 ARG 46 GLY | 0.3424 |  |  |  |  |
| 107 LYS 96 VAL |  |  |  |  | 1 |
| 107 LYS 100 GLU |  | 0.342267 |  |  | 1 |
| 108 LEU 99 GLU |  |  |  |  | 1 |
| 108 LEU 100 GLU |  |  |  |  | 1 |
| 109 VAL 5 THR |  |  |  |  | 1 |
| 109 VAL 100 GLU |  |  |  |  | 1 |
| 109 VAL 96 VAL |  |  |  | 1 |  |
| 110 SER 91 TYR |  |  |  | 1 |  |
| 110 SER 95 ARG | 0.416033 |  |  |  |  |
| 110 SER 97 ALA |  |  |  | 1 |  |
| 110 SER 98 PRO |  |  |  |  | 1 |
| 110 SER 100 GLU |  |  |  |  | 1 |
| 111 ILE 96 VAL |  |  |  |  | 1 |
| 111 ILE 97 ALA |  |  |  |  | 1 |
| 111 ILE 98 PRO |  |  |  |  | 1 |
| 112 GLY 98 PRO |  |  |  |  | 1 |
| 114 GLU 128 ASN |  |  |  |  | 1 |
| 115 GLU 50 LYS | 0.753067 |  |  |  |  |
| 115 GLU 87 HIS |  |  |  | 1 |  |
| 120 ASN 87 HIS |  |  |  |  | 1 |
| 120 ASN 92 ASN |  |  |  | 1 |  |
| 122 LYS 87 HIS |  |  |  |  | 1 |
| 122 LYS 88 HIS |  |  |  | 1 |  |
| 122 LYS 92 ASN |  |  |  | 1 |  |
| 123 MET 91 TYR |  |  |  | 1 | 1 |
| 123 MET 92 ASN |  |  |  | 1 |  |
| 126 GLY 91 TYR |  |  |  |  | 1 |
| 127 MET 97 ALA |  |  |  |  | 1 |
| 129 TRP 95 ARG |  |  |  |  | 1 |
| 129 TRP 96 VAL |  |  |  |  | 1 |
| 130 THR 97 ALA |  |  |  |  | 1 |
| 133 LEU 100 GLU |  |  |  |  | 1 |
| 144 GLU 101 HIS |  |  |  |  | 1 |
| 145 THR 6 THR |  |  |  |  | 1 |
| 146 SER 28 ARG |  |  |  |  | 1 |
| 148 LYS 28 ARG |  |  |  |  | 1 |
| 148 LYS 95 ARG |  |  |  |  | 1 |
| 149 GLU 28 ARG |  |  | 0.4572 |  | 1 |
| 149 GLU 95 ARG |  | 0.587567 | 0.4473 |  |  |
| 201 ASP 147 ARG |  | 0.614433 |  |  |  |
| 226 GLU 350 SER |  | 0.756067 |  |  |  |
| 230 GLY 47 MET |  | 0.4291 |  |  |  |
| 233 ARG 25 ASP | 0.667867 |  |  |  |  |
| 233 ARG 29 ALA |  |  |  |  | 1 |
| 233 ARG 56 ASP |  |  | 0.6359 |  | 1 |
| 233 ARG 57 GLU |  | 0.9218 |  |  |  |
| 233 ARG 93 GLU |  | 0.4402 |  |  |  |
| 233 ARG 167 GLU |  |  | 0.9458 |  |  |
| 234 PRO 95 ARG |  |  |  |  | 1 |
| 235 ASP 91 TYR |  |  |  |  | 1 |
| 235 ASP 95 ARG |  |  |  |  | 1 |
| 236 GLU 28 ARG | 0.8153 | 0.609067 |  |  |  |
| 236 GLU 92 ASN |  |  |  |  | 1 |
| 236 GLU 95 ARG |  |  | 0.9841 |  | 1 |
| 237 LYS 91 TYR |  |  |  |  | 1 |
| 237 LYS 95 ARG |  |  |  | 1 | 1 |
| 862 PRO 353 GLN |  |  |  |  | 1 |
| 863 TYR 350 SER |  |  |  |  | 1 |
| 863 TYR 353 GLN |  |  |  |  | 1 |
| 866 ARG 364 GLU |  | 0.342 |  |  |  |
| 867 ASP 373 LYS |  | 0.611233 |  |  |  |
| 869 VAL 350 SER |  |  |  |  | 1 |
| 869 VAL 351 THR |  |  |  |  | 1 |
| 870 PRO 350 SER |  |  |  |  | 1 |
| 871 GLY 350 SER |  |  |  |  | 1 |
| 872 ALA 350 SER |  |  |  |  | 1 |

## DISCUSSION

We combined HS-AFM, MD simulations, and PCA to investigate how alpha-actinin regulates actin filament architecture and conformational dynamics through different crosslinking configurations. HS-AFM demonstrated that full-length alpha-actinin stabilizes actin helical periodicity and suppresses cofilin-induced supertwisting and cluster formation, with the inhibitory effects strongly dependent on alpha-actinin crosslinking density. Notably, cofilin can still bind to and form clusters along a single actin protofilament without inducing the characteristic helical-pitch shortening observed in canonical cofilactin. Furthermore, myosin II (S1) and cofilin preferentially bind to normal and shortened half-helical structures, respectively. Interestingly, transient binding of the alpha-actinin ABD, including both E235K mutant and wild-type ABD, induces structural remodeling into SP, PPS, and nano-ring conformations while retaining the ability to support cofilin binding and cluster formation. MD simulations further reveal that alpha-actinin crosslinking largely preserves the intrinsic helical architecture and cooperative torsional and longitudinal dynamics of actin filaments, although bipolar crosslinking enhances protomer flexibility and conformational plasticity. Together, these findings establish alpha-actinin as a mechanical regulator that stabilizes higher-order actin organization while modulating filament conformational states and the accessibility of actin-binding proteins.

### High crosslinking densities of full-length alpha-actinin stabilize actin filament architecture to suppress cofilin binding

Our HS-AFM observations show that high-density full-length alpha-actinin crosslinking better preserves canonical actin helical periodicity than low-density crosslinking or bare actin filaments in the presence of cofilin, thereby suppressing cofilin-induced supertwisting and cluster formation. Although distinguishing alpha-actinin-bound from unbound actin protomers directly in HS-AFM images remains challenging, MD simulations revealed broadly similar conformational dynamics between the two populations, with bound protomers exhibiting only slightly larger conformational changes along PC1 (**Figure S31-S32, Videos S20-S21**). These findings suggest that full-length alpha-actinin suppresses cofilin cooperativity primarily through filament-level crosslinking and structural stabilization, rather than through substantial changes in the intrinsic conformational landscape of individual actin protomers.

### Actin protomer twisting and helical shortening drive cofilin binding to canonical actin filament

We propose that the cooperative process of cofilin binding and cluster formation on canonical double-helical actin filaments likely proceeds through the following sequence: initial cofilin binding and cluster forming onto shortened HHP → protomer twisting → HHP shortening accompanied by a reduction in the number of protomers per HHP → enhanced cooperative binding and cluster growth (**Figure 10**). Importantly, two prerequisite steps including protomer twisting and HHP shortening must occur during these processes for cooperativity to emerge. In contrast, when these steps are prevented, cooperative cofilin binding and cluster formation canonical actin filament are significantly perturbed. This cooperativity may align with an induced-fit mechanism (Koshland, 1958), in which cofilin cluster-induced allosteric conformational changes in actin protomers play a central role in promoting helical shortening and full cofilin binding and clustering. Conformational transitions between twisted and flattened actin states have been associated with the dynamic balance between actin polymerization and depolymerization. Twisted G-actin monomers undergo flattening upon polymerization to form actin filament with flattened F-actin form (Oda et al., 2009), whereas the reverse transition accompanies filament depolymerization (Ngo et al., 2024). In addition, cofilin binding and clustering induce a shortening in filamentous double helices, generating twisted actin subunits characteristic of C-actin form within cofilin-decorated HHP region (Galkin et al., 2011; Ngo et al., 2024; Tanaka et al., 2018). Our findings highlight two distinct conformational states within actin filaments: a flattened F-actin conformation that is maintained in alpha-actinin-crosslinked filaments, and a twisted ADP-bound C-actin conformation associated with supertwisted or shortened helices. These structural states likely govern the selective interaction of ABDs, wherein cofilin preferentially associates with and clusters on twisted ADP-bound actin protomers and shortened helices, while S1 exhibits higher affinity for the flattened actin conformation and normal helices (**Figure 10A-B**).

**Figure 10.**
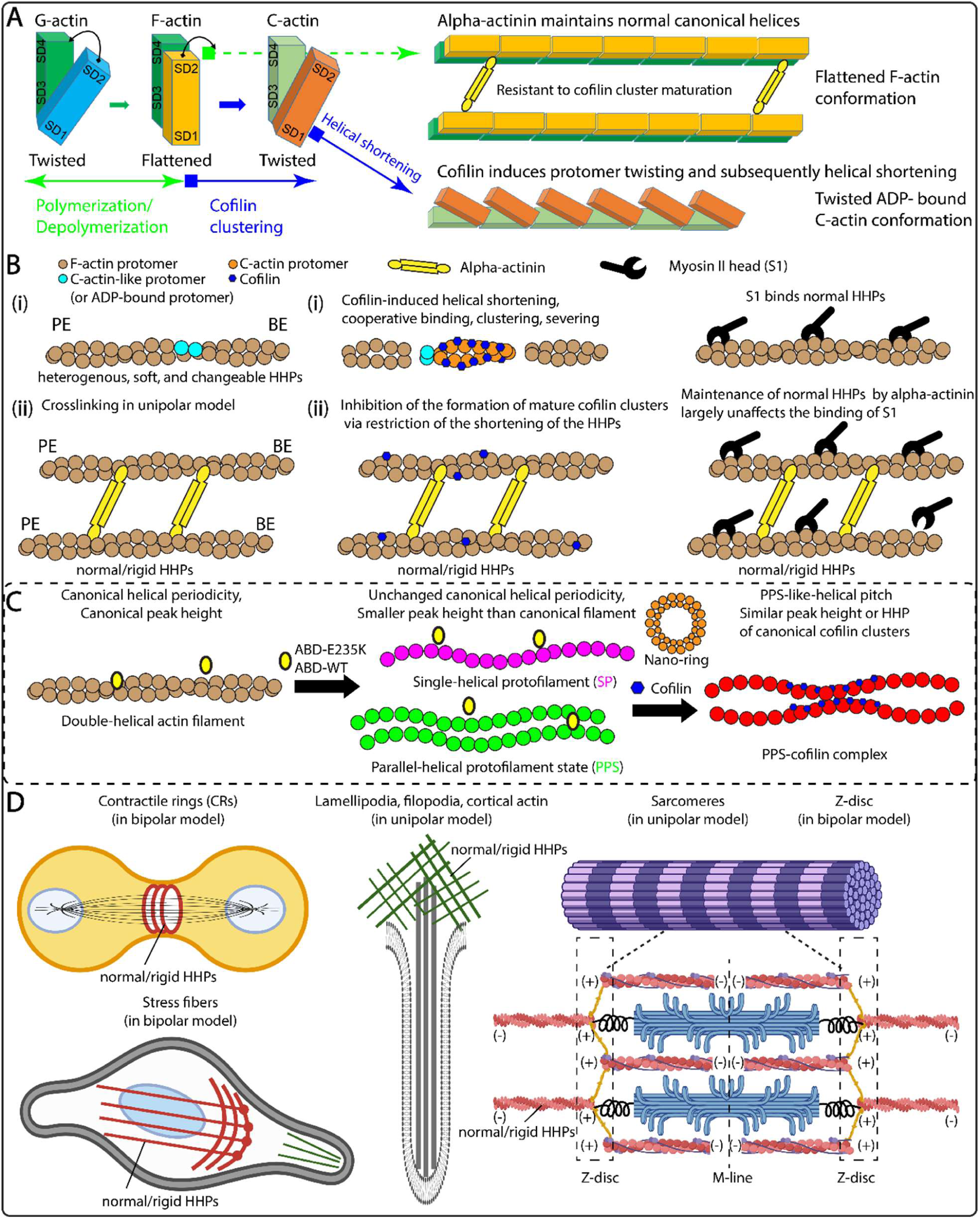
A hypothesis proposing conformation-dependent cooperative binding and clustering of cofilin on canonical actin filaments, utilizing a short helical structure, in contrast to the normal helical structure employed by the myosin II head motor (S1). (**A, B**) Twisted and flattened conformational transitions between G-actin, F-actin, and C-actin structures have been previously linked to actin polymerization and depolymerization and cofilin clustering. Specifically, twisted G-actin polymerizes to form flattened actin filament protomers (F-actin form) (Fujii et al., 2010; Pfaendtner et al., 2010) and vice versa (Ngo et al., 2024), while cofilin-induced clustering leads to twisted protomers (C-actin), and subsequently shortening the HHP in actin filament accompanied by a reduction in the number of protomers per HHP (Galkin et al., 2011; Ngo et al., 2024; Tanaka et al., 2018). Our current study deciphers that alpha-actinin crosslinking helps preserve the canonical double-helical structure of actin filaments, with actin protomers most likely adopting a flattened conformation (F-actin form). This conformation facilitates S1 binding but prevents cofilin-induced helical shortening and cluster maturation. In contrast, the twisted conformation of actin protomers induced by cofilin binding and clustering induces the shortening of helical twists, which supports cofilin’s cooperative binding and clustering (Ngo et al., 2024; Tanaka et al., 2018). The BE and PE denote the barbed end and pointed end of actin filaments, respectively. **(C)** Transient binding of alpha-actinin-1 ABD-E235K or ABD-WT induces the formation of the new SP, PPS, and nano-ring structure from the canonical double-helical actin filaments. The SP and PPS retain a helical periodicity similar to that of canonical actin filaments but exhibit substantially reduced peak heights. Cofilin forms clusters on the PPS to generate PPS-cofilin complexes, which retain the PPS-like-helical pitch while showing increased peak heights with cofilin cluster formation. **(D)** Illustration depicting alpha-actinin-crosslinked actin filaments under the unipolar and bipolar conformations, highlighting their distinct localizations and functions within cells. The plus end (+) and minus end (-) denote the BE and PE. This figure was created using subscribed BioRender.com.

This interpretation agrees with our earlier fluorescence-based results and reinforces our prior hypothesis (Ngo et al., 2016). Nevertheless, without an atomic structure, it remains unclear whether the twisting of protomers in the shortened helical forms prevents S1 binding, a mechanism that would differ from the previously proposed direct competition between cofilin and S1 for actin filament binding sites (Wiggan et al., 2012). The observation of spatially segregated cofilin and S1 clusters on shortened and normal HHPs, respectively (**Figure 3, Video S6**), is consistent with the possibility that cofilin and S1 preferentially interact with twisted and flattened actin protomers, respectively, as referred to previously reported atomic structures (Fujii et al., 2010; Fujii and Namba, 2017; Galkin et al., 2011).

Nevertheless, during long-term HS-AFM imaging in the presence of ADP and ATP at pH 6.8, conditions under which S1 transiently interacts with actin filaments, we observed that many actin binding sites remained accessible to cofilin. Yet, stable cofilin clusters did not form under these conditions, suggesting that S1 and cofilin binding to different actin protomer conformations, leading to mutual exclusive binding as suggested previously (Ngo et al., 2016), and that their stable cluster formation occurs only when sTable1 clustering takes place under ATP-depleted conditions (**Figure 3, Figure S10, Videos S5-S6**). Notably, S1 clusters were observed on normal HHP regions, whereas cofilin preferentially bound to shortened HHP regions. These segregations cannot be simply attributed to cofilin’s preference for ADP-bound actin, as recent atomic structural studies indicate that ADP·BeF₃, ADP·Pi, and ADP-actin filaments share remarkably similar conformations (Oosterheert et al., 2022; Reynolds et al., 2022), suggesting that both ADP-actin and ADP·Pi-actin protomers adopt similarly flattened structures, neither of which represent the favorable protomer conformation required for cofilin cooperativity.

Previously, we observed that ADP-actin filaments contained a higher proportion of short HHPs compared with ADP·Pi-actin filaments. Furthermore, the HHPs in ADP-actin filaments appeared to be more dynamic and flexible than those in ADP·Pi-bound filaments. These characteristics may account for the preferential binding and clustering of cofilin on ADP-actin filaments, as cofilin favors ADP-bound (Suarez et al., 2011) and shorter helical structures (Ngo et al., 2024). Notably, the presence of inorganic phosphate (Pi) only modestly impedes the propagation of the cofilin-induced helical shortening from the decorated cluster into the adjacent bare region of ADP·Pi-actin filaments, thereby still permitting cofilin clustering toward the pointed end side, albeit less efficiently than in the ADP state.

Based on these observations, we hypothesized that cofilin clusters induce twisting of actin protomers and helical shortening within cofilin-decorated region, thereby triggering allosteric, cooperative conformational changes in neighboring protomers within a shortened bare half-helix on the pointed end side (Ngo et al., 2024, 2015). This conversion, from flattened to twisted protomer conformations, was predicted to facilitate helical shortening and further cofilin recruitment and fully cooperative clustering, largely independent of the actin filament’s nucleotide-bound state. Supporting this mechanism, our experiments here show that high alpha-actinin crosslinking densities maintain stable and normal HHPs, more effectively resist cofilin-induced helical shortening as well as cooperative cofilin binding and cluster formation compared with bare filaments. In contrast, bare actin filaments display the expected cofilin-induced helical shortening and cooperative cluster.

Alpha-actinin crosslinking supports distinct cellular functions depending on filament configuration: bipolar bundles contribute to contractile architectures, and stress fibers, whereas unipolar bundles organize sarcomeres, protrusive and cortical actin networks such as lamellipodia and filopodia (**Figure 10D**, **Table 1**). Our HS-AFM observation and MD simulation results suggest that alpha-actinin-crosslinked actin filaments preserve stable and rigid helical structures. How these structural arrangements influence cofilin binding and regulation at the cellular level remains an open question.

### Collective outer domain twisting drives helical shortening of canonical actin filament

Actin protomer twisting, HHP shortening, and the corresponding reduction in the number of protomers per HHP are geometrically coupled through the collective twisting of the outer domains (ODs) of multiple protomers, as described in our previous study (Ngo et al., 2024) and further explained here and in the Methods section “*From local protomer twisting to global helical shortening*”. When two or more cofilin molecules bind to actin protomers, they induce a twisting of the ODs by approximately 10°, which buries the SD2 and D-loops deeper into the filament’s core. This conformational change increases the twist between adjacent protomers (n, n + 2 subunits) along the same protofilament from approximately 26° in canonical F-actin to approximately 36° in cofilactin, corresponding to a change of the twist from −167° to −162° between adjacent protomers (n, n + 1 subunits) on opposite strands (Galkin et al., 2011; Tanaka et al., 2018). These collective protomer rotations, readily visualized in 2D filament projections, shorten the HHP and reduce the number of actin protomer pairs within each helical half-turn. Consequently, cooperative OD twisting drives the formation of the compact, supertwisted C-actin architecture (Ngo et al., 2024). In contrast, alpha-actinin crosslinking stabilizes the flattened F-actin protomer conformation, preserving the canonical helical twist and normal HHP while opposing the structural remodeling required for favorable binding of cofilin.

### A conserved actin-remodeling function of calponin homology domains

The present findings, together with previous work on Rng2-CHD (Hayakawa et al., 2022), suggest that calponin homology (CH) domains may possess a previously unrecognized and conserved function in remodeling the higher-order architecture of actin filaments. The ABD of human alpha-actinin-1 comprises two tandem CH domains (CH1-CH2), in which CH1 forms the primary actin-binding interface whereas CH2 regulates the accessibility and affinity of CH1 through intramolecular interactions (Borrego-Diaz et al., 2006). The E235K mutation is located within the CH2 domain and is therefore unlikely to directly alter the actin-binding surface. Instead, it is proposed to weaken the CH1-CH2 interface, favoring a more open ABD conformation that enhances the accessibility of CH1 and promotes structural remodeling of actin filaments beyond simply increasing binding affinity. This mechanism is derived from previous studies of the human alpha-actinin-4 ABD-K255E (Shams et al., 2016; Weins et al., 2007) and filamin ABD-E254K mutants (Iwamoto et al., 2018), which similarly disrupt the CH1-CH2 interface and increase actin-binding activity.

In this study, HS-AFM revealed that ABD-E235K and ABD-WT convert canonical right-handed double-helical actin filaments into two or three alternative architectures, the single-helical protofilament, parallel-helical protofilament state, and nano-ring structure. Remarkably, these structural transitions closely resemble those induced by the isolated calponin homology (CH) domain of the fission yeast cytokinetic protein Rng2, despite Rng2 containing only a single N-terminal CH domain rather than a tandem CH1-CH2 domains of alpha-actinin (Hayakawa et al., 2022). In the Rng2-CHD study, HS-AFM directly visualized the separation of the canonical double helix into individual protofilaments while demonstrating that each protofilament intrinsically retained its helical periodicity after dissociation, providing compelling evidence that the SP and PPS represent authentic actin conformations rather than imaging artifacts (**Figure S13**). The striking agreement between these independent observations strongly supports the existence of previously underappreciated structural polymorphs of actin filaments (i.e., SP, PPS, and nano-ring). More importantly, it raises the intriguing possibility that diverse CH-domain-containing proteins remodel actin filaments through a common structural mechanism despite their distinct CH domain architectures. Because tandem CH domains constitute the conserved ABD of numerous actin binding proteins, including alpha-actinin, spectrin, dystrophin, utrophin, and filamin (Korenbaum and Rivero, 2002), it is tempting to speculate that their biological functions extend beyond filament binding to include active regulation of actin filament architecture. Such remodeling may represent a general mechanism by which CH-domain proteins tune filament mechanics, cooperativity, and accessibility to other actin binding proteins during cytoskeletal organization. Testing whether this structural remodeling activity is conserved across the broader CH-domain protein family will therefore be an important direction for future studies and may redefine the functional repertoire of one of the most prevalent actin-binding modules in eukaryotic cells.

### CH-domain binding reveals alternative actin filament polymorphs

Our findings reveal an unexpected structural plasticity of actin filament that extends beyond the canonical right-handed double-helical filament. Transient binding of the alpha-actinin-1 ABD-E235K mutant and wild-type induced transitions from the canonical filament into two or three alternative conformations, which we term the SP, PPS, and nano-ring conformation. The SP and PPS structures were not associated with detectable changes in the intrinsic helical periodicity of individual protofilaments, their apparent height was reduced by more than half (**Figure 4-5**), suggesting that each protofilament intrinsically maintains its helical periodicity after dissociation from the intact double helix. The observation of the similar SP and PPS induced by the CH domain of Rng2 further supports that these conformations represent authentic actin filament polymorphs rather than imaging artifacts (**Figure S13**) (Hayakawa et al., 2022).

Strikingly, cofilin remained capable of recognizing and forming clusters on the PPS. However, unlike on canonical double-helical actin filaments, cofilin binding to these alternative architectures did not induce the characteristic shortening of the helices associated with PPS-cofilin complex segments (**Figure S14-S15**). These findings suggest that cofilin cluster formation and cofilin-induced structural remodeling are mechanistically separable processes on the PPS-like-helical pitch. Cofilin can bind to and form clusters along a single actin protofilament without inducing the characteristic helical-pitch shortening observed in canonical cofilactin.

We therefore propose that SP, PPS, and nano-ring conformation represent previously unrecognized structural states within the actin conformational landscape that can be selectively stabilized by transient interactions with CH domain-containing proteins. These findings expand the current view of actin filament as a structurally polymorphic biopolymer.

## CONCLUSION

In our current study, we integrated HS-AFM, MD simulations, and PCA to infer the atomic conformational dynamics of actin filaments and alpha-actinin. Our results demonstrate that high full-length alpha-actinin crosslinking densities stabilize flattened actin protomers and canonical helical geometry, thereby suppressing cooperative cofilin binding and cluster maturation while largely preserving S1 binding. We further show that cofilin assembly on canonical double-helical actin filament requires an initial preference for shortened helical conformations, followed by protomer twisting and helical shortening that drive cluster expansion. In contrast, transient bindings of human alpha-actinin-1 ABD-E235K and ABD-WT remodel canonical double-helical actin filaments into previously unrecognized single-helical protofilament, parallel-helical protofilament state, and nano-ring architectures. Notably, PPS retains a nearly unchanged helical periodicity, and these ABD-induced structural rearrangements only weakly inhibit cofilin cluster formation. The similarity of these structures to those induced by the Rng2 CH domain suggests a conserved actin-remodeling function of CH-domain proteins. Collectively, our findings uncover a molecular- to atomic-level mechanism by which alpha-actinin regulates actin filament architecture and reveal an unexpected structural polymorphism of actin filament that provides an additional layer of regulation for actin binding protein interactions and cytoskeletal.

## MATERIALS AND METHODS

### Proteins and lipids

Rabbit skeletal actin and human cofilin-1 were prepared following the same protocol as in our previous study (Ngo et al., 2015). Recombinant human alpha-actinin-1 ABD-E235K and ABD-WT carrying an N-terminal His tag was expressed in *E. coli* and purified by Ni-NTA affinity chromatography. Chicken gizzard smooth muscle alpha-actinin-2 (Ribeiro et al., 2014) was obtained from Cytoskeleton. Subfragment-1 (S1) was prepared from smooth muscle myosin using the method described by Asakawa and Azuma (Asakawa and Azuma, 1988). The lipid compositions used in this study consisted of neutral 1,2-dipalmitoyl-sn-glycero-3-phosphocholine (DPPC, Avanti) and positively charged 1,2-dipalmitoyl-3-trimethylammonium-propane (DPTAP, Avanti). Liposomes were prepared with a DPPC/DPTAP ratio of 90:10 (wt%), and the lipid membrane was subsequently formed on mica (Ngo et al., 2015). In all experiments, actin filaments were similarly immobilized on DPPC/DPTAP ratio of 90:10 (wt%) for HS-AFM imaging, as done in our previous study (Ngo et al., 2015).

### HS-AFM Analysis

AFM experiments were performed using a custom-built high-speed atomic force microscope (HS-AFM) developed by Ando and colleagues (Ando et al., 2024). HS-AFM operated in intermittent tapping mode with amplitude modulation feedback in solution, using ultra-short cantilevers (USC-F1.2-K0.15; NANOANDMORE) with a nominal spring constant of approximately 0.15 N/m, a resonant frequency of approximately 1.2 MHz, and a quality factor of approximately 3.0 in air. HS-AFM imaging was carried out essentially as described previously (Le et al., 2025; Ngo et al., 2024, 2022) and the detailed protocols (Uchihashi et al., 2012). For HS-AFM imaging, the peak-to-peak free oscillation amplitude (A₀) of the cantilever was set to approximately 1.6–1.8 nm, and the feedback setpoint was adjusted to approximately 0.9 A₀. AFM image rendering and data processing were performed using custom-made analysis software UMEX Viewer and Kodec software, as described previously (Ngo et al., 2024, 2015).

### HS-AFM imaging of actin filaments

We employed HS-AFM to capture the dynamic conformations of actin filament-crosslinked with alpha-actinin under various conditions, providing high-speed imaging and 3D surface structure of the protein at nanometer resolution. Alpha-actinin was diluted to a desired concentration in F-buffer containing 20 mM KCl, 10 mM PIPES-KOH (pH 6.8), 0.5 mM MgCl_2_, 0.25 mM EGTA, 0.25 mM DTT, and 0.5 mM ATP, as similarly done (Ngo et al., 2026). We used F-buffer at pH 6.8 to study cooperative cofilin binding and cluster formation on actin filaments while minimizing cofilin-induced severing and depolymerization, which are known to increase at higher pH (∼7.8) (Wioland et al., 2019, 2017). All experimental conditions were generally replicated three times or more for reproducibility.

To investigate the dynamic conformational changes in actin filaments crosslinked solely by alpha-actinin, actin filaments were gently immobilized on a lipid membrane and similar F-buffer consisting of 0.5 mM MgCl₂, 0.5 mM ATP, and 50 nM alpha-actinin, but without CaCl₂, was employed for HS-AFM imaging.

We attempted to produce actin filaments crosslinked with alpha-actinin in both unipolar and bipolar configurations, as reported previously (Meyer and Aebi, 1990; Taylor et al., 2000), for HS-AFM imaging. However, consistent with observations of Taylor et al. (Taylor et al., 2000) using smooth muscle alpha-actinin on 2D lipid surface, our AFM images revealed only unipolar crosslinking on lipid membranes. This contrasts with the major bipolar arrangements reported by Meyer and Aebi (Meyer and Aebi, 1990). At present, we do not have a definitive explanation for this observation. Meyer and Aebi reported that alpha-actinin from chicken gizzard and *Dictyostelium* predominantly promotes antiparallel (bipolar) crosslinking, whereas *Acanthamoeba* alpha-actinin favors parallel (unipolar) arrangements. However, Taylor et al. proposed that the formation of strictly bipolar or unipolar arrays may require additional factors that either determine filament orientation or modulate the crosslinking activity of alpha-actinin. These include the β-integrins, vinculin, intercellular adhesion molecule (ICAM), titin, zyxin, and nebulin. These proteins, which bind to different domains on alpha-actinin, may affect its interaction with actin in ways not yet understood (Taylor et al., 2000).

To study the dynamic conformational changes in actin filaments crosslinked by alpha-actinin in the presence of myosin II heads (S1), actin filaments were gently immobilized on a lipid membrane and immersed in F-buffer containing 1 mM ATP, 0.1 mM CaCl₂, 50 nM alpha-actinin, and 1 µM S1, but without MgCl₂, for HS-AFM imaging.

To study the competitive interactions between cofilin and S1, which utilize distinct short and normal helical structures on the same actin filament to form their respective clusters, actin filaments were gently immobilized on a lipid membrane. HS-AFM imaging was performed in F-buffer initially containing 1 mM ADP, 0.01 mM ATP, and 300 nM S1, followed by the addition of 100 nM cofilin.

To explore the binding and supertwisting activities of cofilin clusters on the half helices of actin filaments without and with alpha-actinin crosslinking, actin filaments were gently immobilized on a lipid membrane, either without or with 400 nM alpha-actinin crosslinking in F-buffer containing 1 mM ATP, followed by the addition of 600 nM cofilin, for HS-AFM imaging.

To investigate the transient binding of the human alpha-actinin-1 ABD-E235K mutant or ABD-WT to actin filaments in the presence and absence of cofilin, two complementary HS-AFM experiments were performed.

In the first set, actin filaments were gently immobilized on positively charged lipid membranes (10% DPTAP), and ABD-E235K or ABD-WT were added during HS-AFM imaging to directly visualize its transient interactions with the filaments. Cofilin was subsequently introduced into the same imaging chamber without buffer exchange, and its binding and cluster formation were monitored in the same F-buffer (pH 6.8) containing 1 mM ATP. When required, the buffer was manually exchanged to remove unbound ABD-E235K, allowing cofilin cluster formation to be assessed in the presence or absence of ABD-E235K. By doing so, keeping the same image position before and after buffer exchange was not technically practical at this moment. Protein concentrations were provided in the legends of figures or Videos. This condition was also designed to successfully capture ABD-WT-induced formation of the PPS and nano-ring in actin filaments. In the second set, designed to evaluate ABD-E235K-induced formation of the single-helical protofilament and parallel-helical protofilament state, actin filaments were pre-incubated with ABD-E235K or ABD-WT for approximately 30 min at room temperature before immobilization on the lipid membranes for HS-AFM imaging. Cofilin was then added to examine its binding behavior and cluster formation on these actin filaments. Protein concentrations were indicated in the legends of figures or Videos. All HS-AFM experiments were carried out at room temperature and replicated three times or more for reproducibility.

### Quantitative analysis of half helical pitch and protomer pairs

The half helical pitch (HHP), the number of protomer pairs per HHP, and the mean axial distance (MAD) between adjacent actin protomers along the same long-pitch strand within individual HHPs were analyzed for actin filaments crosslinked by alpha-actinin under conditions containing Ca^2+^, ATP, and S1, but without MgCl_2_ were analyzed using HS-AFM. The semi-automated analytical methods used in this study were adapted from our previous research (Ngo et al., 2024).

To quantify HHPs and the number of protomer pairs per HHP, we used Umex Viewer for Drift Analysis to semiautomatically determine the distances between the highest crossover points of the two long-pitch strands. Topographical line profiles were generated along the actin filaments. Each HHP was defined as the distance between two consecutive highest crossover points, whereas one helical pitch (HP) was measured between three consecutive crossover points.

Briefly, longitudinal profiles spanning 1–3 consecutive HHPs were generated along the filaments, with height offsets set to zero relative to the lowest position along each profile. Before analysis, XY-piezo nonlinearity was corrected by nonlinear image scaling, and image noise was reduced using a Gaussian smoothing filter (σ = 0.76 nm). Profiles were obtained by averaging signals within a 5–8 nm-wide band along the filament. To ensure accurate detection of the axial distance (AD) between adjacent protomers within the same long-pitch strand, the minor-pitch detection range was set to 3–6 nm for detecting AD. The minimum major-pitch range was set to 22–25 nm for detecting HHP for bare actin and alpha-actinin-crosslinked actin segments. These facilitate accurate AD and HHP detection and penalize incorrectly identified peaks. In rare cases, missing peaks were manually added and noise-induced false peaks removed by an analyzer (**Figure S2**).

Typically, 10–15 actin filaments were analyzed per condition, with HHPs measured over approximately 20 consecutive frames. To minimize measurement errors in AD, HHPs, and protomer-pair counts, we selected straight filament segments with clearly distinguishable protomers and analyzed profiles with an average width of 3%, covering 1–3 consecutive HHPs. The same procedure was consistently applied to bare actin and alpha-actinin-crosslinked actin filaments.

To calculate the number of protomer pairs per HHP and the MAD between adjacent protomers along the same long-pitch strand, AD values were semiautomatically detected and protomer pairs were counted, as described above and illustrated in Figure S2. The numbers of actin protomer pairs per HHP (as counted major and small peaks) within each HHP subtracted by 0.5 (Ngo et al., 2024). The MAD within each HHP was calculated by dividing the HHP by the corresponding number of protomer pairs.

To enhance the characteristics of actin protomers in the different types of filaments, we converted the raw AFM images into Laplacian-filtered images for a cross-checking of protomer counting when needed. This was achieved by applying a high-pass filter known as the Laplacian of Gaussian (LoG) filter, which can be found at the following link: (https://homepages.inf.ed.ac.uk/rbf/HIPR2/log.htm). The Laplacian filter used in this process is a type of high-pass filter, characterized by the kernel 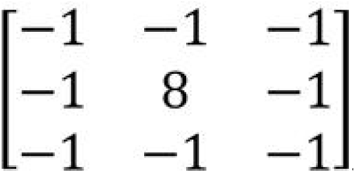. Prior to the application of the Laplacian filter, we employed a Gaussian filter with a typical sigma value of 1.27 pixel/nm. This preliminary step helped reduce sensitivity to noise and enhanced the resulting Laplacian-filtered images, resulting in improved visualization of the actin protomers (**Figure S2**).

### Methods for measuring alpha-actinin crosslinking angles and crosslinking length between filaments

Alpha-actinin molecules crosslink two actin filaments in either a parallel (unipolar) or anti-parallel (bipolar) configuration. In this study, we analyzed the crosslinking angles (a small angle) relative to the filament axis and crosslinking length between actin filaments based on unipolar crosslinking conformations obtained from HS-AFM. Two to four alpha-actinin molecules consistently intercalated between actin filaments at every half-helical turn, forming a parallelogram-like configuration.

To analyze the conformations derived from HS-AFM, we tracked the center of mass (CoM) positions of four crossover points on actin filaments (e.g., A, B, C, D or E, F, G, H as illustrated in **Figure S9**) in a parallelogram, using their vertex coordinates. This process was performed with the Kodec software to semi-automatically detect the CoM positions (Kodec including its source code, (Ngo et al., 2015)).

To compute the interior angles of a parallelogram when the vertex coordinates were known, we followed these steps:

Given four vertices—A (x1, y1), B (x2, y2), C (x3, y3), and D (x4, y4)—arranged sequentially around the parallelogram, the first step was to determine the four vectors representing its sides:

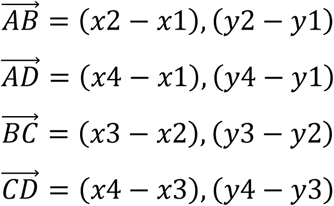

Second, the angle between two vectors were calculated as:

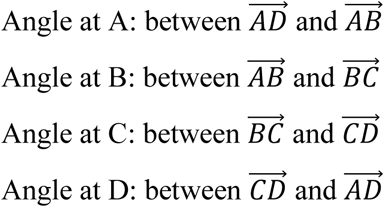

For example, an angle *θ* (rad) at A:

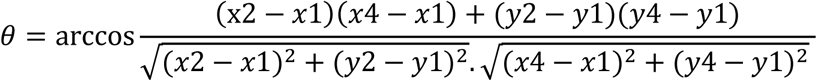

The angle (degrees) at A was then calculated:

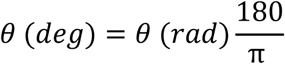

The angles at B, C, and D were calculated using the same approach. Finally, the physical lengths of AB, BC, CD, and DA segments were similarly determined and converted the lengths from pixel to nm unit. We note that the crosslinking length showed by these measurements is longer than that measured in MD simulations, and it is explained below.

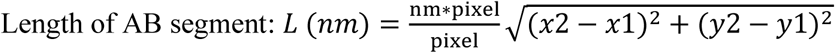

The approach was similarly applied to analyze the crosslinking angles in unipolar and bipolar crosslinking conformations obtained from MD simulations. However, the crosslinking lengths measured from MD simulations were shorter than those obtained from AFM images because the MD analysis defined and measured the direct perpendicular distance between the two filaments, whereas the AFM-based analysis defined and measured the corresponding side-to-side distances of the parallelogram-like configuration (see AD and BC or EG and FH lines in Figure S4).

### Challenging in flexible fitting of alpha-actinin-crosslinked actin filaments captured by HS-AFM

Flexible fitting of alpha-actinin molecules bound to actin filaments imaged by HS-AFM presents several challenges, primarily because their molecular architecture is partially obscured by the actin filaments. In particular, the two actin-binding domains (ABDs) of the alpha-actinin homodimer are concealed upon crosslinking, hindering the reconstruction of the complete molecular structure and the accurate determination of its conformational states (**Figure 1**). Nevertheless, the visible portions of alpha-actinin exhibit reproducible structural dynamics, including bending and twisting motions within the homodimer ABD (Ngo et al., 2026).

Consequently, we were unable to achieve reliable flexible fitting of the MD-generated conformations of alpha-actinin-crosslinked actin filaments to the experimental AFM images. The challenge was also like flexible fitting of actin filaments.

### All-atom MD simulations

For the unipolar and bipolar systems, two actin filaments crosslinked by an alpha-actinin, the initial coordinates were originally given in the Liu and Taylor’ paper (Liu et al., 2004). The link in the paper to download the structures is not available at the present, so a corresponding author, Kenneth A. Taylor, kindly provided them to us. The models of alpha-actinin without Ca^2+^ were similarly constructed in our previous study (Ngo et al., 2026). In the previous calculation using QM/MM (Kanematsu et al., 2022), water coordination around Mg^2+^ and ADP in the actin protomer was investigated. We selected ADP-bound actin protomers to construct the initial model because most actin filaments used in our HS-AFM experiments were aged after polymerization, allowing ATP hydrolysis and subsequent inorganic phosphate (Pi) release to generate ADP-actin filament (Kudryashov and Reisler, 2013). First, a protomer (PDB code: 8d15) (Reynolds et al., 2022) was solvated in a 20 mM NaCl solution with one Mg^2+^ and one ADP. The system consists of one protomer, 73 Na^+^, 63 Cl^−^, 1 Mg^2+^, one ADP, and 23,442 water molecules. All histidine residues were protonated at the epsilon carbon. For the protein, water molecules, monovalent ions, and divalent ions, the same force fields as in the case of alpha-actinin simulations were used: proteins, AMBER ff19SB (Tian et al., 2020); water molecules, TIP3P (Jorgensen et al., 1983); monovalent ions, the Li and Merz 12-6 model (Li et al., 2015); divalent ions, the Li and Merz 12-6-4 model (Li and Merz, 2014). The Carlson model was employed for ADP (Meagher et al., 2003). The system was equilibrated by 100 ps MD simulation with constant temperature (300 K) and pressure conditions (1 bar) using the Berendsen thermostat and barostat (Berendsen et al., 1984). The length of bonds having H atoms was kept constant using the SHAKE algorithm, enabling the use of time step of 2 fs. Long-range interactions were calculated by the particle mesh Ewald method with a 10 Å cutoff in real space. The actin protomer, ADP, and Mg^2+^ were constrained with a force constant of 10 kcal/mol/Å^2^ to their original positions. Then, 3 μs MD simulation with constant temperature (300 K) and volume conditions was performed using the Berendsen thermostat (Berendsen et al., 1984). The actin protomer, ADP, and Mg^2+^ were weakly constrained with a force constant of 0.2 kcal/mol/Å^2^ to their original positions to prevent unwanted large structural change in the bulk solution. Water molecules gradually entered the protomer over 2 μs. The coordination of water molecules around Mg^2+^ and ADP was confirmed to be similar to the previous configuration obtained by QM/MM calculation (Kanematsu et al., 2022). In the remaining 1 μs simulation, the number of water molecules around Mg^2+^ has fluctuated but did not change. The protomer, ADP, Mg^2+^, and 27 water molecules around Mg^2+^ were picked up and aligned to the Liu and Taylor’ configuration by minimizing root-mean-square deviation (RMSD) between Cα atoms (Liu et al., 2004). Then, protomers were elongated while considering the symmetry to form 36 nm filament. The periodic boundary condition was imposed on the filament in a vacuum, and the energy was minimized to remove bad contacts. In the Liu and Taylor’ configuration, a part of the neck domain in the alpha-actinin (residue 251-260) was missing, which was complemented by the Modeller package (Sali and Blundell, 1993). The system consisting of two actin filaments and alpha-actinin was then solvated in a 40 mM KCl solution. The final system of unipolar configuration consists of two actin filaments (26 protomers), one alpha-actinin, 1032 K^+^, 718 Cl^−^, 26 Mg^2+^, 26 ADP, and 1,433,011 water molecules, and total number of atoms in the system was 4,479,739. The system was shortly optimized with 1000 step minimization and was equilibrated by 130 ps MD simulation with constant temperature (300 K) and pressure condition (1 bar). The temperature gradually increased up to 300 K. The pressure was anisotropically controlled using Monte Carlo barostat so that only the box size in z direction was allowed to move, in which the actin filament was placed along the x direction. During the equilibration, the actin filament, the alpha-actinin, Mg^2+^, ADP, and water molecules inside the actin were constrained with a force constant of 2 kcal/mol/Å^2^. The size of the equilibrated system was 360.000 × 480.809 × 260.784 Å^3^. Then, the constraint was turned off and only the box size along the x direction was allowed to move to observe all-atom fluctuation including twist, rise, HHP, and AD values. All-atom MD simulations were performed for a total time of 600 ns for unipolar-flexible and bipolar-flexible systems using three replicas each, and for 100 ns bipolar-fixed systems. In unipolar-flexible and bipolar-flexible models, all atoms were allowed to freely move during simulations. In contrast, in the bipolar-fixed model, all Cα atoms of alpha-actinin were restrained with a force constant of 40 kcal/mol/Å^2^, suppressing their motions. In the simulations, periodic boundary conditions were applied, causing one end of the filament to connect to the other. As a result, 13 protomers form a single unit cell, corresponding to one HHP in this study, and this number remains fixed throughout the simulations. The trajectories saved at each 10 ps, and 10,000 snapshots from the last 100 ns from each replica were used for the analyses. In the same way, the system was built and the simulation was performed for the bipolar configuration, but the number of water molecules was 1,432,853. To build a bare actin filament system, the alpha-actinin was removed from the unipolar system, and 100 ps MD simulation with constant temperature (300 K) and pressure conditions (1 bar) was performed to fill the space originally located by the alpha-actinin. MD simulations were similarly performed for a total time of 500 ns using three replicas. The time evolutions of the SD2-SD4 distance and potential energy of all systems were indicated in Figure S26-S27. Also, RMSD of the protomers in all systems were shown in Figure S25.

### Principal component analysis

After performing MD simulations, we conducted the analyses of conformations of alpha-actinin and actin protomers obtained from unipolar, bipolar models, and bare filament using PCA. This approach allowed us to capture their atomic conformational dynamics, enabling an evaluation of binding and the physical motion effects such as bending, twisting, and rotation of alpha-actinin on fluctuations of the twist, rise, AD, dihedral angles, SD2-SD4 distance in the crosslinked actin filaments for a comparison with that in bare actin filament. For alpha-actinin, before the PCA, the configuration of alpha-actinin was aligned with respect to the center part of the rod domain (residue 485-525). PCA was applied by diagonalizing the covariance matrix (*C*), defined as follows

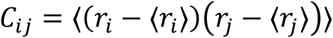

where *r_i_* represents the x, y, and z positions of the *i*th Cα atom and angular brackets represent the time average.

For actin protomers isolated from bare actin, unipolar, and bipolar models, protomers were aligned in the same direction by minimizing RMSD of Cα atoms from a protomer in the 8D15 pdb structure. We calculated PCA separately and obtained one conformational space for each MD simulation (bare actin, unipolar and bipolar).

### Quantifying actin protomer conformations

The dihedral angle of actin protomers, representing the rotational orientations between SD2-SD1-SD3-SD4 or between OD: SD1-SD2 and ID: SD3-SD4 within individual actin subunits, and SD2-SD4 distances (**Figure S26**) were calculated from all-atom MD simulations over 300 ns for bare actin filaments, unipolar, and bipolar filament models. These calculations (Iyer et al., 2025) were based on the centers of mass of the Cα atoms within each subdomain, defined as follows: subdomain 1 (SD1), residues 5-32, 70-144, and 338-375; subdomain 2 (SD2), residues 33-69; subdomain 3 (SD3), residues 145-180 and 270-337; and subdomain 4 (SD4), residues 181-269. Across these analyses, actin protomers were categorized into either flattened or twisted conformations, which are critical for deciphering the actin structure-dependent cooperative cofilin binding and cluster formation.

### Calculation of two-dimensional free energy surface

We calculated free energy surface for the joint probability distribution of SD2-SD4 distance (*d*) and dihedral angle (*θ*) or rotational orientation between SD2-SD1-SD3-SD4 or ID and OD within the actin protomer (**Figure S26-S27**). The free energy surface (*F*(*d, θ*)) was derived from the probability density (*P*(*d, θ*)), which is the normalized counts at *d*, and *θ*:

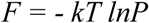

where *k* is the Boltzmann constant and *T* is the absolute temperature.

Our all-atom MD simulations represent standard equilibrium trajectories (300 ns) with three 100 ns replicas, allowing access only to conformations within the lower-energy region (< 5 kcal/mol) of the free energy landscape. Despite this limited sampling range, the stable conformations identified are consistent with those reported by Iyer and Voth et al. for the ADP-actin filament protomer, (Iyer et al., 2025) confirming the robustness of the observed structural states.

### From local protomer twisting to global helical shortening

In our previous study (referred to **Figure 2** in *Ngo et al* (Ngo et al., 2024)), the geometric basis of cofilin-induced helical shortening can be understood by approximating the actin filament as a helical lattice wrapped around an imaginary cylinder. When the cylindrical surface is mathematically “unwrapped” into a 2D plane, the helical trajectory becomes a diagonal line whose slope depends on the rotational angle between adjacent protomers within the same strand.

For a canonical double-helical actin filament, the rotation increment between adjacent protomers within the same strand is approximately:

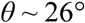

with an axial rise per protomer (**Figure 1**, this study):

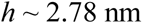

The total angular rotation accumulated after *N* protomers is:

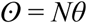

A half helical pitch (*HHP*) corresponds to a rotation of approximately:

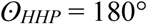

For a canonical actin filament, the number of protomer pairs (*N\**) per *HHP*:

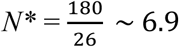

which experimentally corresponds to approximately 6.5 protomer pairs (N∼13 subunits per HHP) and a length of an *HHP* of:

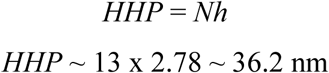

This value is consistent with the experimental canonical *HHP* (∼ 36 nm).

Upon cooperative cofilin binding and mature cluster formation, the outer domains (ODs) of actin protomers undergo coordinated twisting of ∼10° (Galkin et al., 2011), increasing the rotational increment between adjacent protomers to approximately:

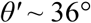

The number of protomer pairs (*N’*) required to achieve the same 180° half turn becomes:

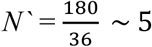

which corresponds to *N^#^* =10 actin protomers per *HHP’*. The newly shortened *HHP’* = *N^#^h* becomes:

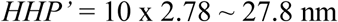

which well agrees with experimentally observed shortened helices of canonical cofilactin *HHP’* of ∼27.9 ± 3.0 nm (Figure 2D) in this study.

Therefore, increasing the rotation angle from ∼26° to ∼36° reduces the number of protomers required to complete a half-turn (180°), resulting in global shortening of the HHP. The unwrapped-cylinder model illustrates this relationship, with larger rotational increments producing steeper helical trajectories and shorter pitches. Thus, the characteristic shortening of cofilactin arises directly from the collective local twisting of actin protomers induced by cooperative cofilin cluster formation (Ngo et al., 2024).

## Acknowledgments

The authors express their gratitude for the facility and financial support provided by the WPI Nano Life Science Institute (WPI NanoLSI), Kanazawa University; the instrumental assistance provided by Toshio Ando, Noriyuki Kodera, and Hiroki Konno (WPI NanoLSI); and the initial bipolar and unipolar models presented by Kenneth A. Taylor (Florida State University); and initial AFM data analysis by Thao Phuong Ngo and Thien Quang Huynh (WPI-NanoLSI). The authors also acknowledge the financial support from KAKENHI (Japan Society for the Promotion of Science) for K.X.N. (19K06581, 23K05713, 23H02452), T.S. (24K01308), and T.Q.P.U. (23H02452). The MD simulations were carried out on the supercomputers at the Research Center for Computational Science in Okazaki, Japan (Project: 26-IMS-090). This study was also partially supported by the French National Research Agency (research grant ANR-23-CE45-0024-01) and within the framework of the “Investissements d’avenir” program (ANR-15-IDEX-02) for S.G.

## Author contributions

K.X.N. designed the research; K.X.N. and T.S. performed the research; K.X.N., T.S., T.Q.P.U., N.T.P.L., H.G.N., R.V., and S.G. analyzed the data; K.X.N. and T.S. wrote and edited the paper. All authors discussed and revised the paper.

## Competing interests

The authors declare that they have no competing interests.

## Data, code, and materials availability

All data and code needed to evaluate and reproduce the results in the paper are present in the paper and/or the Supplementary Materials. This study did not generate new materials.

---

**Figure S1.**
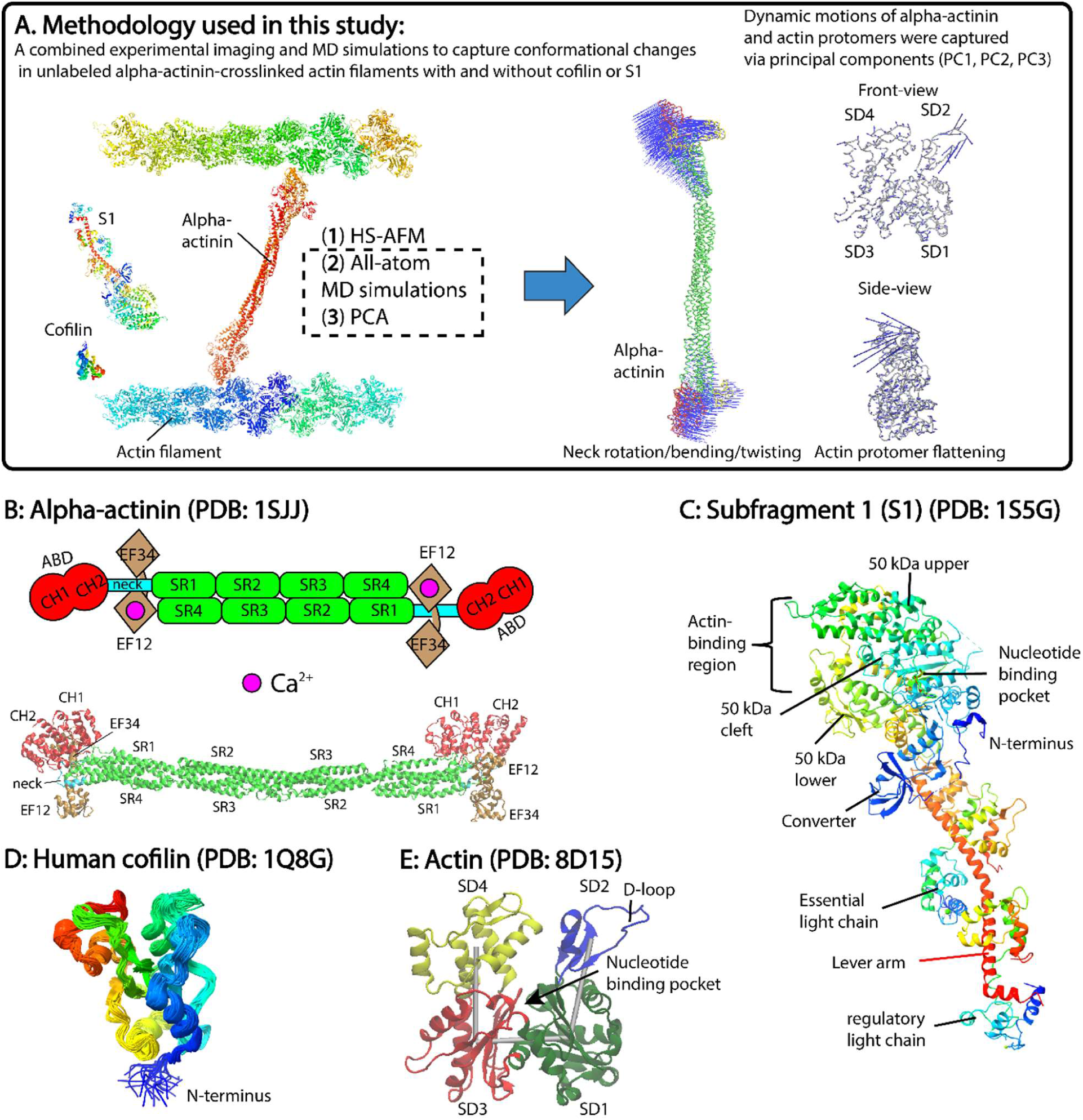
Methods and proteins used in this study. **(A)** The combined approach provides insights into the atomic structural changes and dynamic behavior of actin filaments upon interaction with the actin binding proteins including alpha-actinin, S1, or cofilin. **(B)** Domain structure of chicken gizzard smooth muscle alpha-actinin (PDB: 1SJJ) (Liu et al., 2004; Ngo et al., 2024) shown in ribbon. Alpha-actinin is an antiparallel homodimer (>200 kDa) with an N-terminal actin binding domain (ABD), in which the two calponin homology 1 and 2 (CH1, CH2) domains are in extensive contact, a central region with four spectrin-like repeats (SR1–SR4), and a C-terminal calmodulin-like domain (CaMD) containing two EF-hand motifs (EF12 and EF34), with EF12 responsible for Ca²⁺ binding. (**C, D, E**) Brief descriptions of domain structures and/or ribbon models of myosin II subfragment 1 (S1), human cofilin 1, and actin monomer (Pope et al., 2004; Reynolds et al., 2022; Risal et al., 2004). SD1, SD2, SD3, and SD4 denote subdomains 1-4.

**Figure S2.**
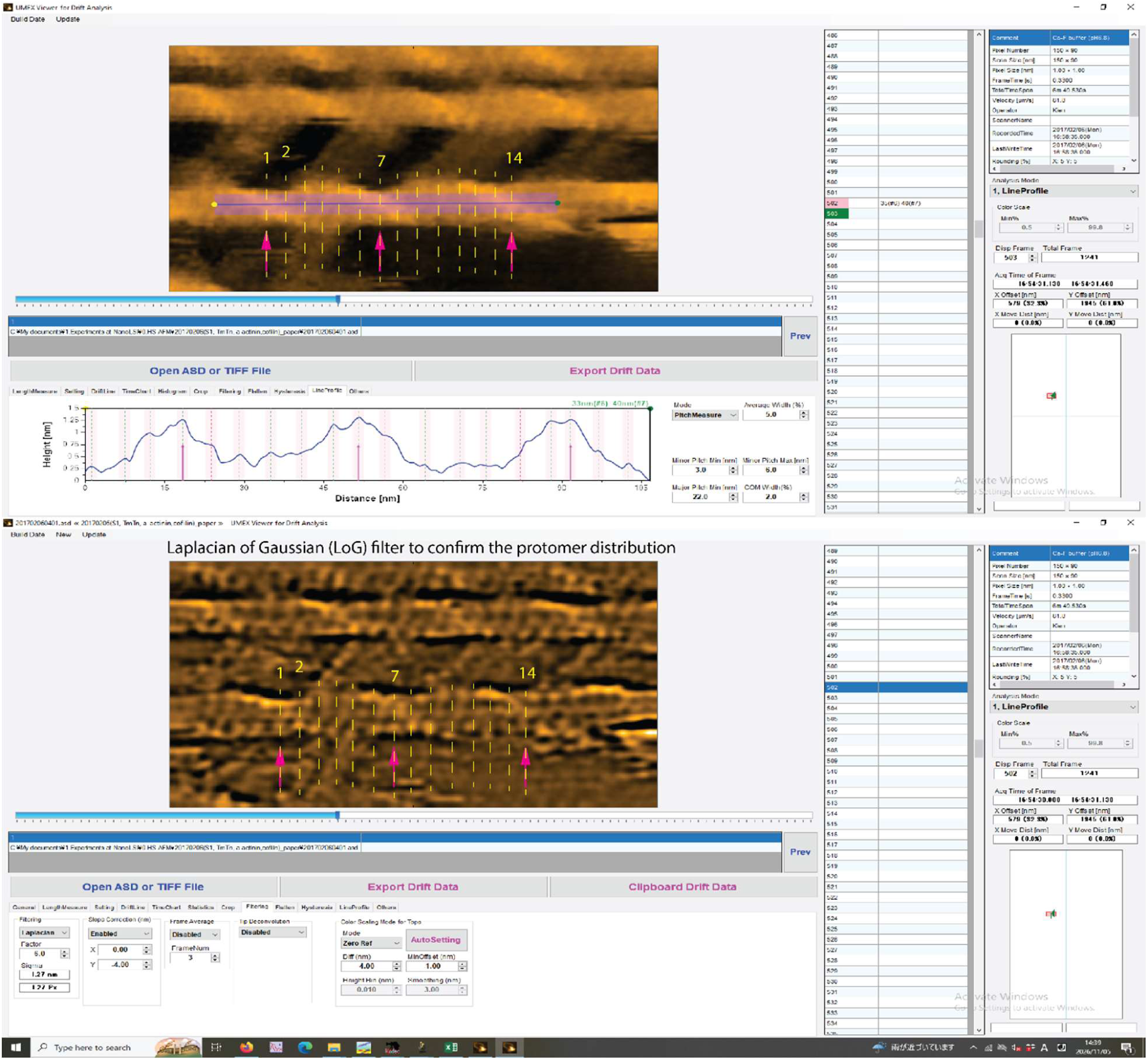
Demonstration of a semi-automatic analysis method of measuring half helical pitch (HHP) and counting the number of protomer pairs per HHP (see further detail in the Methods). Still AFM images were similarly reused in Figure 1B-C, S3, S4B.

**Figure S3.**
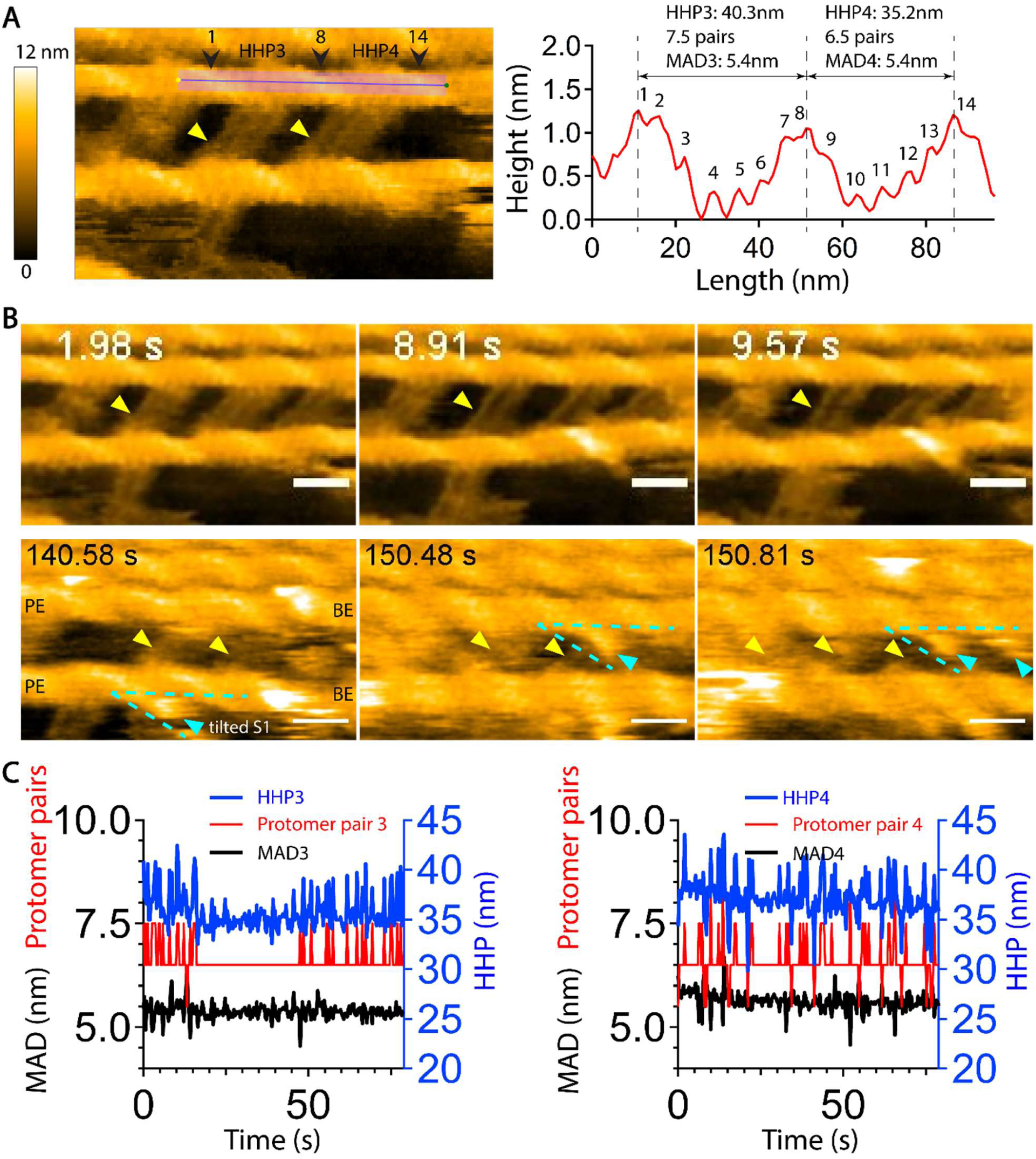
Analysis of the dynamic conformations of actin filaments crosslinked by alpha-actinin in unipolar configuration in the presence of Ca^2+^, ATP, and S1 using HS-AFM. **(A)** A longitudinal section profile illustrating the analysis of two consecutive HHPs, the number of protomer pairs per HHP, and the mean axial distance (MAD) between two adjacent protomers within the same-long pitch strand of actin filament. The height reference was defined along the actin filament. The analytic methods used were developed in our previous study (Ngo et al., 2024). **(B)** Still AFM images with time labels were shown for conditions with ATP present and after ATP depletion to observe the binding of S1 on actin filaments. As ATP levels gradually declined or nearly depleted due to ATP hydrolysis (ATP → ADP + Pi) during the actin-myosin reaction, S1 binding to actin filaments persisted for longer durations, enabling capture by HS-AFM at an imaging rate of 330 ms per frame. All actin filaments in the imaging field were aligned with the pointed end (PE) on the left and the barbed end (BE) on the right, reflecting the actual unipolar crosslinking configuration. As similarly done in our previous study (Ngo et al., 2024), the polarity of actin filaments was confirmed by observing the tilted angles of S1, with its head pointing toward the PE, resembling an arrowhead along the filament. This characteristic orientation allowed differentiation between the PE and the BE. Actin filaments crosslinked by 50 nM alpha-actinin in the presence of 1 mM ATP, 0.1 mM CaCl₂, and 1 µM S1, but without MgCl_2_. Note that two alpha-actinin homodimers may cooperate to crosslink every half-helix of actin filaments in this case, with their orientations varying due to molecular rotation. The effects of high versus low alpha-actinin crosslinking densities on the inhibition of cofilin binding and cooperative cluster formation are discussed separately. Representative yellow and cyan arrowheads indicate alpha-actinin and S1 molecules, respectively. Scale bars: 25 nm. Color scale: 0-12 nm. Still AFM images in A and B were similarly reused in Figure 1B-C, S2, S4B. **(C)** Time-dependent changes in two consecutive HHPs, the number of protomer pairs per HHP, and MADs in actin filaments crosslinked by 50 nM alpha-actinin in the presence of 1 mM ATP, 0.1 mM CaCl₂, and 1 µM S1, but without MgCl_2_.

**Figure S4.**
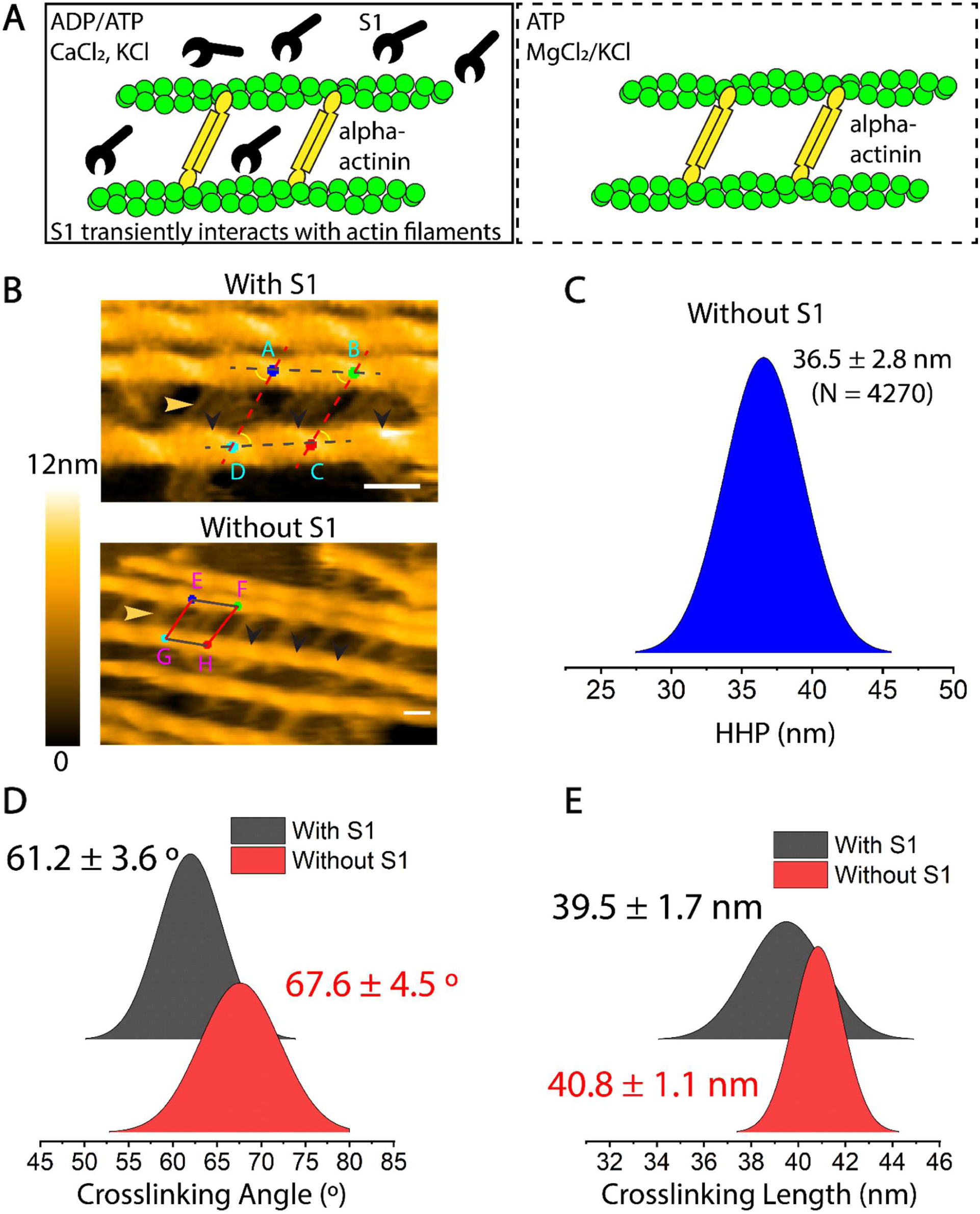
Quantitative analysis of crosslinking lengths, HHP, and alpha-actinin crosslinking angles relative to the filament axis, based on HS-AFM data of alpha-actinin-crosslinked actin filaments in the presence and absence of S1. **(A)** Experimental setup for HS-AFM observations of dynamic conformational changes in alpha-actinin-crosslinked actin filaments in the presence and absence of S1, including HHP, crosslinking angles, and crosslinking length. Actin filaments crosslinked by 50 nM alpha-actinin in the presence of 1 mM ATP, 0.1 mM CaCl₂, 20 mM KCl, and 1 µM S1, but without MgCl_2_ (right). Actin filaments were crosslinked with 50 nM alpha-actinin without S1 and observed in F-buffer supplemented with 0.5 mM MgCl₂, 20 mM KCl, and 0.5 mM ATP, but without CaCl_2_ at pH 6.8. (left). (**B**) Representative AFM images of alpha-actinin-crosslinked actin filaments with and without S1. Yellow arrowheads indicate alpha-actinin molecules; black arrowheads mark the crossover points of double-helical actin filament used to measure the HHP. Two parallelograms (ABCD and EFHG) represent the geometric framework used to measure the crosslinking angles of alpha-actinin and the crosslinking lengths between actin filaments. Points A-D and E-H mark actin crossover points used to derive the corresponding crosslinking lengths and angles. Still AFM images in A and B were similarly reused in Figure 1B-C, S2, S3A-B. (**C**) Histogram of HHP values for alpha-actinin-crosslinked actin filaments without S1. The mean HHP is 36.5 ± 2.8 nm (mean ± SD, N = 4270). (**D, E**) Ridgelines showing the distributions of alpha-actinin crosslinking angles and crosslinking lengths between filaments crosslinked with alpha-actinin, respectively. Note that we focus solely on the four symmetric small crosslinking angles generated by two alpha-actinin molecules bridging two actin filaments, irrespective of their polarity and demonstrated in a parallelogram ABCD. Statistical significance was determined using a two-sample *t-test*, with differences considered significant at *p* ≤ 0.05. See Methods and **Table 2** for details.

**Figure S5.**
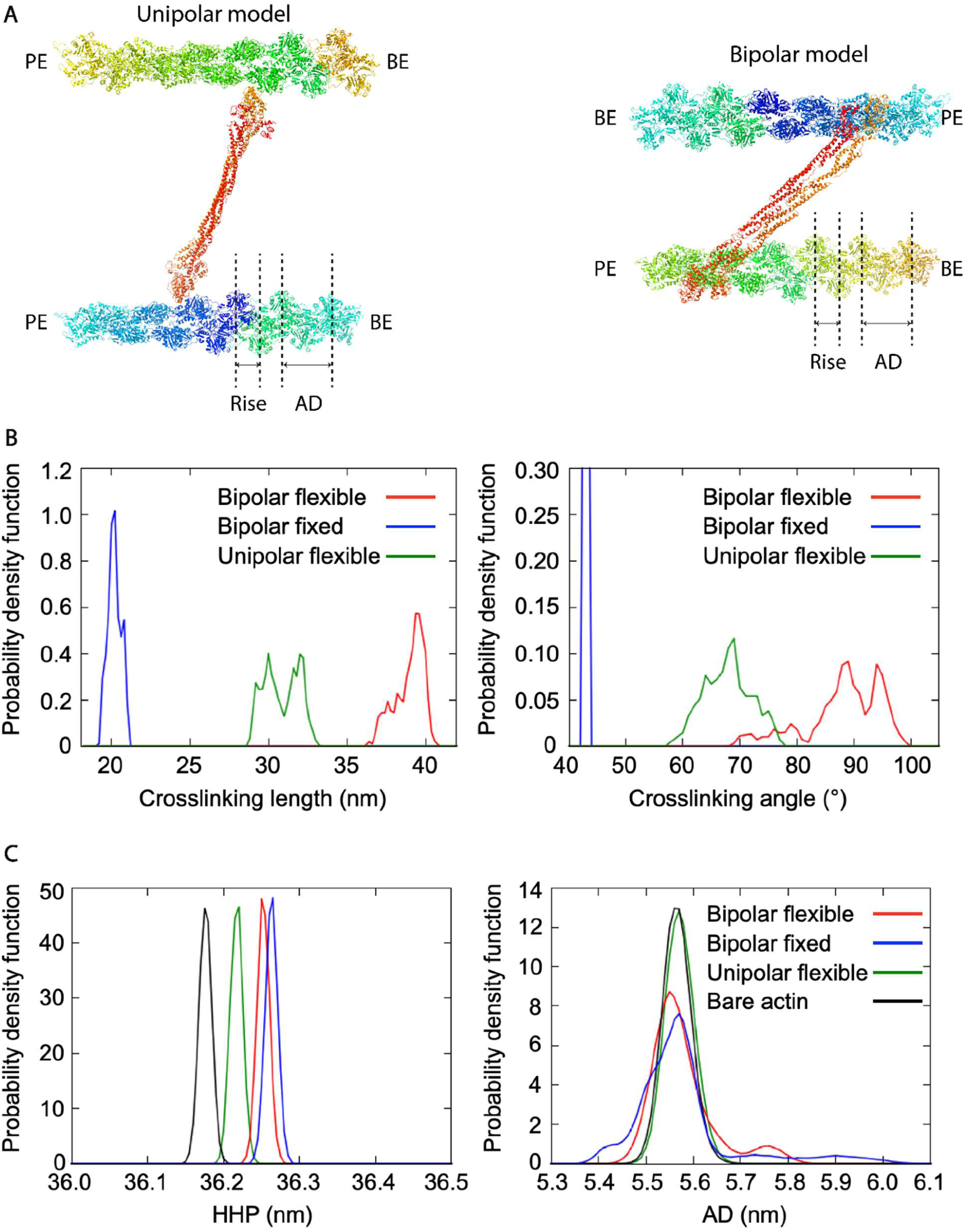
Differences in the conformational changes in the actin filaments crosslinked with alpha-actinin in unipolar and bipolar models derived from all-atom MD simulations. **(A)** Illustrations of the unipolar and bipolar crosslinking models of the actin filaments with alpha-actinin. The unipolar crosslinking configuration occurs when the actin filaments share the same polarity, whereas the bipolar crosslinking configuration appears when the actin filaments align with opposite polarities. The quantitative analyses were performed using conformations obtained from MD simulations for 300 ns with 3 replicas. PE: pointed end; BE: barbed end. (**B**) Quantitative analysis of the crosslinking lengths and alpha-actin crosslinking angles relative to the filament axis in the unipolar-flexible, bipolar-flexible, and bipolar-fixed models obtained from the MD simulations. (**C**) Analysis of the HHP and AD between two adjacent protomers within the same long-pitch strand in the bare actin filaments and in the unipolar-flexible, bipolar-flexible, and bipolar-fixed models using the conformations derived from the MD simulations.

**Figure S6.**
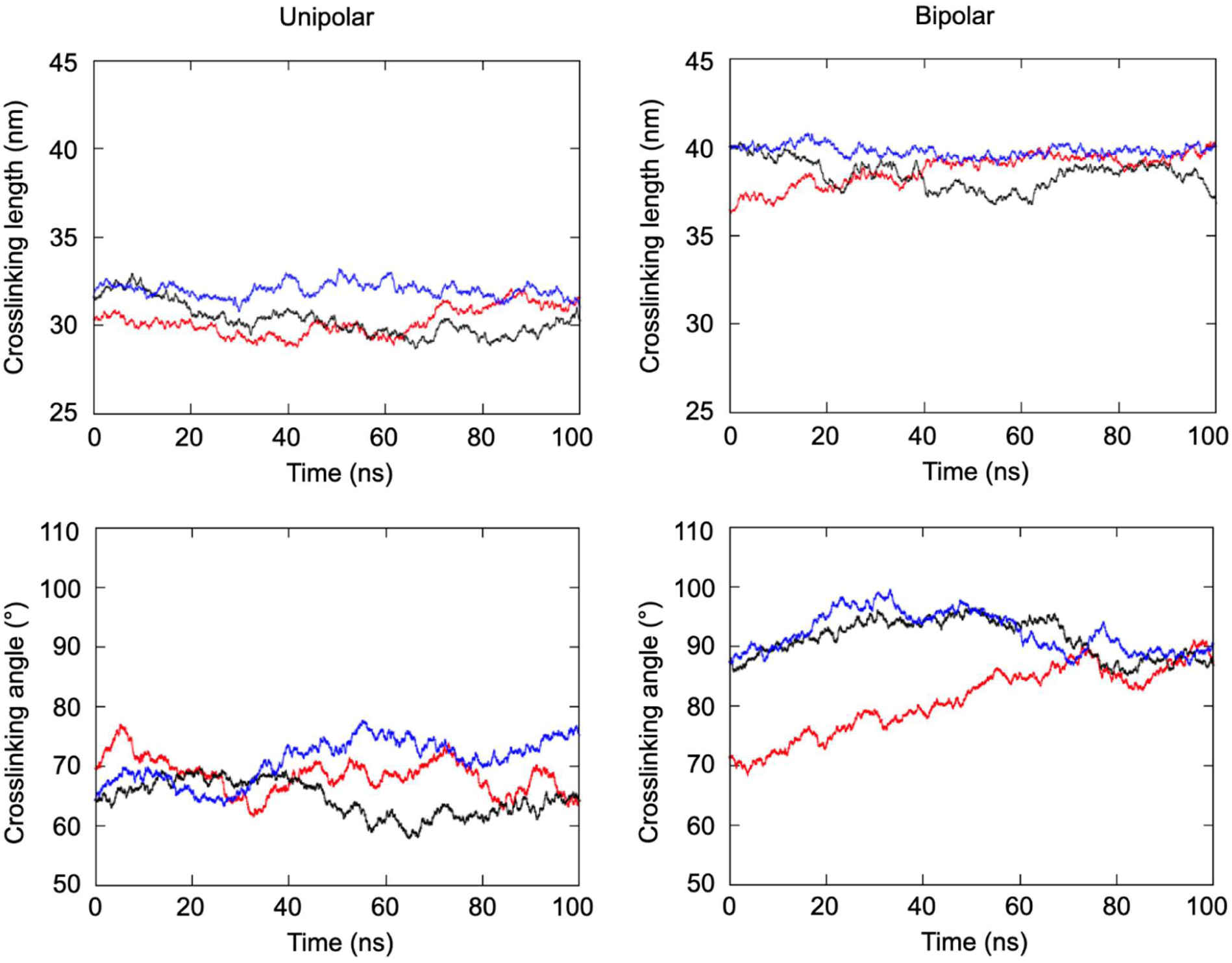
Time evolutions of the crosslinking angle and length of actin filaments. Different colors represent independent replicas.

**Figure S7.**
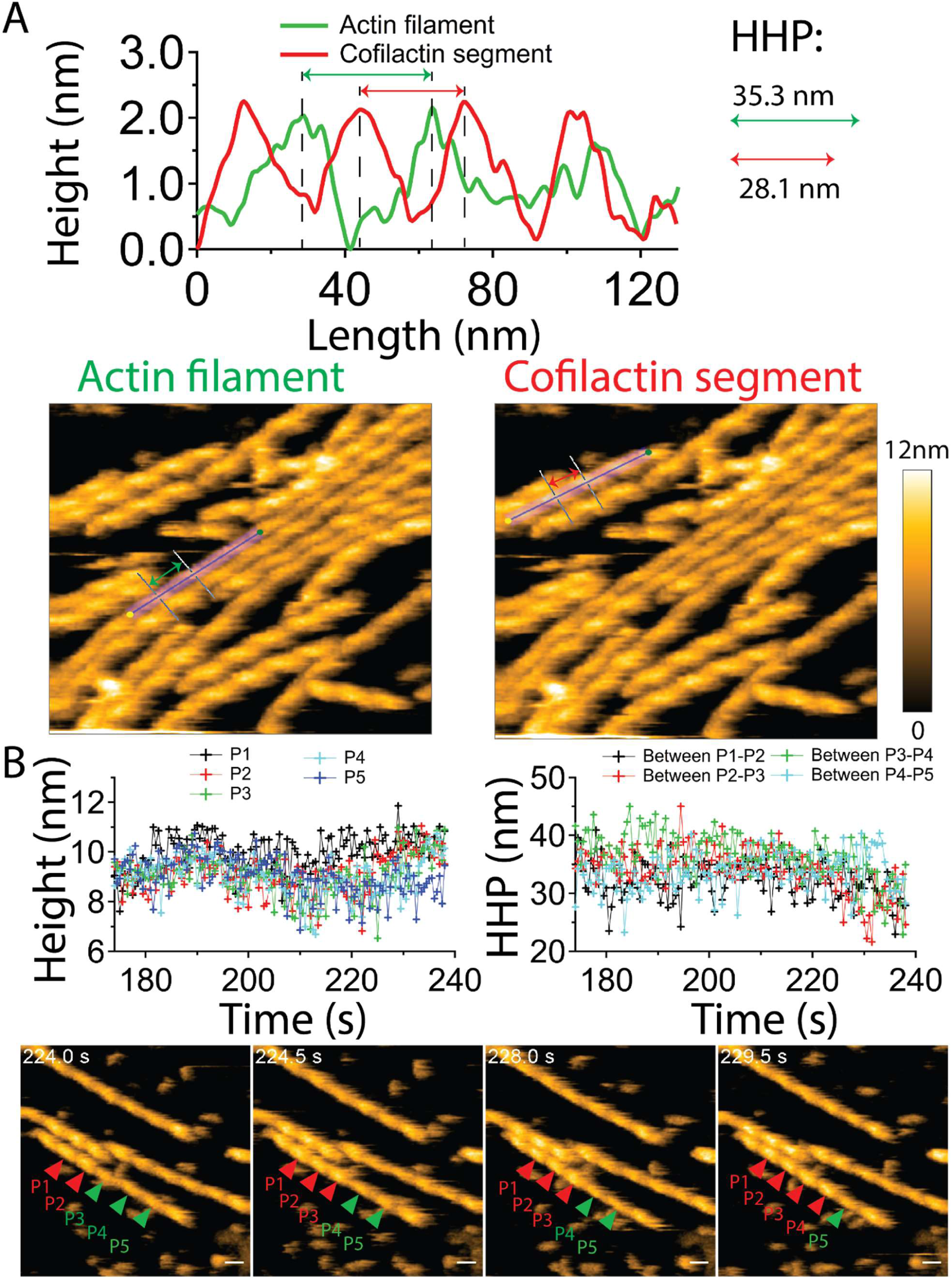
Cooperative binding and cluster formation of cofilin on actin filaments. **(A)** Longitudinal profiles to demonstrate the half-helical pitch (HHP) of normal actin filament and cofilactin segment in the same imaging field. Still AFM images were similarly reused in Figure 2B. (**B**) Still AFM images with time labels were shown to follow the cooperative binding and cluster formation of cofilin. Representative red and green arrowheads indicate the peaks (P1-P5) in cofilin clusters and normal actin segments, respectively. The time courses of peak height and HHP at positions P1-P5 were plotted to characterize their temporal fluctuations and monitor the progression of cooperative cofilin cluster formation via the peak heights and shortened HHP, following the approach used in our previous study (Ngo et al., 2024). Scale bars: 25 nm. Color scale: 0-12 nm.

**Figure S8.**
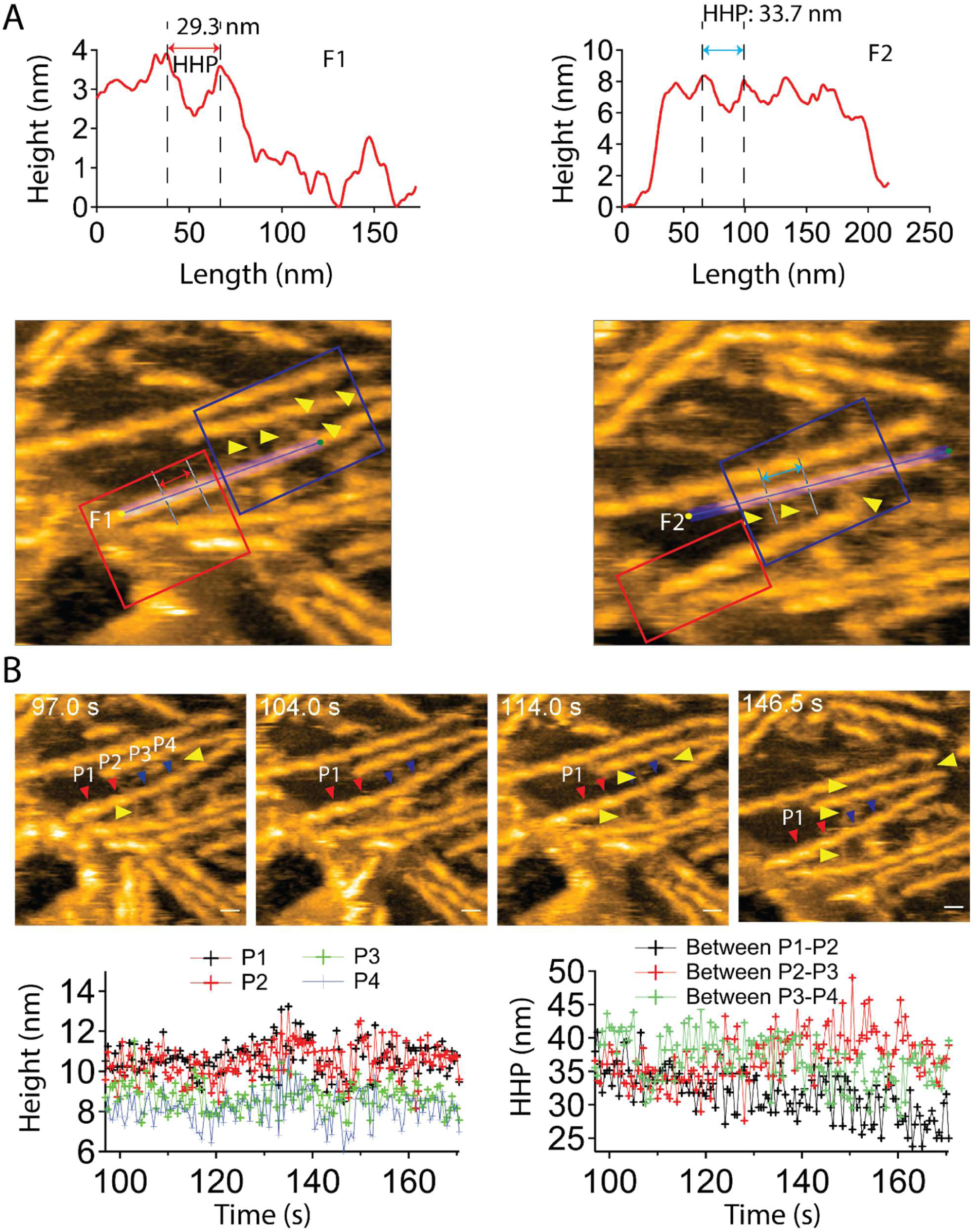
Binding and cluster formation of cofilin on alpha-actinin-crosslinked actin filaments. **(A)** Representative cofilactin (F1) and alpha-actinin-actin filament (F2) segments at low (red box) and high (blue box) alpha-actinin crosslinking densities, with corresponding longitudinal profiles showing shortened and normal HHPs. The height reference was defined along the F1 actin filament, but the reference to lipid membrane surface for F2. Yellow arrowheads representatively denote alpha-actinin molecules. **(B)** Still AFM images with time labels were shown to follow the binding and cluster formation of cofilin. Red and blue arrowheads indicate the peaks (P1-P5) in cofilactin (red) and cofilin-actinin-actin (blue) segments. The time courses of peak height and HHP at positions P1-P5 were plotted to characterize their temporal fluctuations and monitor the progression of cofilin cluster formation via the peak heights and shortened HHP. Yellow arrowheads representatively denote alpha-actinin molecules. Scale bars: 25 nm.

**Figure S9.**
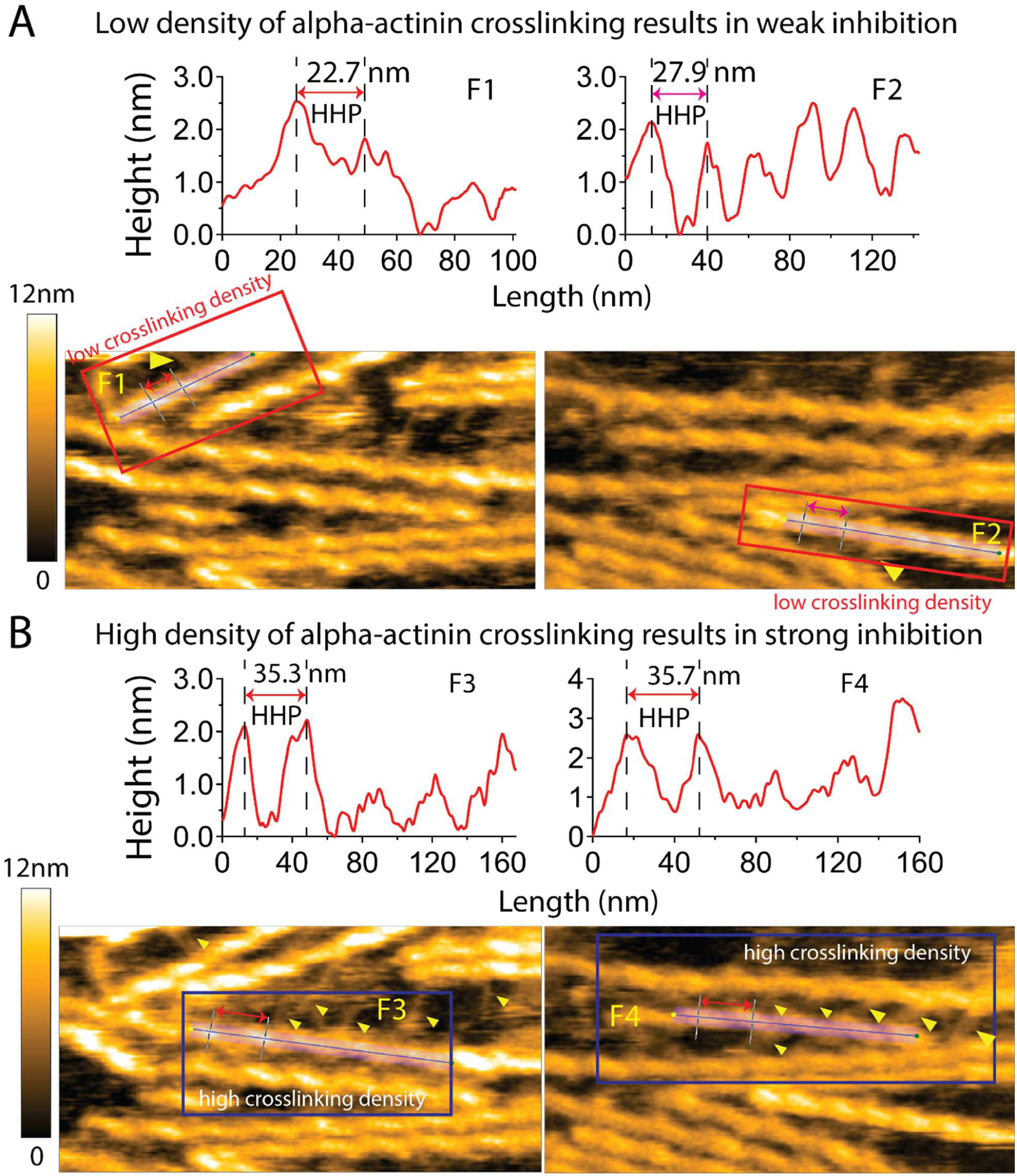
Effect of alpha-actinin crosslinking density on cofilin binding and cluster formation. (A) Representative cofilactin (F1, F2) segments at low alpha-actinin crosslinking densities, with corresponding longitudinal profiles showing shortened HHPs. Yellow arrowheads representatively denote alpha-actinin molecules. Still AFM images in A and B were similarly reused in Figure 2C. The height reference was defined along the actin filament. (B) Representative cofilin-actinin actin segments (F3, F4) at high alpha-actinin crosslinking densities, with corresponding longitudinal profiles showing normal HHPs. Yellow arrowheads representatively denote alpha-actinin molecules. The height reference was defined along the actin filament. Color scale: 0-12 nm.

**Figure S10.**
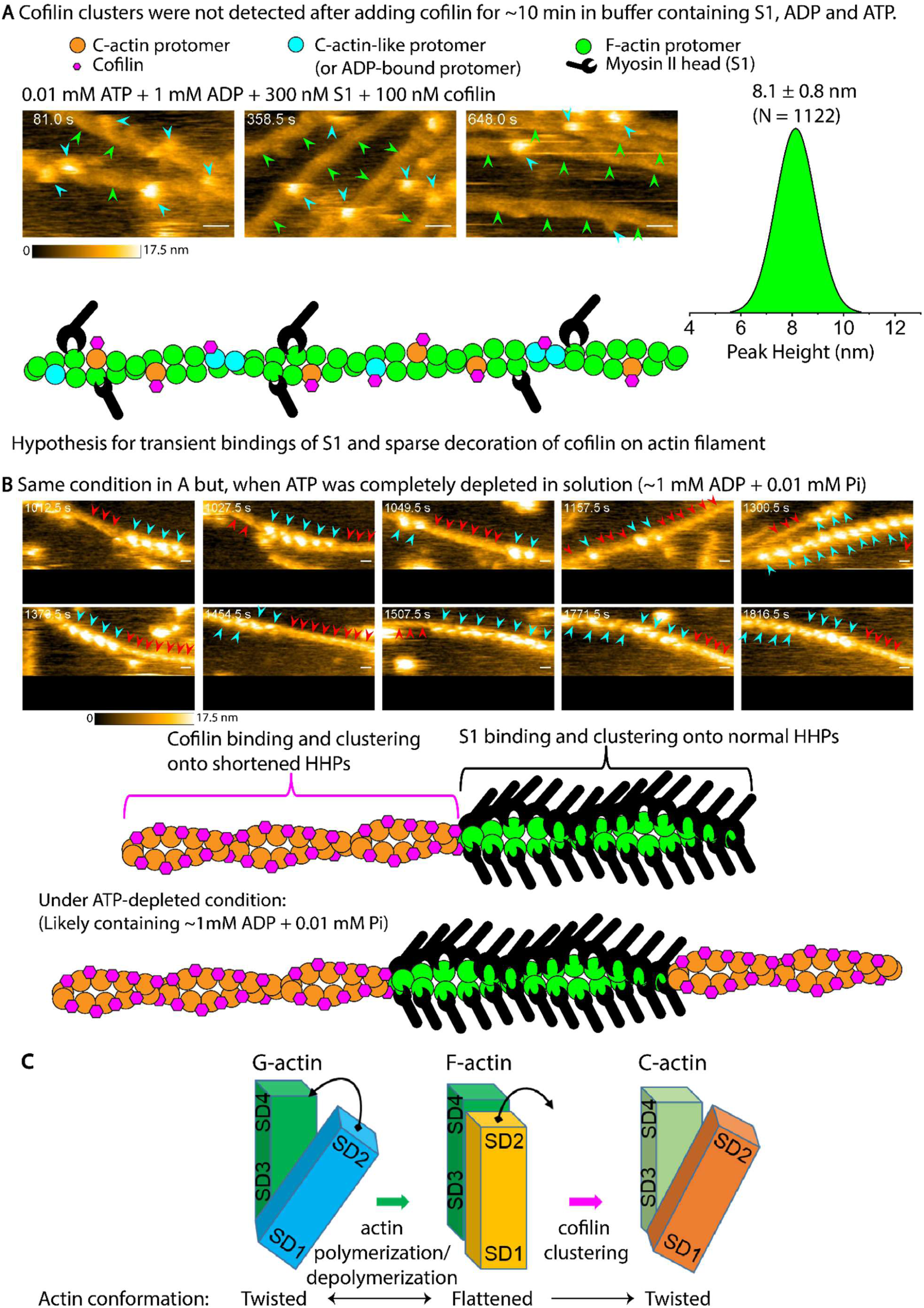
Segregation of cofilin and S1 clusters, as identified by their distinct short and normal HHP conformations and corresponding peak-height differences, visualized using HS-AFM. **(A)** Cofilin and S1 favorably interact with and utilize different HHP structures on the same actin filament to form their respective clusters. Green, cyan, and red arrowheads denote bare actin (lowest height, normal HHP), S1 molecules and S1-actin clusters (highest height, normal HHP), and cofilactin clusters (intermediate height between bare actin and S1-actin clusters, with a ∼25% reduction in HHP). Those are represented in these still AFM images, respectively. Actin filaments were initially imaged in F-buffer containing 0.01 mM ATP, 1 mM ADP, and 300 nM S1, followed by adding 100 nM cofilin. Owing to the high vertical sensitivity of HS-AFM (height detection limit of ∼0.1 nm), under the transient S1 binding condition in the presence of cofilin, we measured the peak height of every bare actin filament segment for ∼10 min following the cofilin addition. The histogram of peak heights for all bare actin segments is presented as mean ± SD (8.1 ± 0.8 nm, N = 1122 HHPs, analyzed from 10 actin segments). **(B)** As ATP levels gradually declined due to ATP hydrolysis (ATP → ADP + Pi) during the actin-myosin reaction, S1 binding to actin filaments persisted for longer durations and formed stable clusters when ATP was completely depleted. Time indicates the time after adding cofilin before (in A) and after (in B) ATP was completely depleted in solution. Scale bars: 25 nm. Color scales: 0-17.5 nm. Still AFM images were similarly reused in Figure 3A. **(C)** Schematic illustration of conformational transitions among G-actin monomers, F-actin protomers, and cofilin-bound C-actin protomers. The differences in protomer conformations were depicted with reference to previously reported atomic structures (Fujii et al., 2010; Fujii and Namba, 2017; Galkin et al., 2011).

**Figure S11.**
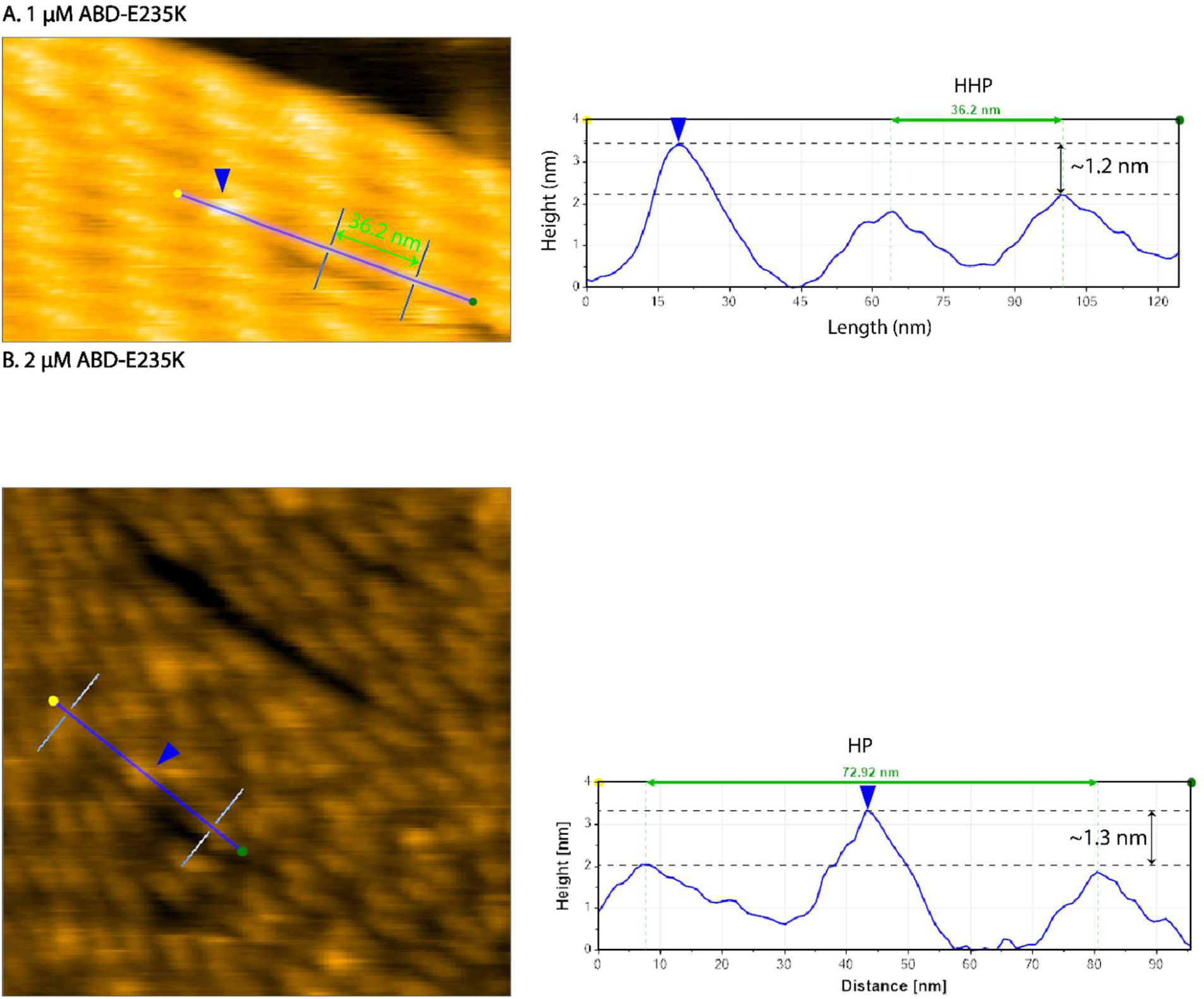
Transient binding of ABD-E235K to actin filaments does not affect length of half helices. (**A, B**) Representative HS-AFM images showing transient binding events of individual ABD-E235K molecules on actin filaments at concentrations of 1 and 2 μM, respectively. Length and height profiles confirm the binding of single ABD-E235K molecules to actin filaments and demonstrate that the filaments retain the normal long HHP during these transient interactions. The height reference was defined along the actin filament. Blue arrowheads indicate single ABD-E235K molecules bound onto actin filaments. Still AFM image in A was similarly reused in Figure 4A.

**Figure S12.**
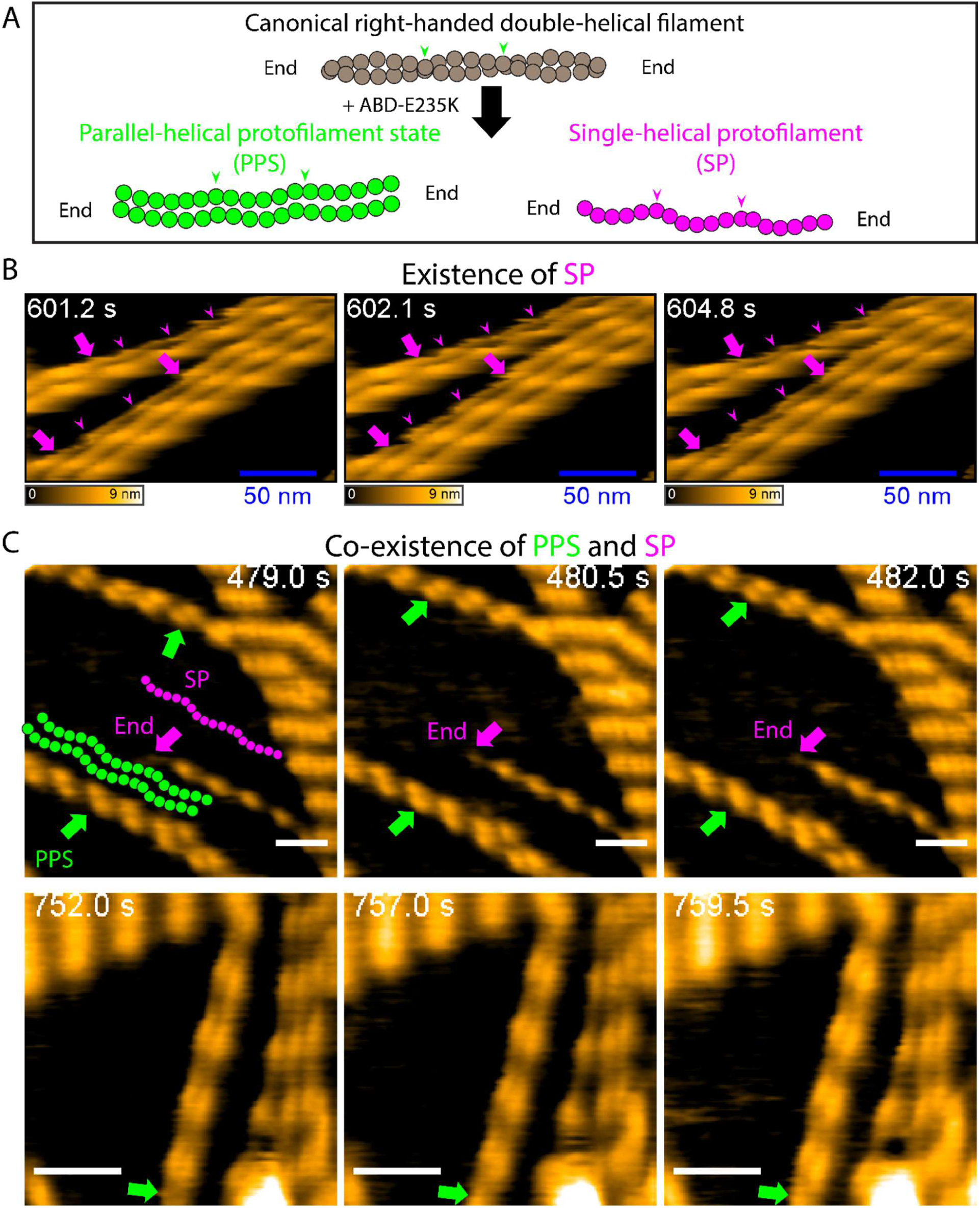
HS-AFM visualization of the single-helical protofilament (SP) structures and parallel-helical protofilament state (PPS) induced by ABD-E235K. **(A)** Schematic illustration of the ABD-E235K-induced conformational transition of canonical right-handed double-helical actin filaments into the SP and PPS with reduced heights but similar filament helical lengths. Purple and green arrowheads denote the peak heights of the half-helical repeats of the SP and PPS, respectively. (**B, C**) HS-AFM Video showing the emergence of the SP and PPS in actin filaments following the addition of 2.0 μM ABD-E235K. Time stamps indicate the elapsed time after ABD-E235K addition. To obtain high-resolution images of the resulting structures, the imaging buffer was exchanged to remove transiently bound ABD-E235K molecules. Actin filaments were pre-incubated with ABD-E235K for approximately 30 min at room temperature before immobilization on lipid membranes for HS-AFM imaging. Purple arrows indicate the ends of the SP. Green arrows indicate the PPS. Purple arrowheads denote the peak heights of the half-helical repeats of the SP. Scale bar, 50 nm. Still AFM images in B and C were similarly reused in Figure 4C,D or Figure S13A.

**Figure S13.**
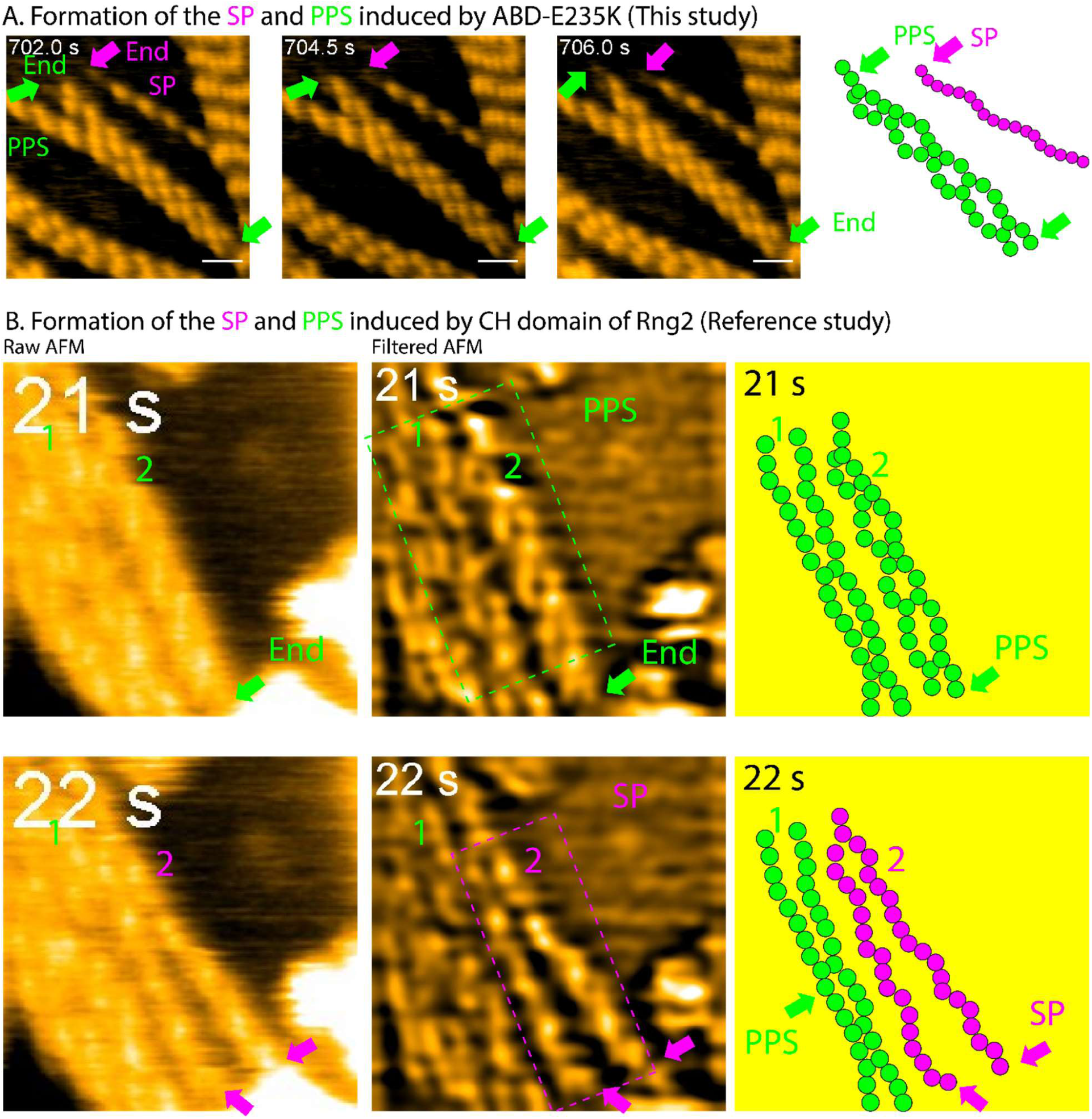
Alpha-actinin ABD-E235K and Rng2-CHD induce a transition of canonical right-handed double-helical actin filaments into the single-helical protofilament (SP) structure and parallel-helical protofilament state (PPS). **(A)** This study: HS-AFM images revealing the SP and PPS of actin filaments induced by incubation with 2 μM ABD-E235K. Actin filaments were pre-incubated with ABD-E235K for approximately 30 min at room temperature before immobilization on the lipid membranes for HS-AFM imaging. Purple and green arrows denote the ends of the SP and PPS, respectively. Still AFM images were similarly reused in Figure S12C. **(B)** Reference study: Still AFM images were similarly reused in Figure S5 in our previous study (Hayakawa et al., 2022), we reported that the calponin homology domain (CHD) of Rng2 induces separation of canonical double-helical actin filaments. In those experiments, actin filaments were gently immobilized on positively charged lipid membranes composed of DPPC/DPTAP (90/10 wt%). HS-AFM imaging was performed in buffer containing 25 mM KCl, 2 mM MgCl₂, 50 μM ATP, 1 mM ADP, 10 mM DTT, 20 mM HEPES (pH 7.4), and 5.7 μM CHD. Upon careful reanalysis of the HS-AFM Videos, we found that the Rng2 CHD not only separated the two protofilaments but also induced the similar transformation of the canonical double-helical filament into the SP and PPS. These structural transitions are illustrated by the AFM images and cartoons between the 21 s and 22 s frames (green and purple dashed boxes). During this process, one SP of filament 2 detached from the lipid membrane while the other remained membrane-bound. Notably, the helical periodicity remained clearly visible in the PPS and each individual SP after complete separation. This structural feature serves as an important reference for the SP and PPS for the present study, while further supporting the notion that it represents a previously underappreciated property of actin filament. Purple and green arrows denote the ends of the SP and PPS, respectively.

**Figure S14.**
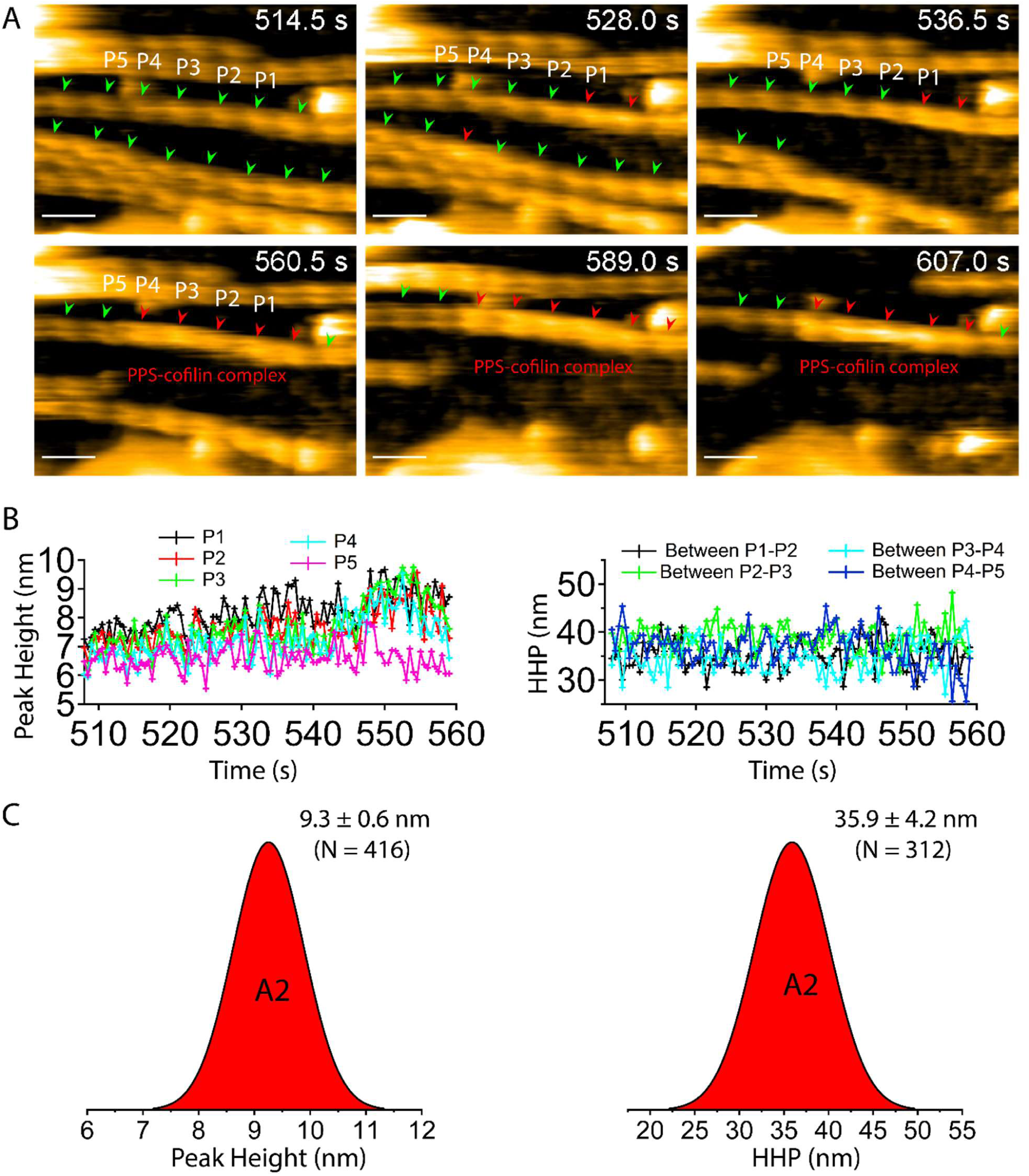
HS-AFM visualization of cofilin binding and cluster formation on the PPS-like-helical pitch segments. **(A)** Actin filaments were incubated with 2.0 μM ABD-E235K in solution for approximately 30 min before being immobilized on a lipid membrane. The imaging buffer was then exchanged to remove transiently bound ABD-E235K molecules prior to the addition of 3.5 μM cofilin. Time stamps indicate the elapsed time after cofilin addition. Green arrowheads denote peak heights of PPS-like-helical pitch segments, whereas red arrowheads denote peak heights of PPS-cofilin complex segments, used to measure HHPs accordingly. Scale bar, 50 nm. Still AFM images were similarly reused in Figure S15. **(B)** Kinetic analysis of peak height and HHP fluctuations between P1 and P5 to follow cofilin cluster growth and HHP changes. As seen, the mean HHPs were almost unchanged during cofilin cluster growth. (**B**) Histograms of the peak height and HHP of PPS-cofilin complex segments. Data are presented as mean ± SD. N = 416 peak heights and 312 HHPs analyzed from 5 PPS-cofilin complex segments. Histograms for PPS-cofilin segments (A2, in Figure S15) were made from peak heights and HHPs.

**Figure S15.**
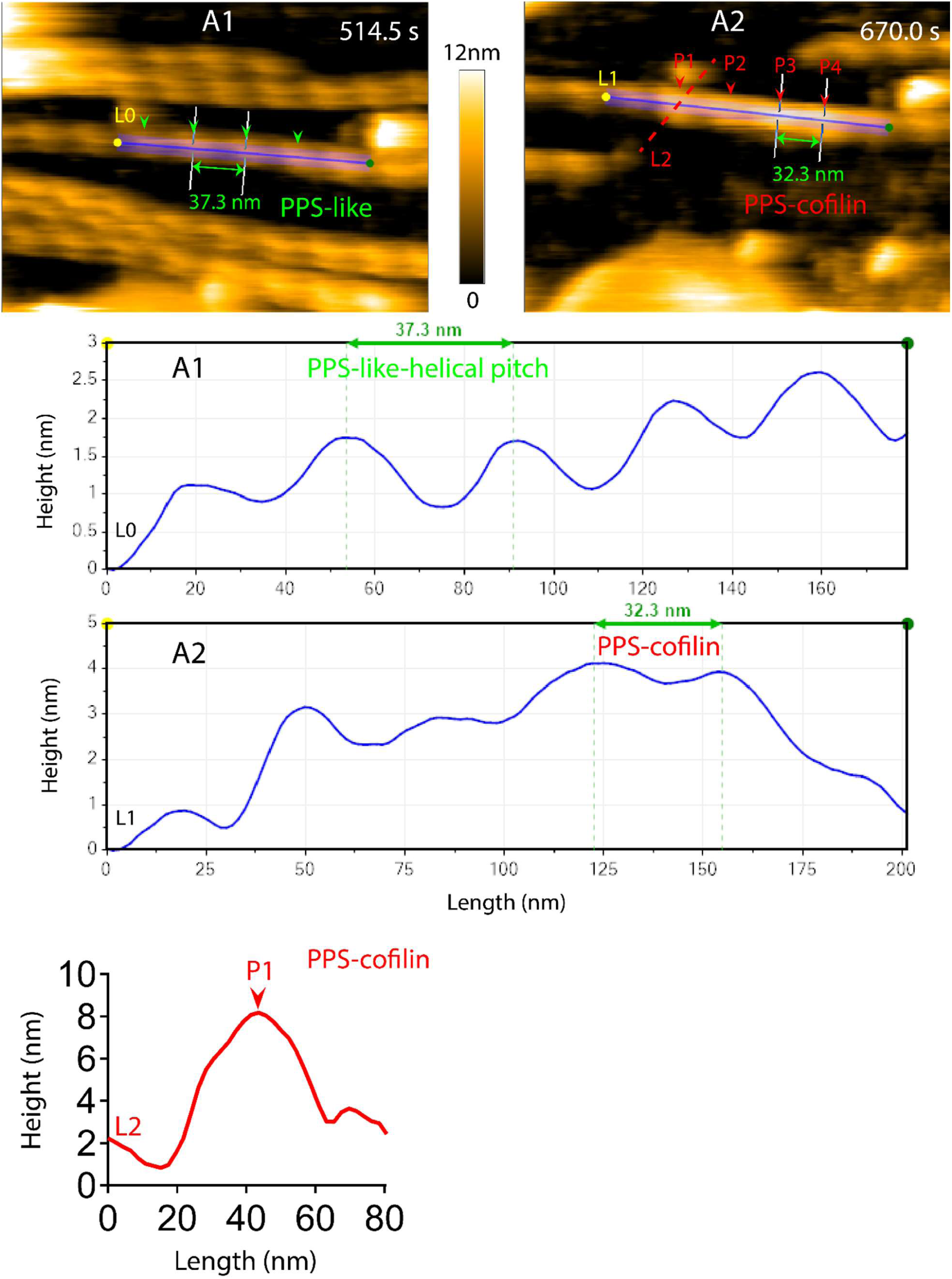
HS-AFM visualization of cofilin binding and cluster formation on the PPS-like-helical pitch segments. Actin filaments were incubated with 2.0 μM ABD-E235K in solution for approximately 30 min, gently immobilized on a lipid membrane, and imaged by HS-AFM after buffer exchange to remove unbound ABD-E235K. Subsequently, 3.5 μM cofilin was introduced into the imaging chamber. Representative HS-AFM images show the coexistence of PPS-like (A1) and PPS-cofilin (A2) segments within the same actin filament. Time stamps indicate the elapsed time after cofilin addition. Green arrowheads denote peaks of PPS-like-helical pitch segments, whereas red arrowheads denote peaks of PPS-cofilin complex segments. Representative length-height profiles of the regions indicated by L0, L1, and L2 are shown. For the L0 and L1 profiles, the height reference was defined along the actin filament, resulting in lower measured heights than for L2, where the substrate surface was used as the reference. Color scale: 0 – 12 nm. Still AFM images were similarly reused in Figure S14.

**Figure S16.**
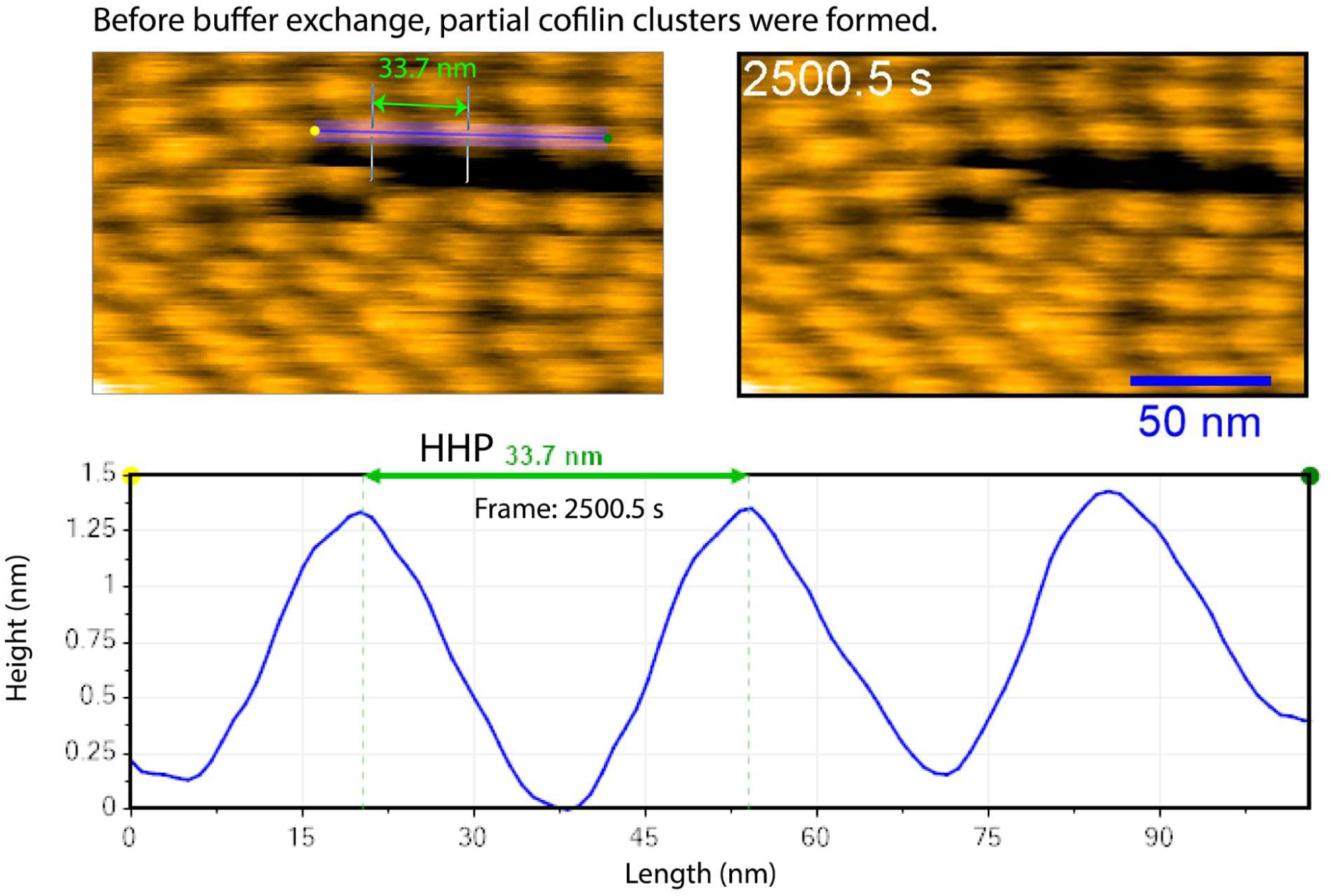
Transient binding of ABD-E235K perturbs complete cofilin-induced helical shortening and decoration of actin filaments. Actin filaments were first immobilized on a lipid membrane, after which 2.0 μM ABD-E235K was added and imaged by HS-AFM for approximately 30 min, followed by the addition of 3.5 μM cofilin. The height reference was defined along the actin filament. Time labels indicate the elapsed time after cofilin addition. Scale bars, 50 nm.

**Figure S17.**
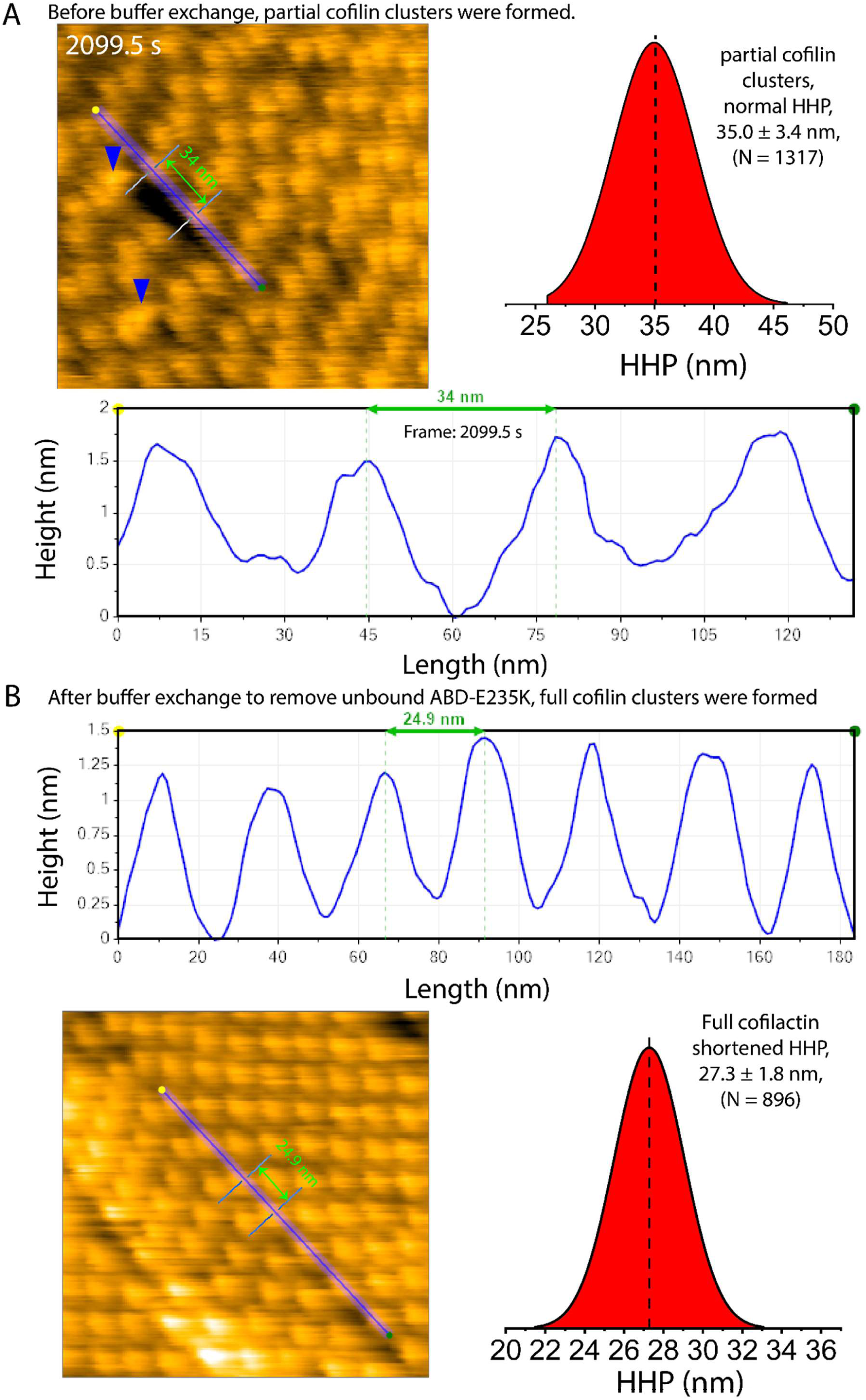
Actin filaments are fully decorated by cofilin clusters following removal of unbound ABD-E235K. **(A)** Before buffer exchange: Actin filaments were first immobilized on a lipid membrane, after which 2.0 μM ABD-E235K was added and imaged by HS-AFM for approximately 30 min, followed by the addition of 3.5 μM cofilin. The height reference was defined along the actin filament. Blue arrowheads denote individual ABD-E235K binding on actin filaments. Time labels indicate the elapsed time after cofilin addition. Related **Video S10A**. **(B)** After buffer exchange: Continuation of the experimental conditions as shown in **A and Video S10A**, buffer exchange was performed for approximately 5 min to remove unbound ABD-E235K before continued HS-AFM imaging. The height reference was defined along the actin filament. Related **Video S10B**.

**Figure S18.**
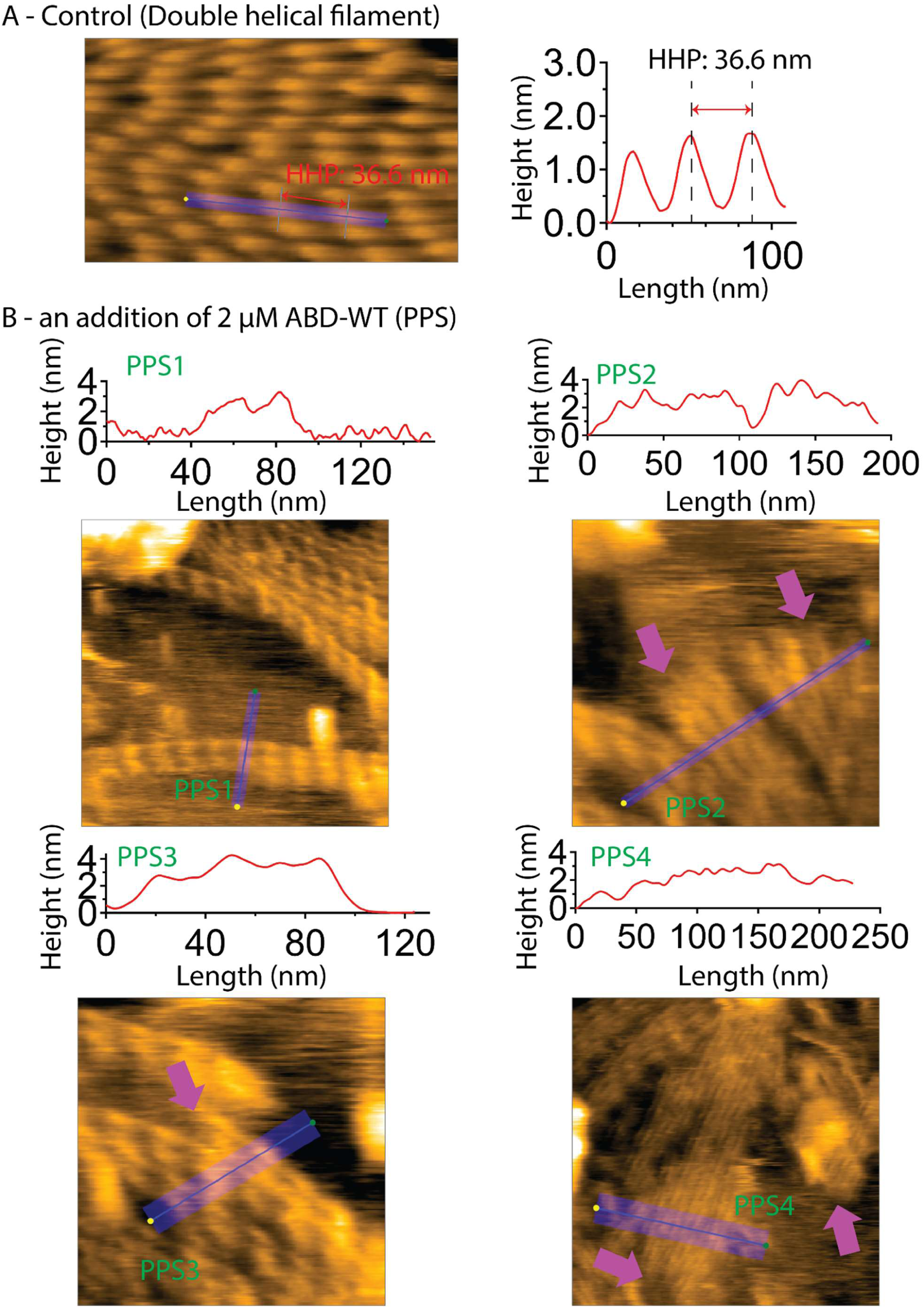
Binding of ABD-WT induces a transition of the canonical right-handed double-helical actin filament to the parallel-helical protofilament state (PPS). **(A)** Representative HS-AFM image of canonical right-handed double-helical actin filament and its longitudinal profiles. The height reference was defined along the actin filament. **(B)** Representative HS-AFM images of the PPS formed in actin filaments following incubation with 2 μM ABD-WT. Actin filaments were gently immobilized on lipid membranes containing 10 % DPTAP, subsequently added into the imaging chamber with 2 μM ABD-WT and HS-AFM imaging. Purple arrows indicate the ends of the PPS. Related to conditions shown in Figure 5 and Video S11. The height reference was defined as the lipid membrane surface. A still AFM image in B was similarly reused in Figure 5B.

**Figure S19.**
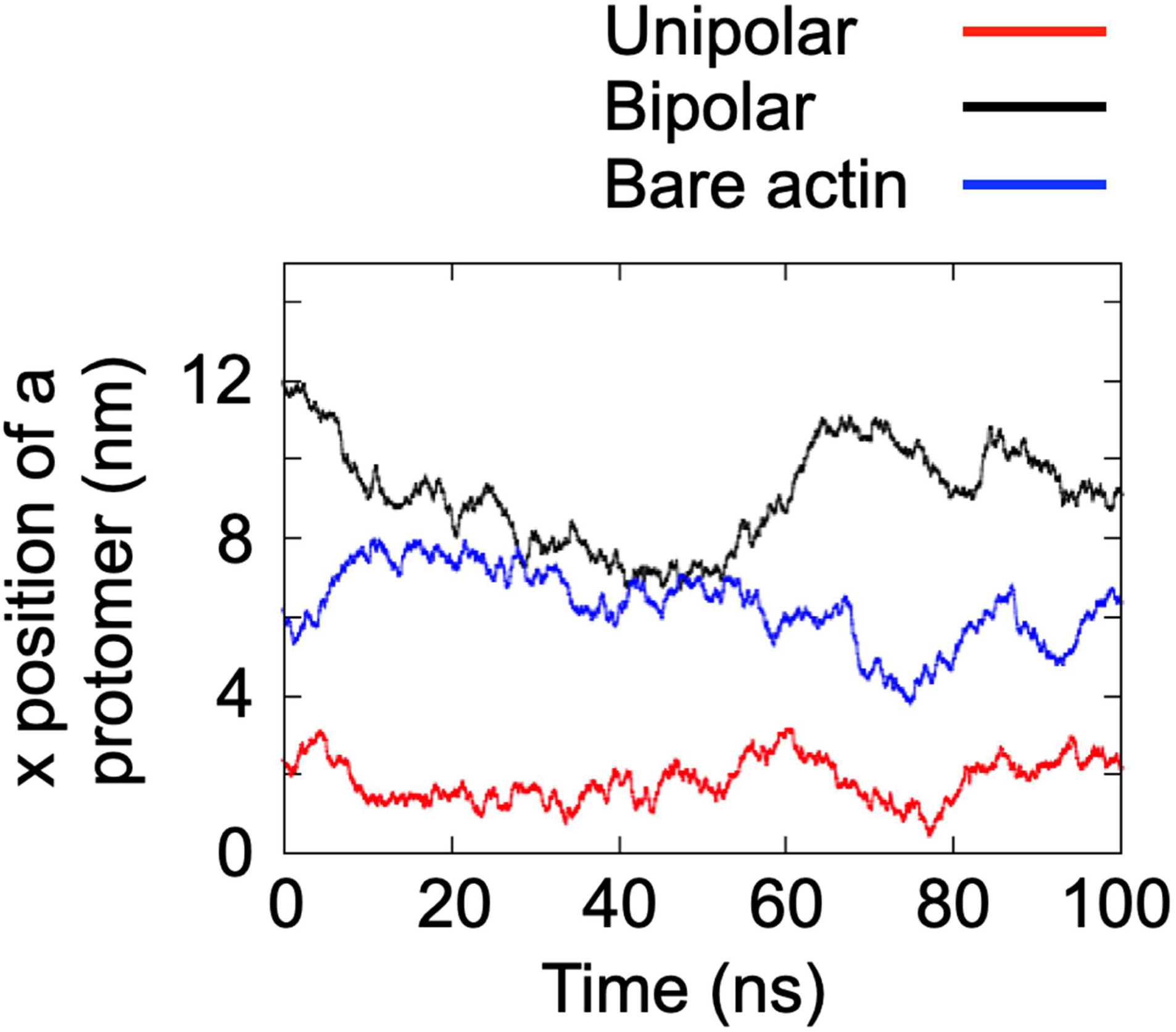
Time evolutions of the x-position of the representative protomer in the actin filament.

**Figure S20.**
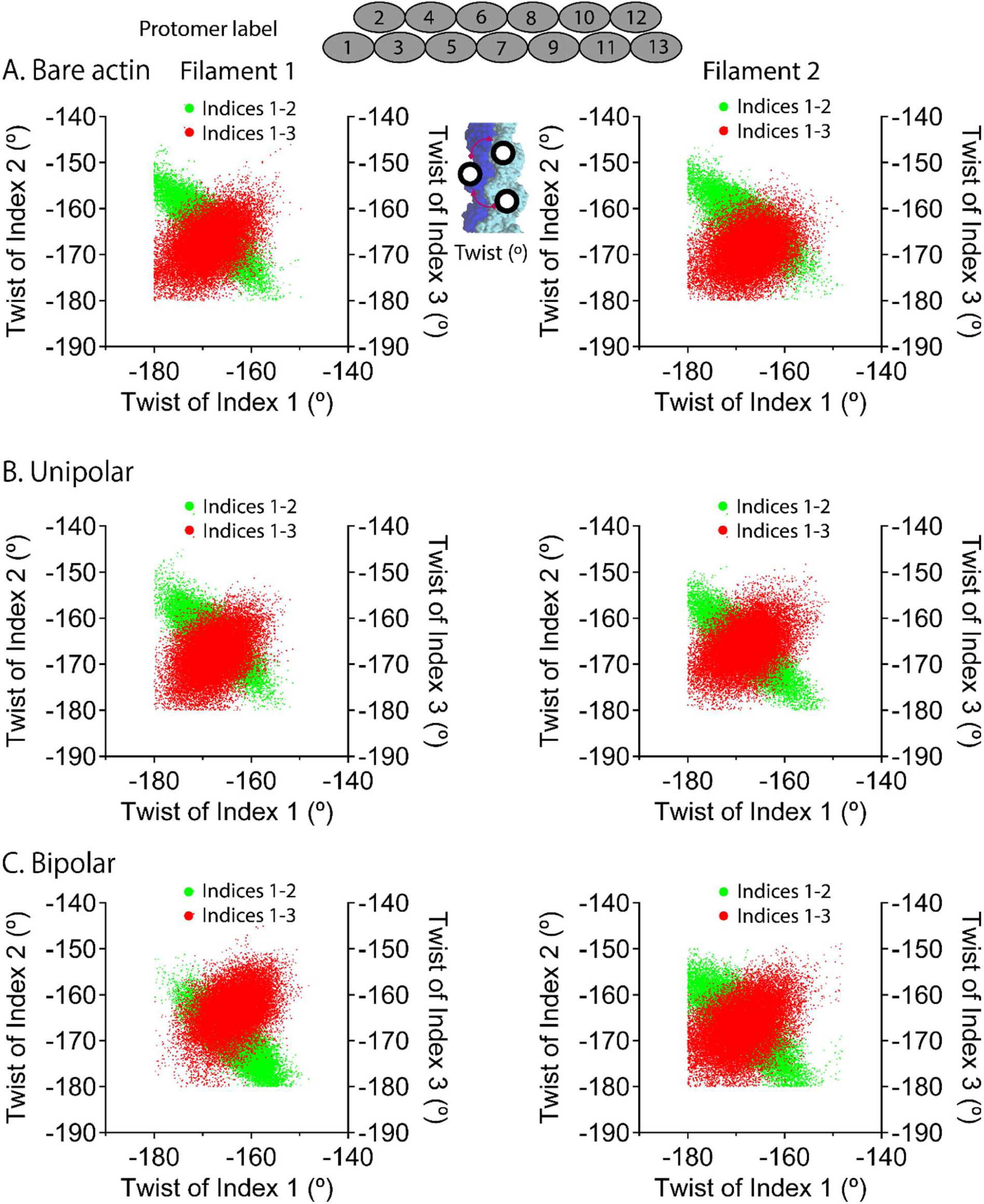
Correlations of actin subunit twist dynamics. (**A–C**) Representative correlation analyses of the twist motions of actin subunits 1, 2, 3 and 4 derived from MD simulations of bare actin filaments and alpha-actinin-crosslinked actin filaments in unipolar and bipolar arrangements. Negative correlations of twist dynamics between adjacent subunits on opposite strands are shown in green (indices 1-2), whereas positive correlations between neighboring subunits within the same strand are shown in red (indices 1-3). Correlation matrices were generated from 25,000 simulation snapshots acquired over 300 ns. Filaments 1 and 2 correspond to F1 and F2, respectively, and actin subunits are numbered as in Figure 6. Twist of index 1: Twist between protomer n and n + 1; Twist of index 2: Twist between protomer n + 1 and n + 2; Twist of index 3: Twist between protomer n + 2 and n + 3. Where n = 1.

**Figure S21.**
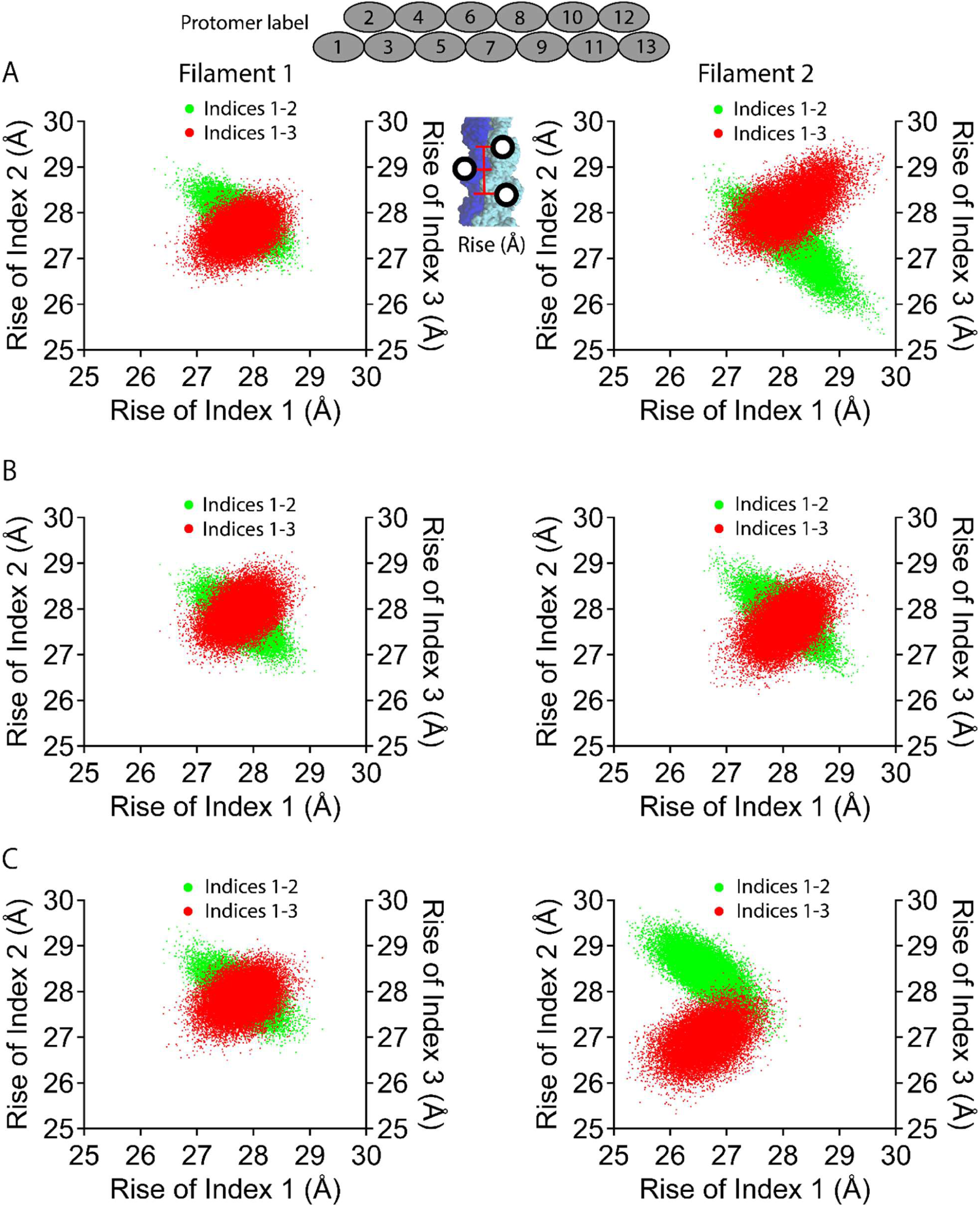
Correlations of actin subunit rise dynamics. **(A–C)** Representative correlation analyses of the rise dynamics of actin subunits 1, 2, 3 and 4 derived from MD simulations of bare actin filaments and alpha-actinin-crosslinked actin filaments in unipolar and bipolar arrangements. Negative correlations of rise dynamics between adjacent subunits on opposite strands are shown in green (indices 1-2), whereas positive correlations between neighboring subunits within the same strand are shown in red (indices 1-3). Correlation matrices were generated from 25,000 simulation snapshots acquired over 300 ns. Filaments 1 and 2 correspond to F1 and F2, respectively, and actin subunits are numbered as in Figure 6. Rise of index 1: Rise between protomer n and n + 1; Rise of index 2: Rise between protomer n + 1 and n + 2; Rise of index 3: Rise between protomer n + 2 and n + 3. Where n = 1.

**Figure S22.**
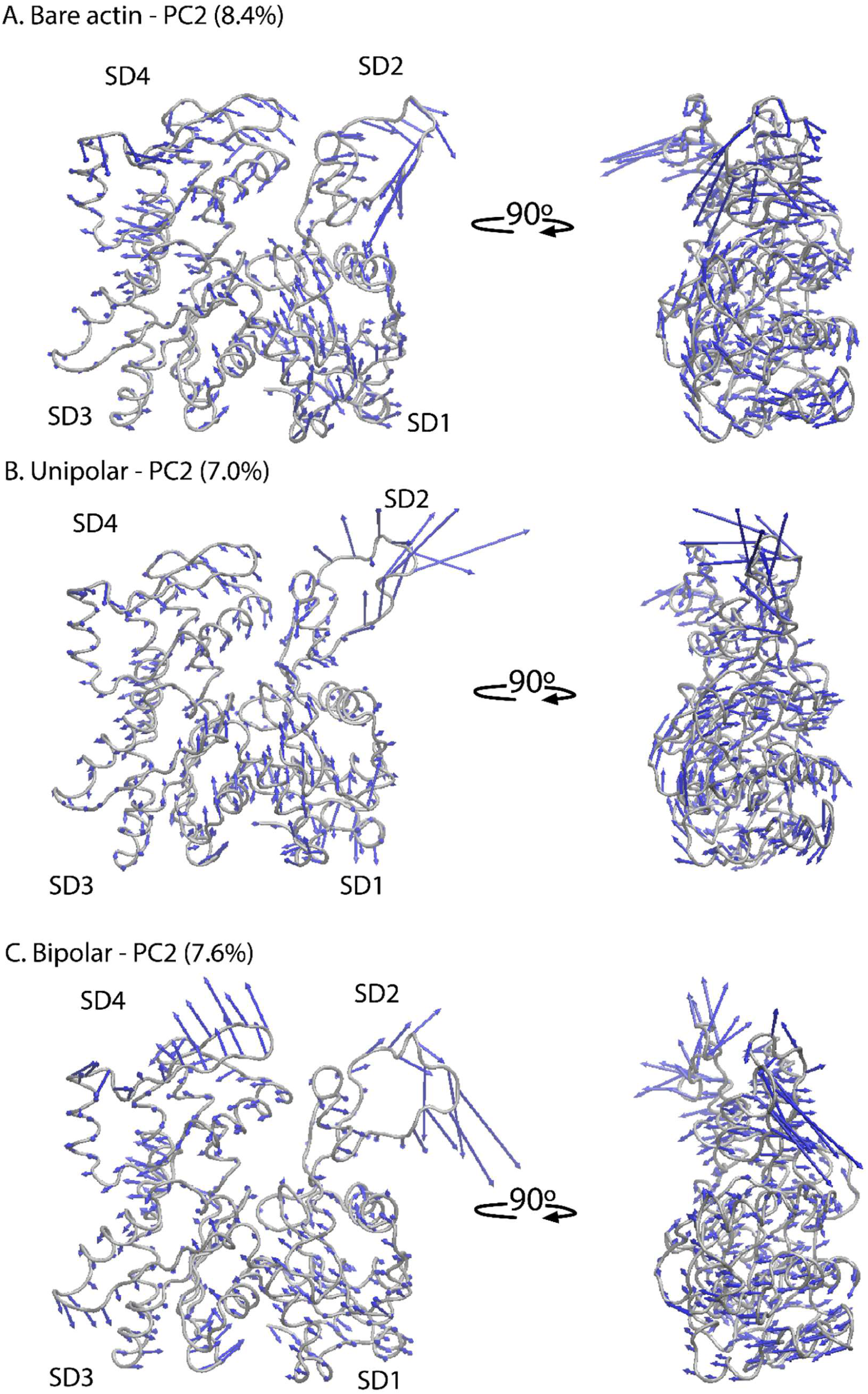
Analysis of atomic conformational dynamics of actin protomers from the bare actin, unipolar, and bipolar filament models using MD simulations and PCA. (**A, B, C**) Principal component 2 (PC2) captured vibration of the flattening motions of actin protomers in bare actin, unipolar, and bipolar filament models. Actin structures are shown in tube representation with the four subdomains (SD1-SD4) highlighted. The eigenvalues of PC2 indicate the percentage contribution of the dominant motions. Eigenvector components depicting the direction of motion are indicated by blue arrows.

**Figure S23.**
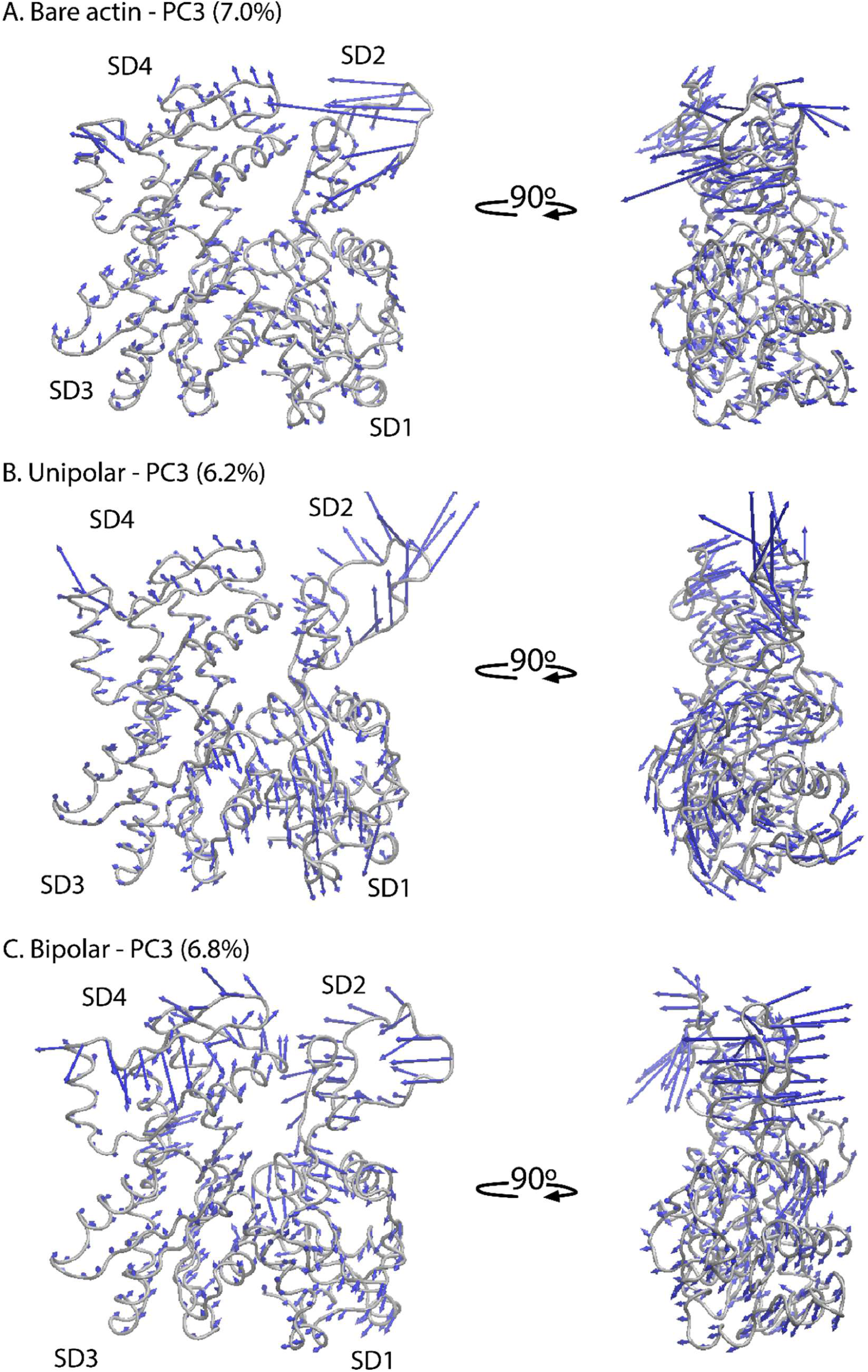
Analysis of atomic conformational dynamics of actin protomers from the bare actin, unipolar, and bipolar filament models using MD simulations and PCA. (**A, B, C**) Principal component 3 (PC3) captured vibration of the flattening motions of actin protomers in bare actin, unipolar, and bipolar filament models. Actin structures are shown in tube representation with the four subdomains (SD1-SD4) highlighted. The eigenvalues of PC3 indicate the percentage contribution of the dominant motions. Eigenvector components depicting the direction of motion are indicated by blue arrows.

**Figure S24.**
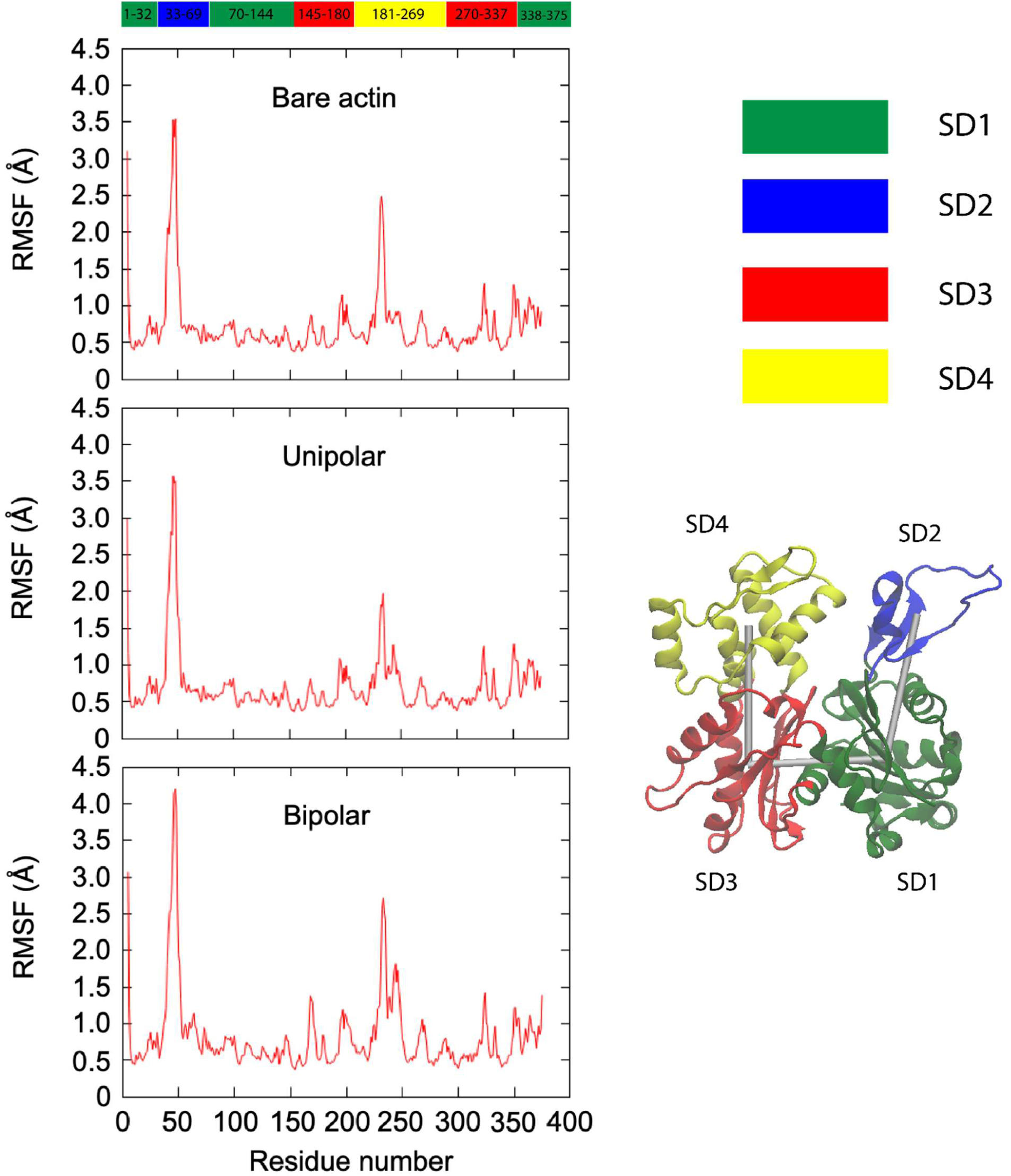
Root mean square fluctuation (RMSF) analysis of actin protomers. Residue-wise RMSF profiles of actin protomers calculated from molecular dynamics (MD) simulations of bare actin filaments and alpha-actinin-crosslinked actin filaments in the unipolar and bipolar configurations.

**Figure S25.**
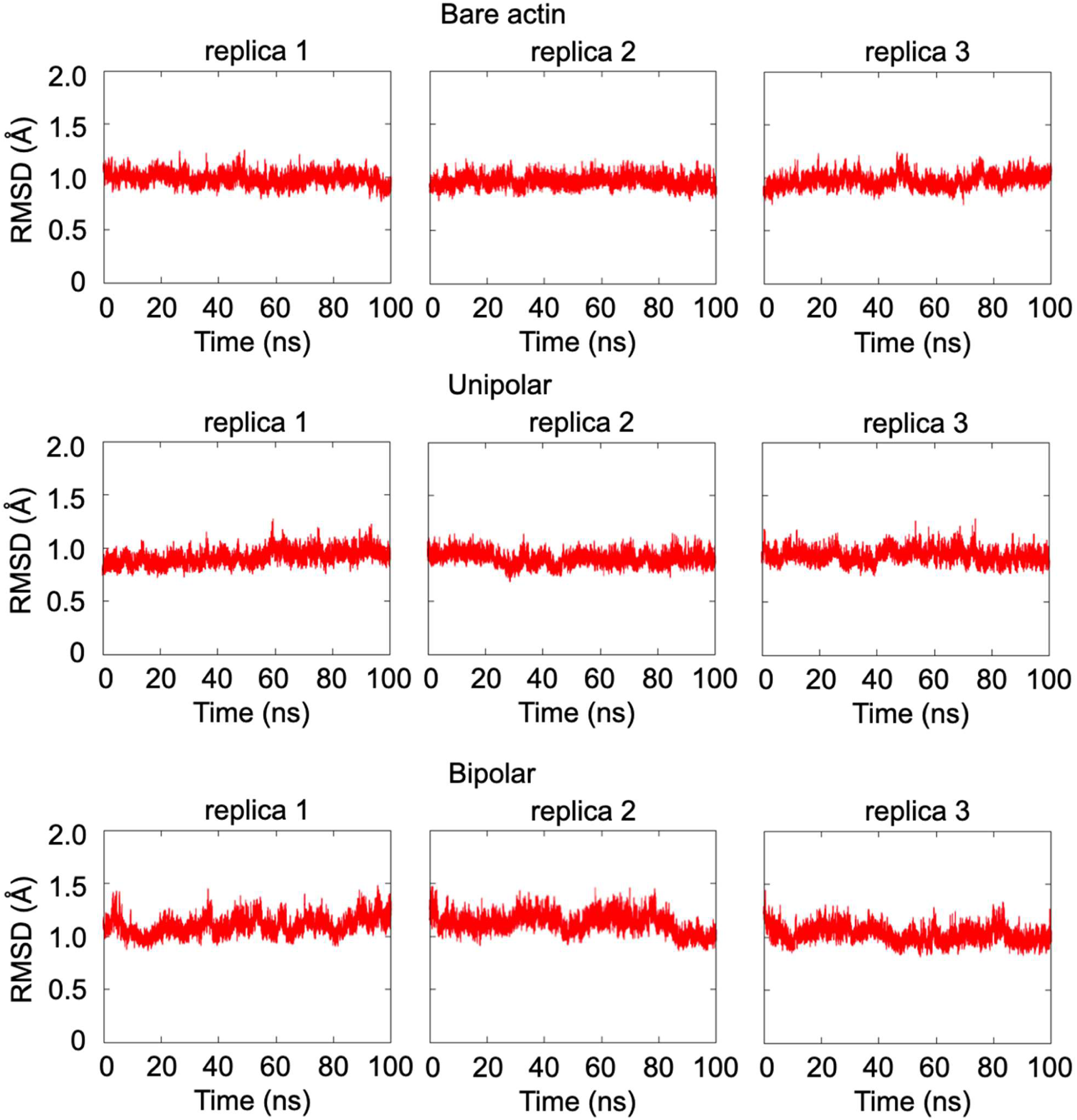
RMSD of the alpha-carbons in protomers from the 8D15 pdb structure.

**Figure S26.**
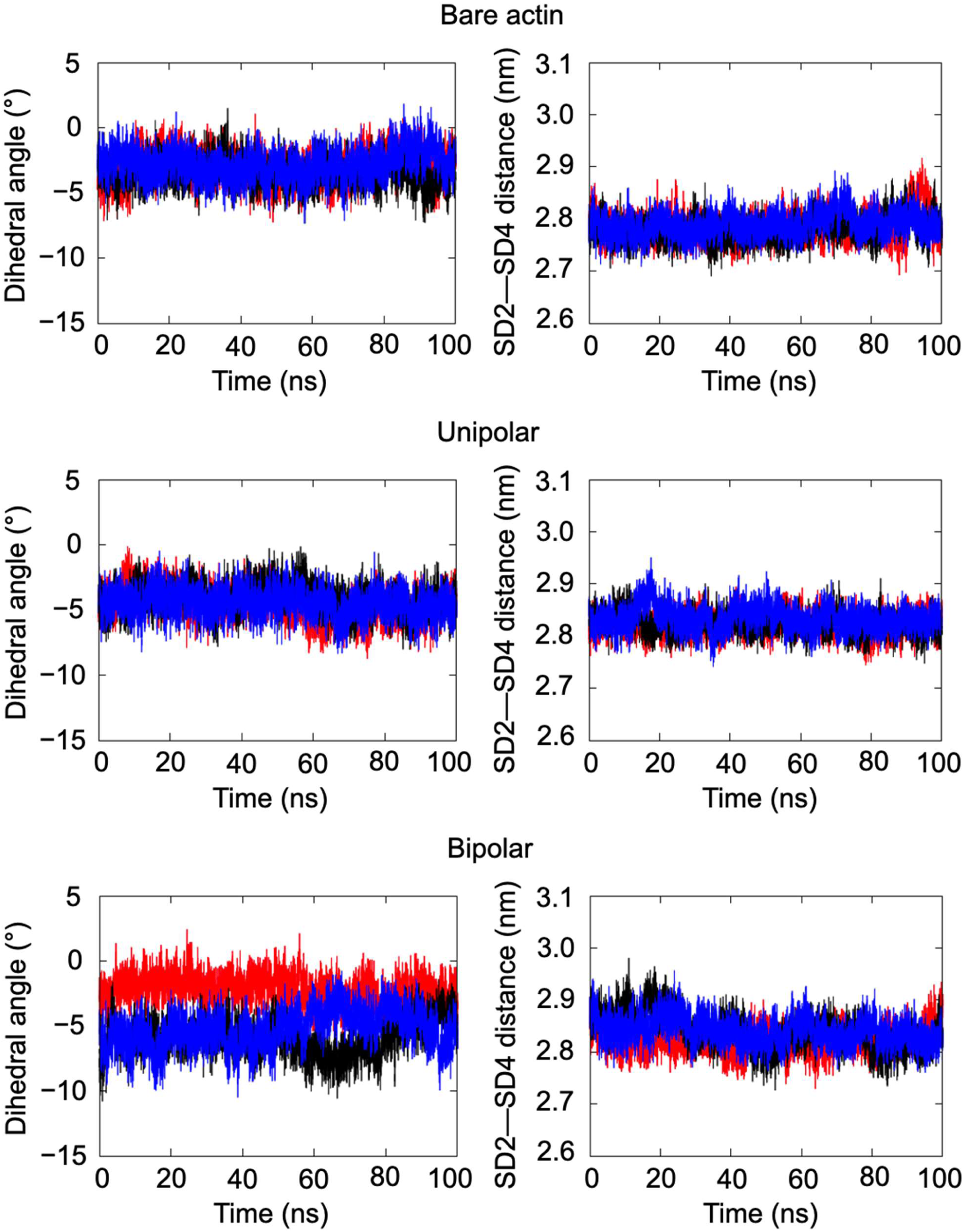
Time evolutions of the dihedral angle and SD2-SD4 distance in protomers. Different colors represent independent replicas.

**Figure S27.**
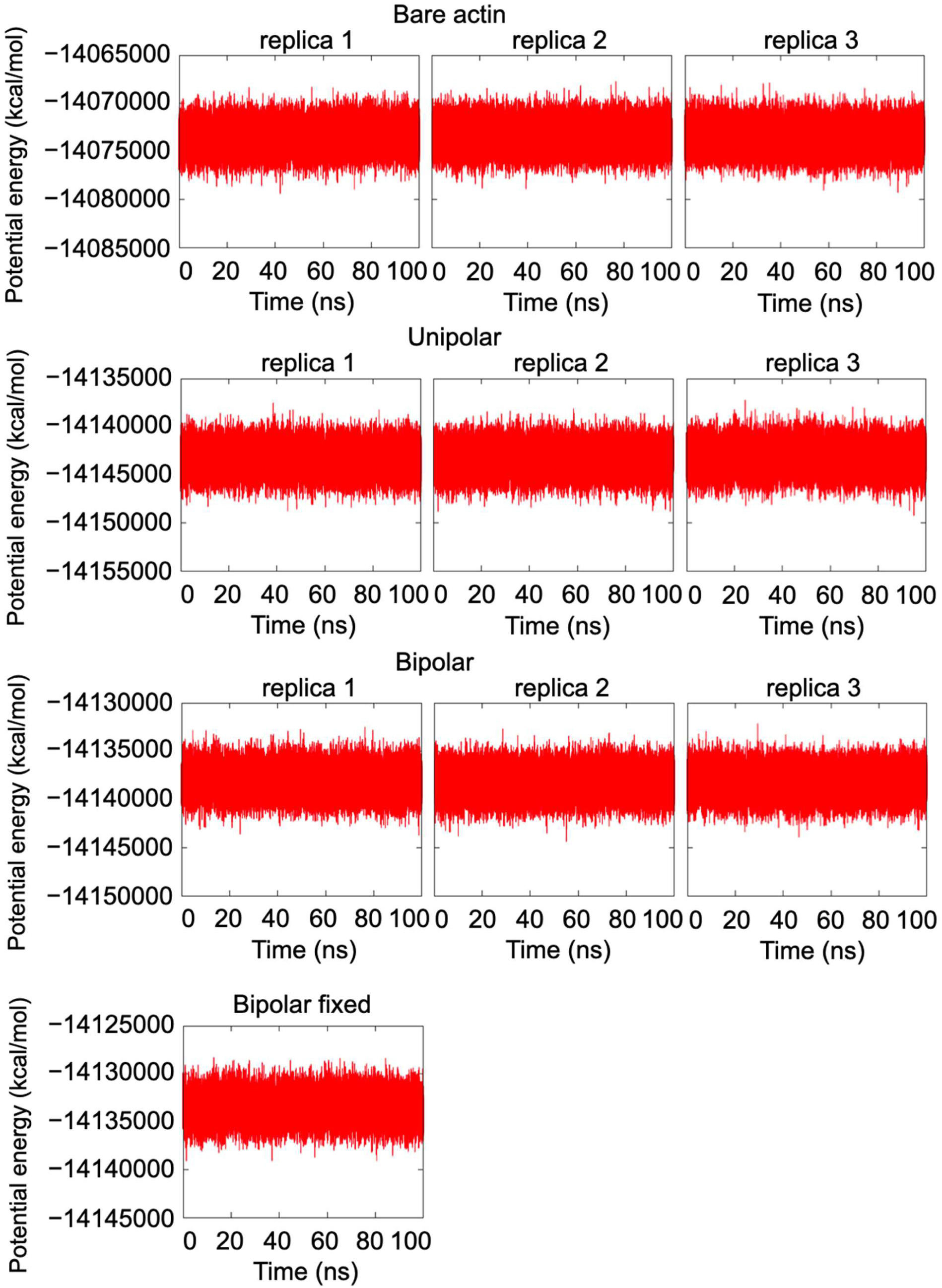
Time evolutions of the potential energy of the systems. As expected, the total energy varied between systems, but there were no differences among the replicas. During the simulations, the potential energy remained constant, indicating that the systems under simulation were stable.

**Figure S28.**
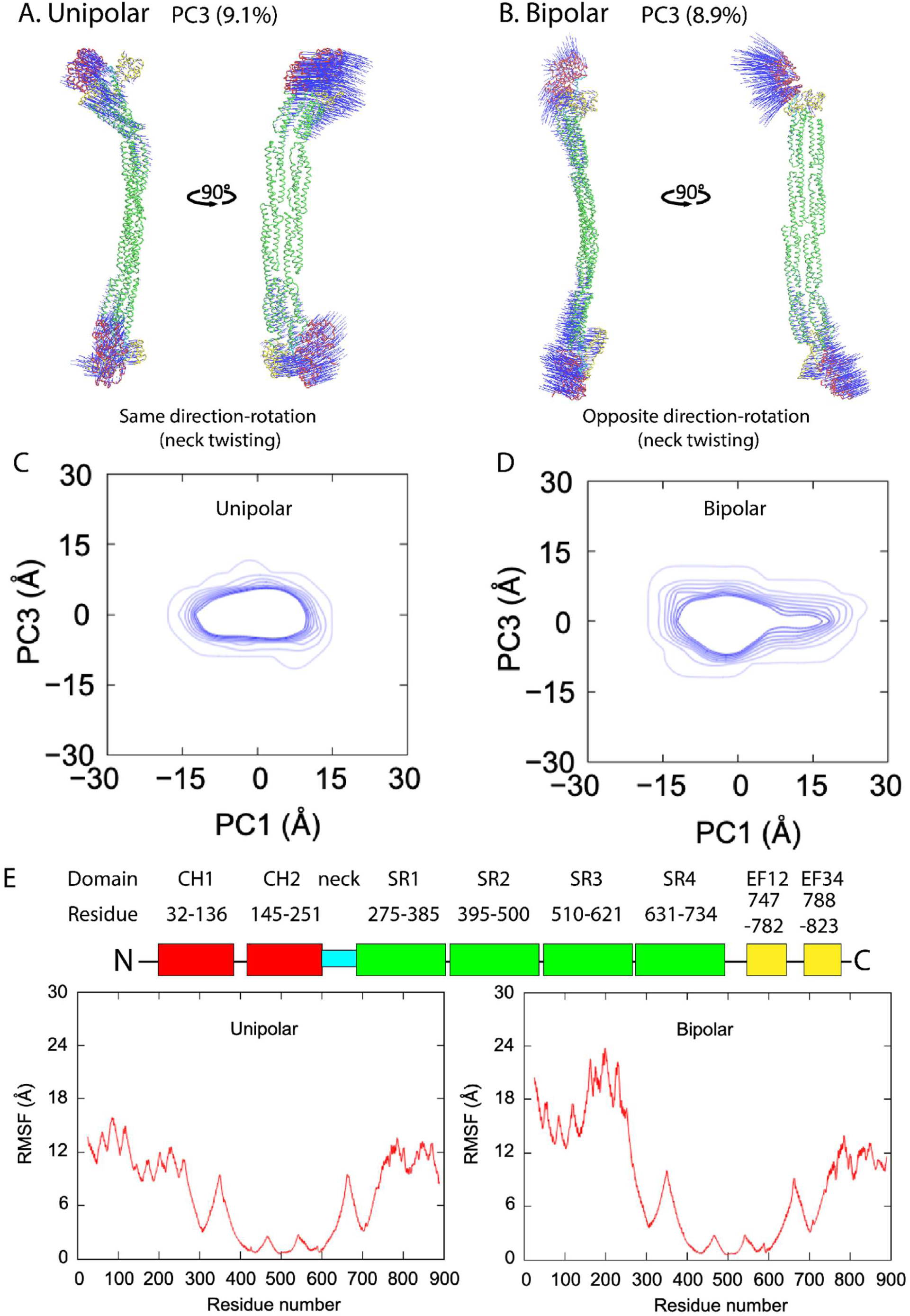
Analysis of the atomic conformational dynamics of alpha-actinin derived from the unipolar and bipolar models using all-atom MD simulations and PCA. (**A, B**) The alpha-actinin structures are illustrated in tube conformations, with the ABD, neck, rod, and EF12-EF34 domains represented in red, cyan, green, and yellow, respectively. The calponin homology 1 and 2 (CH1-CH2) domains in ABDs retained in open conformations when binding to actin filaments. The eigenvalues of principal components 3 (PC3) were used to determine the percentage contribution of each motion. Eigenvector components indicating the direction of motion are shown with blue arrows. The all-atom MD simulations were run for 300 ns (see Methods). (**C, D**) Distributions of conformational ensembles projected onto the common PC1-PC3 space. Data were obtained from conformational libraries generated by MD simulations. (**E**) Root mean square fluctuation (RMSF) values plotted against the residue number. The illustration of the alpha-actinin domains along with their corresponding amino acid residues is shown. From the N-terminus to the C-terminus: CH1, CH2: calponin homology 1 and 2; SR1-SR4: spectrin-like repeat 1-4; and EF12, EF34: EF hand motifs 12, 34.

**Figure S29.**
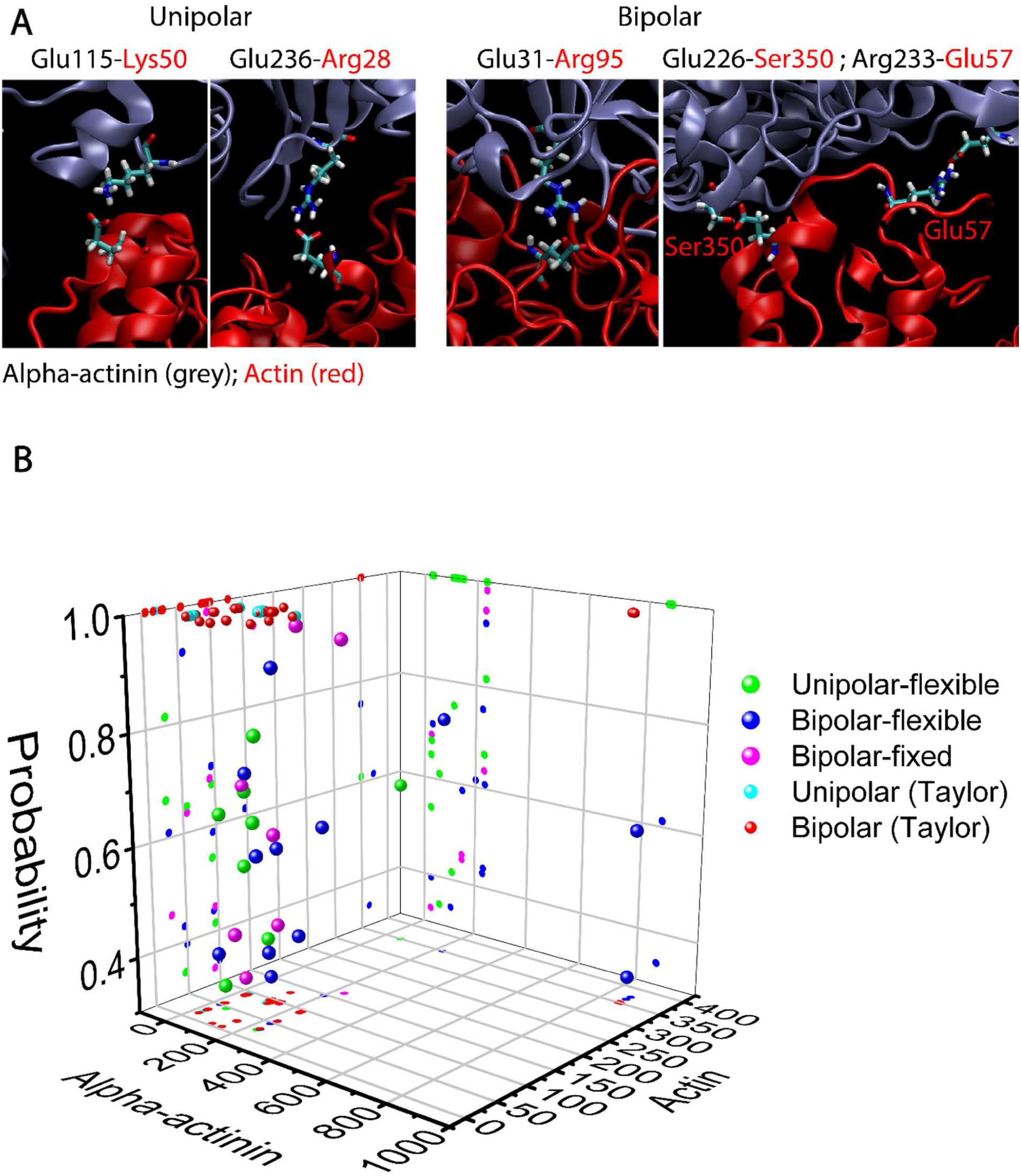
Residue-level interactions between alpha-actinin actin binding domains (ABDs) and actin protomers in the unipolar-flexible and bipolar-flexible crosslinking models. **(A)** Structural representations and residue-contact maps illustrating interactions between alpha-actinin ABDs (red) and actin protomers (grey). **(B)** Probability of residue contacts derived from MD simulations. Two residues were considered to be in contact when the shortest distance in all pairs between atoms in two residues was ≤2 Å. Key alpha-actinin ABD-actin interacting residue pairs were identified based on contact probability densities exceeding 0.5. Because each alpha-actinin molecule contains two ABDs that interact with actin filaments via CH domain, the maximum contact probability density is 2. The results were also compared with that in Liu and Taylor model (Liu et al., 2004).

**Figure S30.**
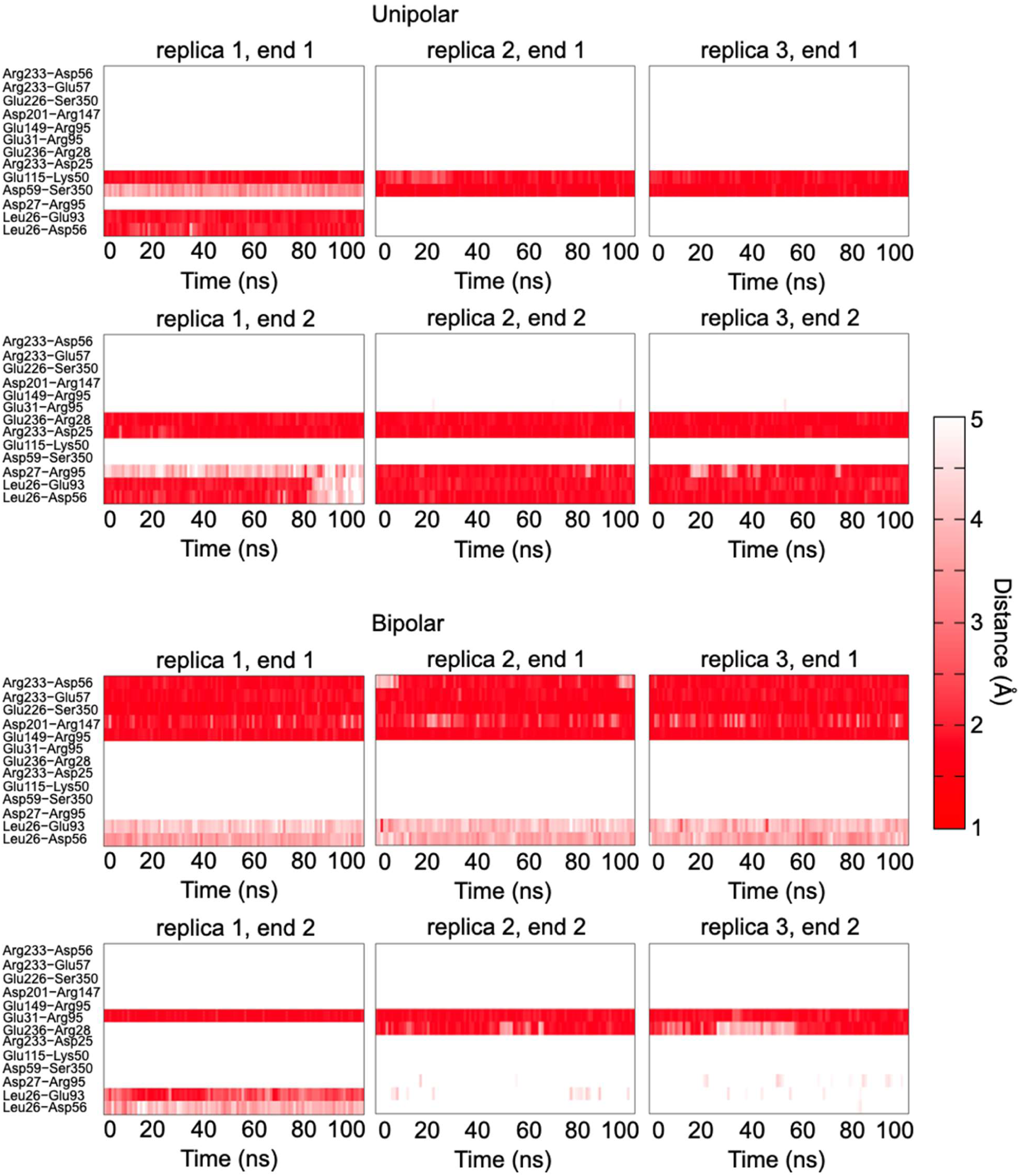
Time evolutions of the minimum distances between actin and alpha-actinin residues with contact probabilities greater than 0.5.

**Figure S31.**
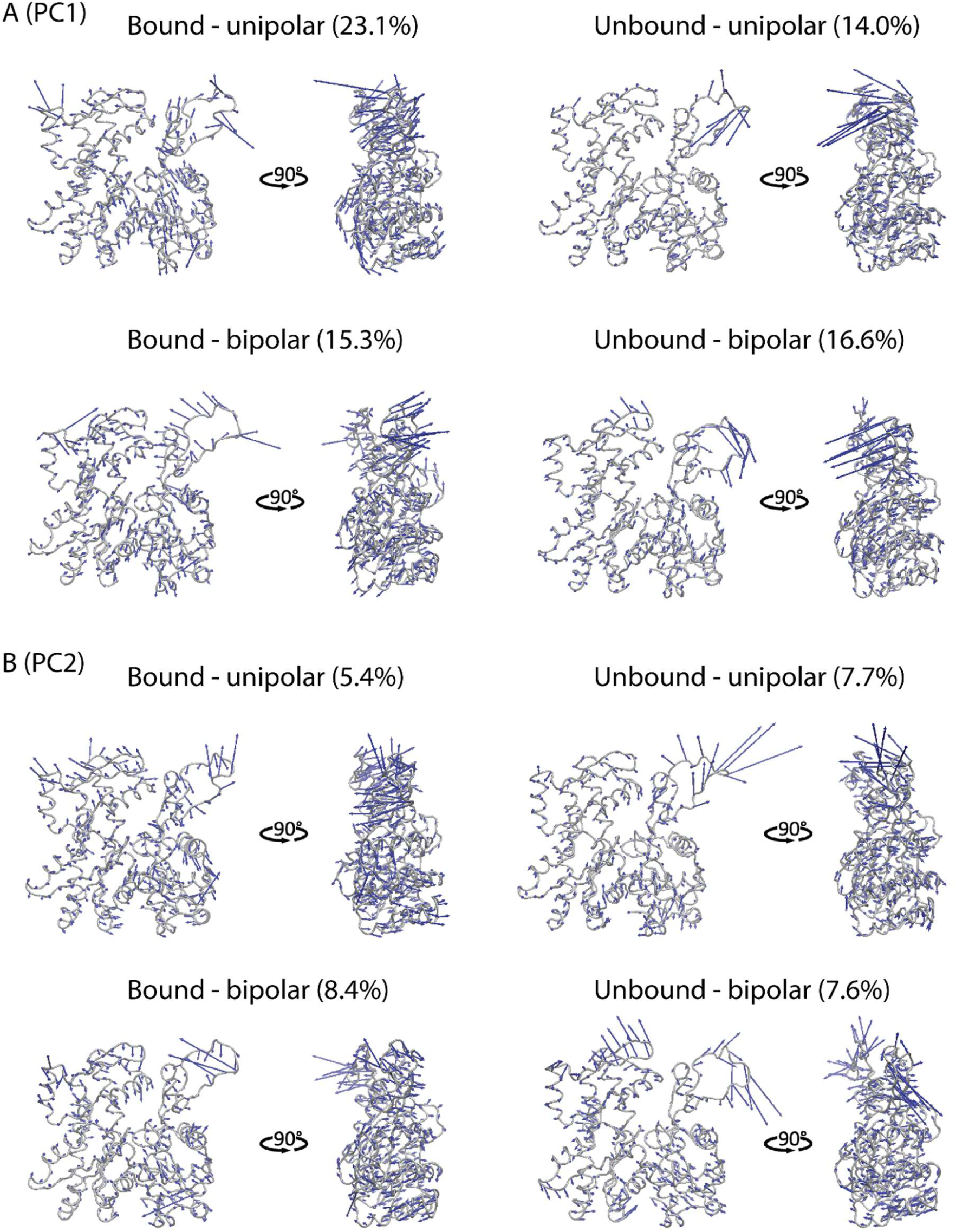
Analysis of atomic conformational dynamics of bound and unbound actin protomers from the unipolar and bipolar filament models using MD simulations and PCA. (**A, B**) Principal components (PC1, PC2) captured flattening motions of bound and unbound actin protomers in the unipolar and bipolar filament models. Actin structures are shown in tube representation with the four subdomains (SD1-SD4) highlighted. The eigenvalues of PC1 indicate the percentage contribution to the positional variance of the dominant motions. Eigenvector components depicting the direction of motion are indicated by blue arrows.

**Figure S32.**
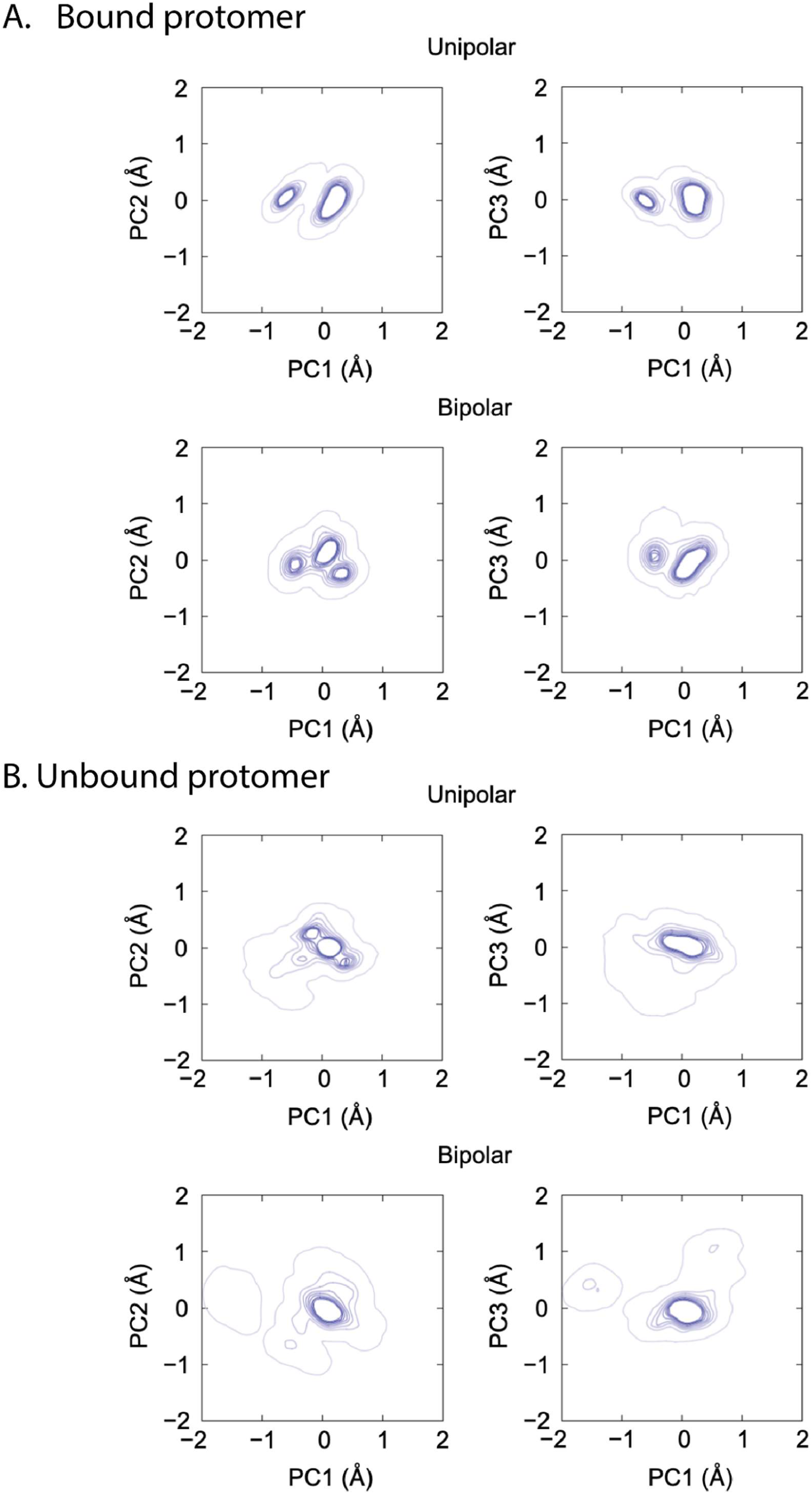
Distributions of conformational ensembles of bound and unbound actin protomers projected onto the common PC1-PC3 space. Data were obtained from conformational libraries generated by MD simulations over 300 ns with three 100 ns replicas.

## Legends for Videos

**Video S1 HS-AFM imaging of the dynamic conformational changes in actin filaments crosslinked by alpha-actinin in the presence of ATP, Ca²⁺, and S1.** Actin filaments were incubated with 50 nM alpha-actinin under conditions containing 1 mM ATP, 0.1 mM CaCl₂, 20mM KCl, and 1 µM S1, but without MgCl_2_ at pH 6.8. The AFM images were processed using a Gaussian filter (σ = 1). The Video plays at 5 frames per second. Scale bar: 25 nm. For annotation, refer to Figure 1 and S3.

**Video S2 HS-AFM imaging of the dynamic conformational changes in actin filaments crosslinked by alpha-actinin in the absence of S1.** Actin filaments were crosslinked with 50 nM alpha-actinin without S1 and observed in F-buffer supplemented with 0.5 mM MgCl₂, 20 mM KCl, and 0.5 mM ATP, but without CaCl_2_ at pH 6.8. The AFM images were recorded at an imaging rate of 500 ms per frame and processed using a Gaussian filter (σ = 1). The Video plays at 5 frames per second. Scale bar: 25 nm. For annotation, refer to Figure S4B.

**Video S3 HS-AFM analysis of helical structures in actin filaments without and with alpha-actinin crosslinking in the presence of cofilin.** Actin filaments were gently immobilized on a lipid membrane and imaged in F-buffer containing 1 mM ATP, 600 nM hs cofilin, and either (A) without or (B) with 400 nM alpha-actinin. Alpha-actinin crosslinking was found to suppress the shortening of half helices and the formation of mature cofilin clusters. AFM images were captured at an imaging rate of 500 ms per frame and processed with a Gaussian filter (σ = 1). Time labels indicate the elapsed time after cofilin addition. The Video plays at 5 frames per second. Scale bar: 25 nm. For annotation, refer to Figure 2 and S7-S9.

**Video S4 HS-AFM analysis of cooperative binding and cluster formation of cofilin on the canonical double-helical actin filaments and alpha-actinin-crosslinked actin filaments.** Experimental conditions were similar to that in Video S3. For annotation, refer to Figure 2 and S7-S9.

**Video S5 HS-AFM imaging was conducted to explore the competitive interactions between cofilin and S1 on actin filaments in the presence of ADP and ATP.** Actin filaments were gently immobilized on a lipid membrane and imaged in F-buffer initially containing 1 mM ADP, 0.01 mM ATP, and 300 nM S1, followed by the addition of 100 nM cofilin. When ATP was gradually or nearly depleted, sTable1 clusters were formed near cofilin clusters on actin filaments. AFM images were acquired at a rate of 500 ms per frame and processed using a Gaussian filter (σ = 1). Time labels indicate the elapsed time after cofilin addition. The Video plays at 5 frames per second, with a scale bar of 25 nm. For annotation, refer to Figure S10.

**Video S6 HS-AFM imaging was performed to observe actin filaments utilizing distinct shorter and normal HHPs for binding and forming the stable cofilin clusters and S1 clusters, respectively.** Actin filaments were gently immobilized on a lipid membrane and imaged in F-buffer initially containing 1 mM ADP, 0.01 mM ATP, and 300 nM S1, followed by the addition of 100 nM cofilin. As ATP levels completely depleted, sTable1 clusters formed in proximity to cofilin clusters on the same actin filaments. AFM images were recorded at a rate of 500 ms per frame and processed using a Gaussian filter (σ = 1). Time labels indicate the elapsed time after cofilin addition. The Video plays at 5 frames per second. Scale bar: 25 nm. For annotation, refer to Figure 3.

**Video S7 Transient binding of alpha-actinin ABD-E235K to actin filaments.**

(**A, B**) HS-AFM Videos showing the transient binding of ABD-E235K to actin filaments in the presence of 1 μM and 2 μM ABD-E235K, respectively. Time labels correspond to frames following the addition of ABD-E235K. Scale bars: 50 nm. For annotation, refer to Figure S11.

**Video S8 HS-AFM visualization of the parallel-helical protofilament state (PPS) and single-helical protofilament (SP) structures of actin filaments induced by ABD-E235K.** Actin filaments were incubated with 2.0 μM ABD-E235K in solution for approximately 30 min before being immobilized on a lipid membrane for HS-AFM imaging. To obtain high-resolution images of the resulting structures, the imaging buffer was exchanged to remove transiently bound ABD-E235K molecules. Time stamps indicate the elapsed time after ABD-E235K addition. Scale bar, 50 nm. For annotation, refer to Figure 4 and S12.

**Video S9 HS-AFM visualization of the binding and cluster formation of cofilin on the PPS-like-helical pitch segments.** Actin filaments were preincubated with 2.0 μM ABD-E235K in solution for approximately 30 min, subsequently immobilized on a lipid membrane, and washed to remove transiently bound ABD-E235K before the addition of 3.5 μM cofilin. Time stamps indicate the elapsed time after cofilin addition. Scale bar: 50 nm. For annotation, refer to Figure S14-S15.

**Video S10 Video S16 ABD-E235K perturbs complete cofilin decoration and shortening of helices of actin filaments.**

**(A)** Before buffer exchange to remove unbound ABD-E235K, cofilin decorated actin filaments only partially, preventing the formation of full cofilin clusters and shortening of HHP. Actin filaments were first immobilized on a lipid membrane, after which 2.0 μM ABD-E235K was added and imaged by HS-AFM for approximately 30 min, followed by the addition of 3.5 μM cofilin. Time labels indicate the elapsed time after cofilin addition. Scale bar, 50 nm.

**(B)** Continuation of the experiment shown in A. Buffer exchange was performed for approximately 5 min to remove unbound ABD-E235K before HS-AFM imaging was resumed. HS-AFM images were acquired at 0.5 s per frame, and the Video is displayed at 5 frames s⁻¹. Scale bar, 50 nm. For annotation, refer to Figure S16-S17.

**Video S11 Binding of alpha-actinin ABD-WT to actin filaments.**

(**A, B, C**) HS-AFM Videos showing the transient binding of ABD-WT to double-helical actin filaments, the parallel-helical protofilament state (PPS), and nano-rings at the final concentration of 2 μM. Actin filaments were gently immobilized on lipid membranes containing 10 % DPTAP, subsequently added into the imaging chamber with 2 μM ABD-WT and HS-AFM imaging. Time labels indicate the time after adding 2 μM ABD-WT. White scale bars: 50 nm. For annotation, refer to Figure 5 and S18.

**Video S12 Stable binding and cluster formation of alpha-actinin ABD-WT on actin filaments.** (**A, B**) HS-AFM Videos showing the stable binding and cluster formation of ABD-WT on actin filaments in the presence of 1 μM. Actin filaments were preincubated with 1.0 μM ABD-WT in solution for 30 min at room temperature, prior to immobilizing on lipid membrane for HS-AFM imaging. Time labels correspond to frames following the HS-AFM imaging. Scale bars: 50 nm.

**Video S13 Molecular dynamics of unipolar-flexible crosslinking model obtained from all-atom MD simulations.** The simulation box remained unchanged. Consequently, slight drifting of the model in x-axis in the box may cause the actin protomers to move in or out of the box during simulation.

**Video S14 Molecular dynamics of bipolar-flexible crosslinking model obtained from all-atom MD simulations.** The simulation box remained unchanged. Consequently, slight drifting of the model in x-axis in the box may cause the actin protomers to move in or out of the box during simulation.

**Video S15 Molecular dynamics of bipolar-fixed crosslinking model obtained from all-atom MD simulations.** In the model, all Cα atoms in alpha-actinin were computationally constrained to prevent vibrations. The simulation box remained unchanged. Consequently, slight drifting of the model in x-axis in the box may cause the actin protomers to move in or out of the box during simulation.

**Video S16 Molecular dynamics of bare actin filament model obtained from all-atom MD simulations over 300 ns with three 100 ns replicas.**

**Video S17** PC1 captured a conserved flattening motion across bare actin, unipolar, and bipolar filaments, indicating that the dominant conformational mode of actin protomers was preserved despite crosslinking. MD simulations were carried out for 300 ns with three 100 ns replicas.

**Video S18** PC1, PC2, PC3 described physical motions alpha-actinin as derived from the unipolar crosslinking model obtained from all-atom MD simulations. MD simulations were carried out for 300 ns with three 100 ns replicas.

**Video S19** PC1, PC2, PC3 captured physical motions alpha-actinin as derived from the bipolar crosslinking model obtained from all-atom MD simulations. MD simulations were carried out for 300 ns with three 100 ns replicas.

**Video S20** PC1 captured a conserved flattening motion of actin protomers bound with alpha-actinin derived from the unipolar and bipolar models. MD simulations were carried out for 300 ns with three 100 ns replicas.

**Video S21** PC1 captured a conserved flattening motion of actin protomers unbound with alpha-actinin derived from the unipolar and bipolar models. MD simulations were carried out for 300 ns with three 100 ns replicas.

## Notes

### Competing Interest Statement

The authors have declared no competing interest.

### Summary of Updates

We would like to update the latest version of our paper.

